# Bacteria-to-human protein networks reveal origins of endogenous DNA damage

**DOI:** 10.1101/354589

**Authors:** Jun Xia, Li-Ya Chiu, Ralf B. Nehring, María Angélica Bravo Núñez, Qian Mei, Mercedes Perez, Yin Zhai, Devon M. Fitzgerald, John P. Pribis, Yumeng Wang, Chenyue W. Hu, Reid T. Powell, Sandra A. LaBonte, Ali Jalali, Meztli L. Matadamas Guzmán, Alfred M. Lentzsch, Adam T. Szafran, Mohan C. Joshi, Megan Richters, Janet L. Gibson, Ryan L. Frisch, P.J. Hastings, David Bates, Christine Queitsch, Susan G. Hilsenbeck, Cristian Coarfa, James C. Hu, Deborah A. Siegele, Kenneth L. Scott, Han Liang, Michael A. Mancini, Christophe Herman, Kyle M. Miller, Susan M. Rosenberg

**Author notes:** Senior author (Christophe Herman, Kyle M. Miller). These authors contributed equally to this work (Jun Xia, Li-Ya Chiu). Graduate School of the Stowers Institute for Medical Research,1000 East 50th Street, Kansas City, MO 64110, USA. Doctorate in Biomedical Science, Universidad Nacional Autónoma de México, México (Meztli L. Matadamas Guzmán). Multidisciplinary Centre for Advance Research and Studies (MCARS), Jamia Millia Islamia, New Delhi 110025, India (Mohan C. Joshi). DuPont Industrial Biosciences, 200 Powder Mill Road. Wilmington, DE 19803. **Correspondence:** **(CH),** **(KMM),** **(SMR)**.

## Abstract

DNA damage provokes mutations and cancer, and results from external carcinogens or endogenous cellular processes. Yet, the intrinsic instigators of DNA damage are poorly understood. Here we identify proteins that promote endogenous DNA damage when overproduced: the DNA-damaging proteins (DDPs). We discover a large network of DDPs in *Escherichia coli* and deconvolute them into six DNA-damage-causing function clusters, demonstrating DDP mechanisms in three: reactive-oxygen increase by transmembrane transporters, chromosome loss by replisome binding, and replication stalling by transcription factors. Their 284 human homologs are over-represented among known cancer drivers, and their expression in tumors predicts heavy mutagenesis and poor prognosis. Half of tested human homologs, when overproduced in human cells, promote DNA damage and mutation, with DNA-damaging mechanisms like those in *E. coli*. Together, our work reveals DDP networks that provoke endogenous DNA damage and may indicate functions of many human known and newly implicated cancer-promoting proteins.

## INTRODUCTION

DNA damage often underlies “spontaneous” mutations (Hastings et al., 1976; Tubbs and Nussenzweig, 2017), which drive cancer, genetic diseases, pathogen drug resistance and evasion of immune responses, and evolution generally. DNA damage can be caused by exogenous agents such as radiation or tobacco smoke, and indeed the vast majority of known carcinogens are DNA-damaging agents, and because of that, mutagens (Chatterjee and Walker, 2017). However, most DNA damage is generated endogenously within cells (Jackson and Loeb, 2001; Tubbs and Nussenzweig, 2017), by intrinsic cellular processes that involve macromolecule components including proteins. Presumably, proteins that promote endogenous DNA damage are required by cells, but cause DNA damage as side effects of their necessary functions and/or when dysregulated. The identities and functions of the proteins that promote endogenous DNA damage in cells in any organism are poorly understood or unknown (Figure 1A). We sought to identify these proteins systematically, and to understand how they might promote endogenous DNA damage, here.

**Figure 1.**
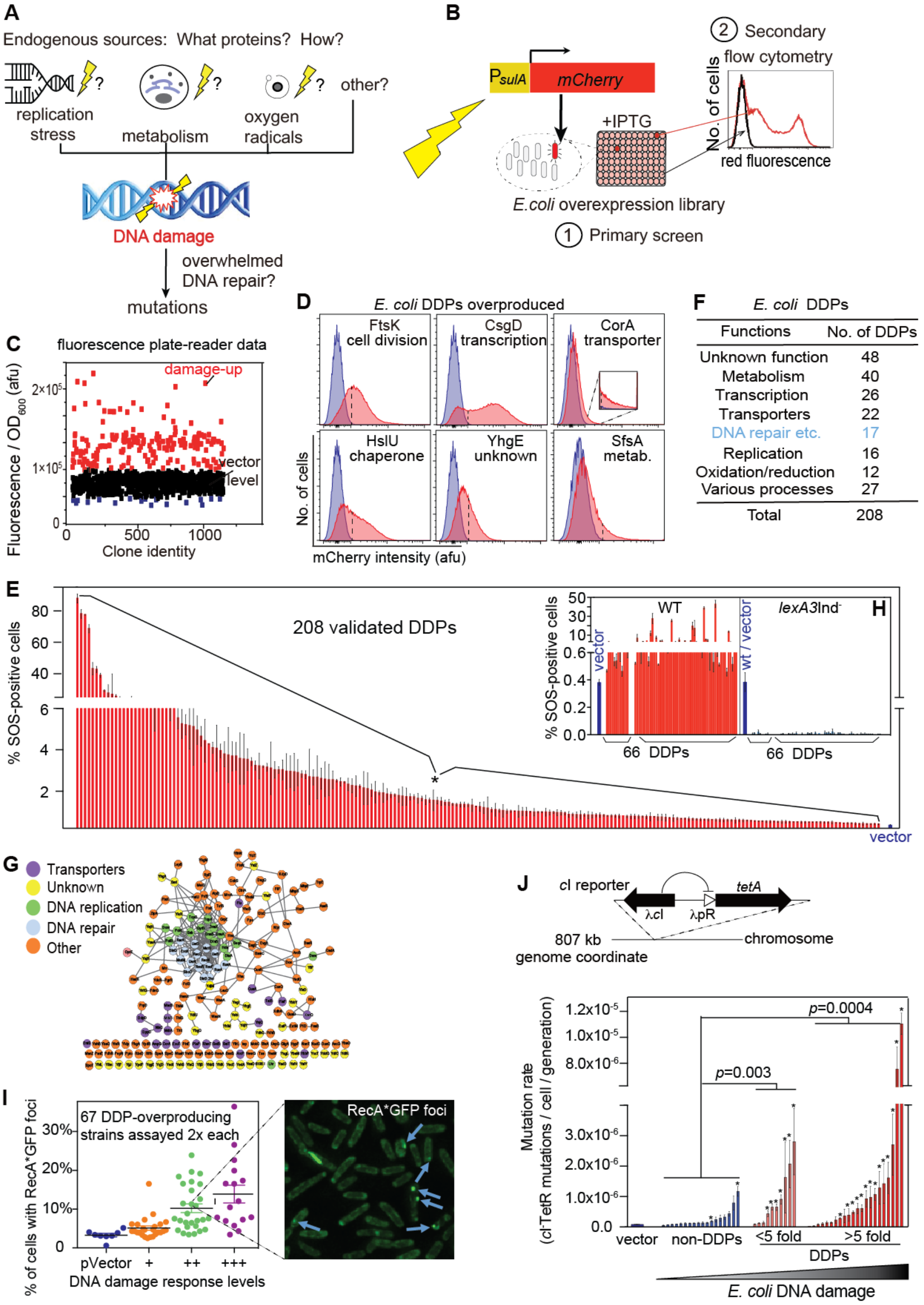
Comprehensive Discovery of *E. coli* DNA-Damaging-Protein Network. (A) We sought comprehensive identification of gain-of-function *E. coli* “DNA-damaging” proteins (DDPs): proteins that provoke DNA damage when overproduced, modeling frequent protein overproduction via various mechanisms common in cancers and bacteria. Although hypotheses for the origins of endogenous DNA damage have been suggested (shown), how these may arise, whether they are common naturally/spontaneously, and what proteins cause them are unclear. (B) Scheme of *E. coli* comprehensive overproduction screen for DDPs. (1) Primary screen: the complete *E. coli* overexpression Mobile library was screened by plate reader for increased fluorescence from an SOS-DNA-damage-response-reporter gene, P_*sulA*_*mCherry* (Nehring et al., 2016). (2) Secondary screen: potential positive clones from the primary screen were validated, and false-positives eliminated, by sensitive flow-cytometric assay, which reports fluorescence per cell at the single-cell level. (C) Representative results of primary plate-reader screen: afu, arbitrary fluorescence units (SOS activity), per OD_600_ unit, indicating biomass. Red, potential “damage-up” DDP hits with fold change >30%. (D) Representative flow-cytometric validation of SOS-positive (DDP) overexpression clones from plate-reader screens. Dashed line, flow cytometry “gate” above which cells are scored as SOS-positive (STAR Methods). Validated DDP clones have significantly higher frequencies of SOS-positive cells than the vector control. Blue, vector control; red, DDP-overproduction clones. (E) Quantification of increased DNA damage measured as % SOS-positive cells in the 208 validated *E. coli* DDP clones (identities with data, Table S1). (F) *E. coli* DDP network summary; proteins of many different functions are DDPs. Functions of the 208 *E. coli* DDPs (Table S1). Few (8%) are known DNA-repair proteins (blue, Table S1). (G) Protein-protein associations of the *E. coli* over-production DDP network, CytoScape software generated from STRING 10.0 database. Other, defined Table S1; specific protein associations, Figure S2 (discussed, text). (H) LexA-dependence of increased fluorescence of representative DDPs shows that fluorescence results from activation of the SOS DNA-damage response. (I) Correlation of DDP (SOS-positive) phenotype with RecA*GFP foci, indicating persistent single-stranded (ss)DNA. A representative sample of 67 DDPs overproduced showed 32 (48%) with significantly more RecA*GFP foci than the vector control (*p* < 0.05, unpaired two-tail *t*-test), a less sensitive assay than the stringent flow-cytometric assay for SOS-positive cells. RecA*GFP foci are correlated positively with the SOS-response assay: r = 0.7, *p* = 1.3×10^−10^, Pearson’s correlation. **Left**, DDP clones assayed. Each dot, one DDP clone, assayed twice (mean). Blue, negative control; orange, low SOS activity (DNA damage); green moderate DNA damage; purple, high DNA damage, strain-by-strain data, Table S1. **Right**, representative foci. (J) Mutation-rate increase in representative DDP-overproducing clones. Above, *c*I mutation assay design (Gutierrez et al., 2013). Loss-of-function mutations in a chromosomal *c*I gene, encoding phage lambda transcriptional repressor, allow transcription of *tetA*, which confers tetracycline resistance (TetR) (STAR Methods). Below, increased mutation rates are associated with increased DNA-damage levels (SOS induction, flow-cytometric assay) in *E. coli* DDP-overproducing clones. Each bar, the mean mutation rate (± SEM) of each strain, 3 experiments (fluctuation tests, STAR Methods). The DDPs overproduced (Supplemental Discussion 3) represent various classes (Table S1). Table S1 for mutation rates. *P*-values, one-way Fisher’s exact test of the number of clones with mutation rate significantly higher than the vector-only control (unpaired 2-tailed Student’s *t*-test).

One way to identify the proteins that *promote* endogenous DNA damage is by overproduction, which is a natural event that occurs frequently by copy-number alteration and other routes, and is a major source of cancer-driving functions (Zack et al., 2013). Given the conservation of DNA biology across the tree of life (Aravind et al., 1999; Makarova and Koonin, 2013), identification of proteins that promote spontaneous endogenous DNA damage carried out in any organism could potentially inform strategies for prevention, diagnosis, and treatment of disease, including therapeutic inhibition of cancer development, aging, and evolution of pathogens (Fitzgerald et al., 2017).

Some proteins that *prevent* or *reduce* endogenous DNA damage levels in cells have been identified by loss-of-function mutations/knock-downs that increase DNA damage (Alvaro et al., 2007; Lovejoy et al., 2009; Paulsen et al., 2009). DNA-repair proteins are among this category because they *reduce* levels of endogenous DNA damage. Similarly, proteins that *prevent/reduce* endogenous DNA damage, along with other kinds of proteins, will be among genes the loss-of-function of which promotes mutagenesis or genome instability (Putnam et al., 2016; Yuen et al., 2007). By contrast, no unbiased screen has been reported for proteins that actively *promote* endogenous DNA damage in cells. Though a limited nucleus-specific screen identified some (Lovejoy et al., 2009), the range of functions and numbers of proteins, processes, and mechanisms that cause endogenous DNA damage are unknown.

Here we report the comprehensive discovery in *Escherichia coli* of a large, diverse network of proteins that promote endogenous DNA damage when overproduced: the “DNA-damaging” proteins (DDPs). We show that their upregulation promotes mutagenesis, and use massive function-based assays to identify kinds, causes, and consequences of intrinsic DNA damage provoked by overproduction of the 208 *E. coli* DDPs. We found that they group into six discreet function clusters, and determine molecular mechanisms of DNA-damage generation from three of these. We identify their human homologs, also a large network, and find that they are overrepresented among known cancer instigators, and that their expression in human cancers predicts poor outcomes and high mutation loads. We show that overproduction of human homologs in human cells also promotes endogenous DNA damage and mutation. We determine the mechanisms of DNA-damage instigation for two cancer-associated human DDPs, both of which mimic bacterial mechanisms and suggest unexpected roles in cancer. The identities and functions of the proteins in bacterial and human DDP networks provide an important general model for illuminating mechanisms of genesis of endogenous DNA damage, and may inform cancer-promoting function discovery of many known and newly discovered cancer-driving proteins.

## RESULTS

### Large Diverse Protein Network Promotes DNA Damage

We screened an inducible overproduction library of all >4000 *E. coli* proteins in two steps to identify clones with increased endogenous DNA-damage levels (STAR Methods, Figure 1B). For both the primary and secondary screens, we measured fluorescence of cells that carry a fluorescence-reporter gene driven by an SOS DNA-damage-response-activated promoter at a non-genic chromosomal site (Nehring et al., 2016) (Figure 1B). The promoter fusion reports DNA-damage-response induction, and not spurious promoter firing (Pennington and Rosenberg, 2007) (Figures S1A and S1B). First, in the primary screen, we used a fluorescence plate-reader, which is high-throughput but low-resolution, to identify potential clones with increased DNA damage and so increased fluorescence (Figures 1B and 1C). This identified potential positive candidates (414 proteins). Second, we eliminated false positives from the primary plate-reader screen using a sensitive flow-cytometry secondary screen in the same strains (Figures 1B and 1D). Flow cytometry, though low-throughput, is highly sensitive and reports DNA damage at the single-cell level (Pennington and Rosenberg, 2007) (STAR Methods, Figure 1D). The stringent, high-sensitivity flow-cytometry secondary screen validated 208 of the proteins identified in the primary screen as genuine DDPs that cause increased DNA damage when overproduced (Figures 1D-G; Table S1).

Further, we tested a representative sample of 66 of the DDP-overproduction clones (Table S1) for whether their increased fluorescence requires the SOS-response-activator protein RecA and a functioning (inducible) LexA SOS repressor, as expected if the fluorescence results from activation of the SOS DNA-damage response (Pennington and Rosenberg, 2007). All 66 showed RecA- and LexA-dependent high fluorescence, demonstrating the presence of DNA damage and a genuine SOS response (Figures 1H and S1C; Table S1). We also ruled out the possibility that DDP overproduction might increase mCherry protein fluorescence itself, independently of DNA damage, by showing that a separate representative sample of 40 of the 208 DDPs did not cause increased fluorescence from the same mCherry reporter gene under the control of a non-SOS promoter (Figure S1D; Table S1).

DNA-damage-promotion by overproduction of the 208 DDPs is observed additionally in three ways. First, 95% of the 208 validated *E. coli* DDPs displayed an independent DNA-damage-related phenotype in at least one of nine assays that are either less sensitive or more DNA-damage-type specific than the SOS-flow cytometric assay in the secondary screen. For example, we tested a representative sample of 67 of the DDP-overproduction clones for the presence of damaged, single-stranded DNA using a less sensitive assay for microscopically visible foci of a fluorescent DNA-damage-sensor protein RecA*GFP (Renzette et al., 2005), which is less sensitive because it uses a partially functional RecA protein (Renzette et al., 2005). The data nevertheless show a significant association of RecA foci with SOS-positive (DDP) clones in that 32 of the 67 tested showed increased foci (r = 0.7, *p* = 1.3×10^−10^, Pearson’s correlation) (Figure 1I; Table S1 for clone-by-clone results). Also, later in this paper, we explored the kinds and causes of endogenous DNA damage promoted by the 208 *E. coli* DDPs, using seven assays for specific kinds of DNA damage—all more DNA-damage-type-specific than the SOS-response assay used here, and we additionally assayed DDPs for mutagenesis, described below (summarized Table S1; Figure 1J). All but 12 of the 208 clones were positive in either the RecA*GFP-focus assay, and/or one of the 7 assays described below, or mutagenesis (summarized Table S1). This equates to 95% of the 208 proteins showing DNA-damage-related phenotypes in an independent assay. Finally, all of the twelve not validated in a non-SOS-based assay (Table S1) were shown to increase fluorescence SOS-response dependently, showing only RecA-LexA-dependent fluorescence increase (Figures 1H and S1C; Table S1) and not general fluorescence increase (Figure S1D; Table S1). Collectively, these data demonstrate independently that all 208 DDPs promote DNA damage when overproduced.

The 208 proteins span many different classes that function in diverse cellular and metabolic processes (Figure 1F; Table S1), and only 8% encode known DNA-repair proteins (blue font, Figure 1F; Table S1). Although DNA-repair proteins *reduce* DNA damage when expressed normally, their overproduction, here, increased DNA damage, which might occur by perturbing undamaged DNA, by titrating repair partner proteins away from DNA damage and/or inhibiting DNA repair. We call all of these proteins DNA-damaging proteins (DDPs).

The DDPs constitute a network both functionally and by protein-protein associations. Functionally, they cause the same phenotype—increased DNA damage on overproduction—but their reported protein functions are remarkably diverse (Figure 1F; Table S1). We used STRING—a database that contains known and predicted associations between protein pairs—to examine whether these diverse proteins had any other known or predicted connections indicative of a network. STRING measures protein-protein associations or interactions of many kinds including indirect associations (e.g., co-occurrence in different organisms’ genomes, or in papers in the literature) and direct protein-protein interactions, among others (STAR Methods). Using STRING with an interaction score cut-off of ≥0.6 (medium-to-high confidence, STAR Methods), we found that the 208 DDPs form a significant network via protein-protein interaction data (Figure 1G, specific interactions, Figure S2A) with more interactions than random sets of 208 *E. coli* proteins (*p* = 2.0 × 10^−31^, hypergeometric test, Supplemental Discussion 1). When known DNA-repair proteins are removed, the STRING network is still significant compared with random sets of the same number of *E. coli* proteins (*p* = 9 × 10^−7^, hypergeometric test), indicating that this association network is not solely via DNA-repair-protein associations. When both DNA-repair and DNA-replication proteins are removed—both known to interact directly with DNA—the STRING network is no longer significant compared with random sets of the same number of *E. coli* proteins (*p* = 0.08, hypergeometric test). These data suggest that, as might be expected from the highly diverse protein functions (Figure 1F), these proteins work in many different cellular processes that may share only their various effects on DNA damage, seen by significant association as a network only when DNA-repair and replication proteins are included, which we identify as the hubs of the DDP STRING interaction network (Figure 1G).

The DDP network is estimated to be larger than the 208 proteins identified (Supplemental Discussion 2, Figure S1E). Although the premier overproduction (Mobile) library was used (Saka et al., 2005), its composition of some native genes and some genes encoding five additional amino acids is indicated by our data to have prevented detection of some additional DDPs (Supplemental Discussion 2).

### Endogenous DNA Damage Increases Mutations

We tested the hypothesis that triggering endogenous DNA damage would increase mutation rates (Figure 1A) using DDPs: a useful test because they were discovered based on DNA damage, rather than genome instability/mutation rate. Mutation rates of a sample of 32 representative *E. coli* DDP clones were assayed by a modified forward-mutation fluctuation-test assay (Figure 1J, STAR Methods). We chose 10 DDP clones from the low-damage group (<5-fold increase in endogenous DNA-damage levels) and 22 from the high-damage group with a >5-fold increase in endogenous DNA-damage levels (Supplemental Discussion 3; Table S1). The data in Figure 1J show that increased endogenous DNA-damage levels are associated significantly with elevated mutation rates. We confirmed that the mutagenesis assay reported genuine loss-of-function mutations in the *c*I mutation-reporter gene (Figure S1F), rather than gene-regulatory or epigenetic changes, by sequencing mutations in a sample of independent isolated mutants of 10 representative DDP clones (Figure S1F). The sequence analyses of selected DDP clones revealed additionally that high DNA damage led to various kinds of mutations including base substitutions, indels, transposition events, and gross chromosomal rearrangements (GCRs, including large deletions, Figure S1F, right). These mutations mimic the increased small mutations and GCRs seen in various cancers (Stratton, 2011). Although the SOS response induces mutations by upregulation of low-fidelity DNA polymerases (Pols) V and IV (Kobayashi et al., 2002; Maor-Shoshani et al., 2000; Wagner and Nohmi, 2000), some of the mutations (Figure S1F, blue font) differ from common Pol V and Pol IV errors (Figure S1F, red font). The data suggest that the type of DNA damage, and not merely induction of the SOS response, may influence the kinds and rates of mutations made (Figures 1J and S1F, Table S1). The data show that overproduction of many functionally diverse *E. coli* proteins (Figure 1F; Table S1) causes increased DNA-damage loads (Figures 1C-E; Table S1), and genome instability with mutations of essentially all kinds (Figures 1J and S1F; Table S1).

### Human Homologs of Bacterial DDPs a Network Associated with Cancers

As an unbiased quantitative way to find human DDPs, we identified 284 human homologs of the *E. coli* DDPs via BLASTp and deltaBLAST searches (STAR Methods, Figure 2A; Table S2). “Homologs” are defined here as proteins with amino-acid similarity that may result from possible evolutionary relatedness (STAR Methods). The 284 human homologs are used here as candidate human DDPs (hDDPs), and are homologs of 68 of the *E. coli* DDPs (shown, Table S2). The remaining *E. coli* DDPs are mostly analogs of human proteins, which function similarly but are not homologous (Serres et al., 2001), and many are of unknown function. The hDDP candidate proteins also constitute a protein-protein interaction network (Figure 2B, specific interactions Figure S2B), with significantly more interactions than sets of 284 random human proteins (*p* = 1.2 × 10^−327^, hypergeometric test), or random human homologs of *E. coli* proteins, which also differ from random human proteins, but less so (*p* = 1.8 × 10^−49^, hypergeometric test, discussed Supplemental Discussion 1). Only 5.6% of the human homologs are known DNA-repair proteins (blue font, Figure 2A), again indicating a different class of candidate genome-integrity-affecting proteins. Like the *E. coli* network (Figure 1G), DNA-repair and -replication proteins are central hubs (Figure 2B).

**Figure 2.**
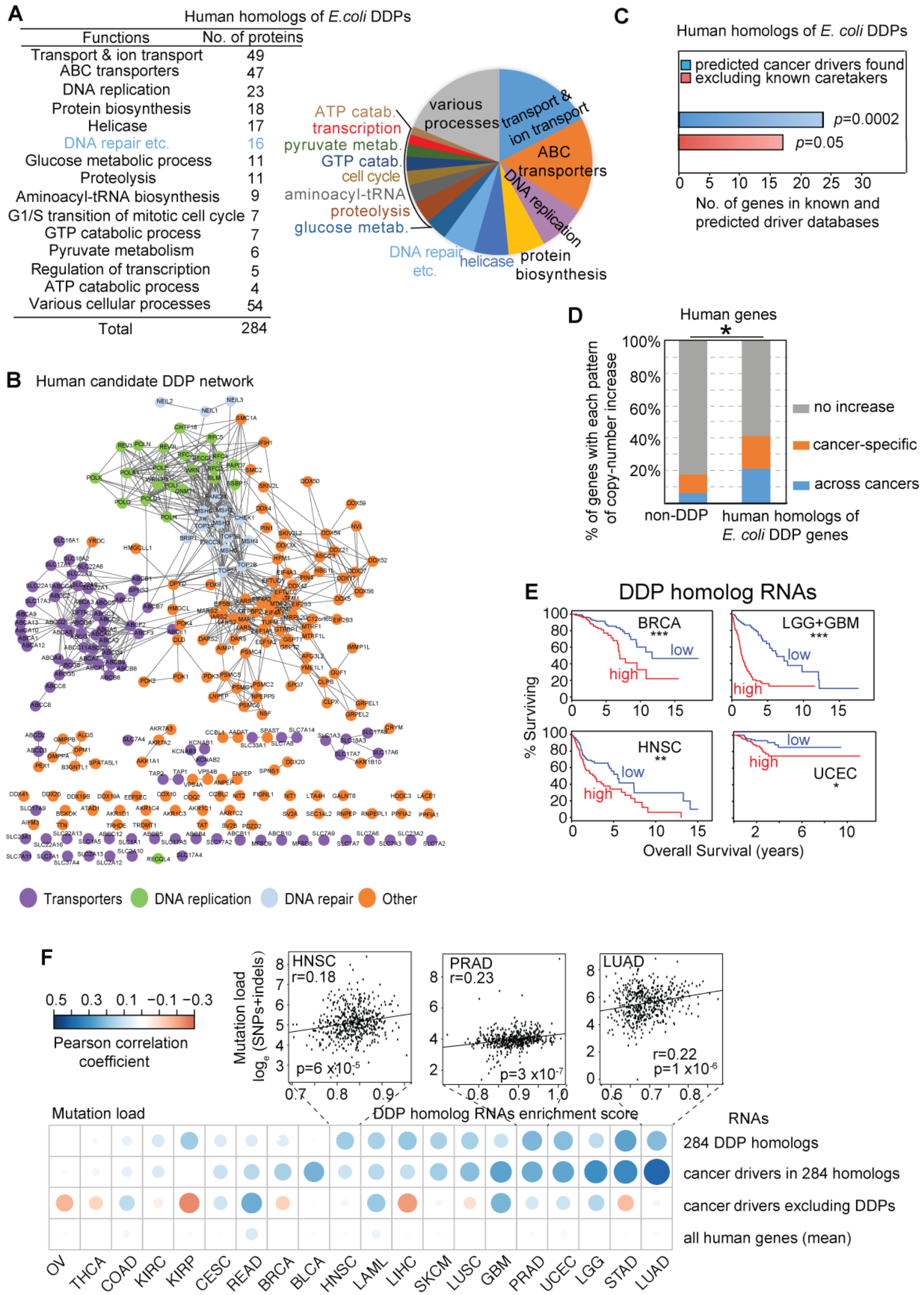
Human Homologs of *E. coli* DDPs a Network Associated with Cancers. (A) Summary of 284 human proteins identified as homologs of *E. coli* DDPs (STAR Methods). Only 5.6% are known DNA-repair genes (blue, Table S2). (B) Protein-protein association network of human homologs of *E. coli* DDPs (Figure S2 for specific interactions). Network displayed per Figure 1G. Other, defined Table S2. (C) Human homologs of *E. coli* DDPs (hDDP candidates) are significantly overrepresented among known (Forbes et al., 2015) and predicted (D’Antonio and Ciccarelli, 2013) cancer drivers (blue bar). After subtracting known DNA-repair (“caretaker”) genes, the remaining hDDP candidate genes are still enriched for known and predicted cancer drivers (red bar). (D) hDDP candidate genes are enriched among genes with cancer-associated copy-number increases, indicating selection for overexpression in cancers. Pan-Cancer copy-number-increase analysis (GISTIC threshold copy-number gain ≥ 1) of the 284 hDDP candidates in 26 cancer types (per Figure S3). Blue, human genes with increased copy numbers across cancers (*p* < 0.05, FDR < 0.10, Wilcoxon test); orange, genes with cancer-specific copy-number increase; grey, not particularly cancer associated. Complete data for the network Table S4. (E) Decreased cancer survival is associated with high DDP-homolog RNA levels in cancers [our analyses of data from TCGA (Gao et al., 2013), STAR Methods]. BRCA, breast invasive carcinoma; LGG+GBM, gliomas (low-grade glioma + glioblastoma multiforme); HNSC, head and neck squamous cell carcinoma; UCEC, uterine corpus endometrial carcinoma. *, **, ***, survival of the cancers with high and low levels of the 284 RNAs differ at*p* ≤ 0.05; ≤ 0.01, and ≤ 0.001 respectively, log-rank test. (F) High RNA levels of the 284 hDDP candidates are associated with tumor mutation burden in data from TCGA (Gao et al., 2013). Each dot represents a Pearson correlation coefficient between the RNA-enrichment score, relative to total RNAs, of a gene set and mutation burden in the tumor. The average correlation strength of 284 hDDP-candidate RNA levels with mutation loads across 20 TCGA cancers was in the top 0.5% of correlations for randomly selected groups of genes across all human genes. Blown-up: correlation of increased mutation loads (y axis) with increased hDDP-candidate RNA enrichment scores (x axis, STAR Methods) in three represented cancers. Cancer-types: OV ovarian serous cystadenocarcinoma; THCA thyroid carcinoma; COAD colon adenocarcinoma; KIRC kidney renal clear cell carcinoma; KIRP kidney renal papillary cell carcinoma; CESC cervical squamous cell carcinoma and endocervical adenocarcinoma; READ rectum adenocarcinoma; BLCA bladder urothelial carcinoma; LAML acute myeloid leukemia; LIHC liver hepatocellular carcinoma; SKCM skin cutaneous melanoma; LUSC lung squamous cell carcinoma; PRAD prostate adenocarcinoma; STAD stomach adenocarcinoma; LUAD lung adenocarcinoma.

We tested whether the human homologs of *E. coli* DDPs are both relevant to human cancers, and behave as a network, by examining their associations with various kinds of data from human cancers. We observe strong associations of the 284-protein network in cancer data of several kinds. First, the human homologs (Figure 2A; Table S2) of *E. coli* DDPs are significantly overrepresented among known (Forbes et al., 2015) and predicted (D’Antonio and Ciccarelli, 2013) cancer drivers in a curated consensus in the Sanger Institute’s Catalogue of Somatic Mutations In Cancer (COSMIC) (Forbes et al., 2015) and the database of D’Antonio and Ciccarelli (D’Antonio and Ciccarelli, 2013) (*p* = 0.0002, Fisher’s exact test; Figure 2C; Table S3), which contain gain- and loss-of-function drivers. Human homologs of random *E. coli* proteins are not overrepresented (*p* = 0.48, Fisher’s exact test). Thus, the cancer association is specific to DDP homologs, not conserved proteins generally. The human homologs remain overrepresented among known and predicted drivers when homologs of *E. coli* DNA-repair proteins and other known human DNA-repair proteins (Figure 2A) are excluded (*p* = 0.05, Figure 2C). No other comprehensive overexpression screen for DNA damage has been reported; however, we analyzed cancer association in published data from a limited, selected-candidate overexpression screen in human cells (Lovejoy et al., 2009), and found that these also show cancer association (Supplemental Discussion 4). These overlap with our 284 hDDP candidate network by only one protein—FIGNL—indicating that many new candidates were revealed by the *E. coli* screen.

Additionally, we found that candidate human DDP genes show increased copy numbers in 26 cancer types in the cohort of patients in The Cancer Genome Atlas (Gao et al., 2013) (TCGA) (Figures 2D and S3A-C; Table S4). About 40% of the 284 human homolog genes have increased copy numbers in cancers (GISTIC threshold copy-number gain ≥ 1), either cancer-specifically or across cancers, compared with fewer than 20% of non-DDP genes amplified in those cancers (Figure 2D). The fractions of patients with increased copy numbers of each of the genes encoding the 284 human homologs of *E. coli* DDPs are shown for all 284 in Table S4 (examples, Figure S3A-C). The human homologs are enriched as copy-number increases in cancers compared with non-DDP human genes (Figure 2D, *p* = 0.04, one-way Fisher’s exact test), suggesting that their overexpression is associated with cancers.

We next examined the outcomes for cancer survival relative to mRNA levels of the 284 candidate human DDPs using cancer-patient and RNA data in TCGA (Gao et al., 2013). We found that in at least four cancer types, increased levels of the 284 RNAs, relative to the total RNAs, is associated with decreased overall survival (Figure 2E). This association results not just from the known (Forbes et al., 2015) and predicted (D’Antonio and Ciccarelli, 2013) cancer driver genes in the network, but is seen also in the network genes not known previously to drive cancers (Figure S3D-F). These data indicate an association of candidate hDDP gene overexpression with poor survival in these cancers, and further highlight the network properties/predictive power of the 284 DDP-homolog protein/gene set. Moreover, increase of the 284 human DDP-homolog RNAs, relative to all RNAs, is also associated with total genomic mutation burden in cancers (relative to the patient normal tissue) in at least 12 cancer types in TCGA (Gao et al., 2013) (Figure 2F). Even stronger association is seen for the subset of the 284 homologs that are known (Forbes et al., 2015) or predicted (D’Antonio and Ciccarelli, 2013) cancer-driving genes (Figure 2F). These data support the possibility that overexpression in the candidate hDDP network is associated with mutagenesis in human cancers.

### Human Homologs Promote DNA Damage and Mutation

We validated a sample of human candidate DDPs as genuine DNA-damage instigators in human cells (Figure 3). We tested our hypothesis that overproduction of these proteins, which can result from gene amplification, can promote DNA damage relevant to cancers by testing a sample in which about half the homologs were known to be amplified in cancers in TCGA (Gao et al., 2013) and the other half were not (Supplemental Discussion 5). We were also limited by availability in human cDNA-clone collections (Yang et al., 2011) (Supplemental Discussion 5). Because many genes in those collections are not full length (Table S5), we cloned several de novo (STAR Methods) to create 70 full-length sequence-verified overexpression GFP-fusion genes encoding human homologs of *E. coli* DDPs, 3 human homologs of *E. coli* damage-down proteins, as possible negative controls (Table S5), and 20 control random non-DDPs (Table S5, ~half of which are random human homologs of *E. coli* proteins). Using transient transfection of these human overexpression clones, we performed three flow-cytometric assays, which we developed (Figure 3A), to screen for increased DNA damage at the single-cell level in human cells that produce the candidate hDDPs, shown by GFP (Supplemental Discussion 6 for the infeasibility of stable clones). We screened for—(i) increased γH2AX levels, a marker for DNA double-strand breaks (DSBs) (Kinner et al., 2008); (ii) increased γH2AX in a sensitized screen in cells treated with a nonhomologous-break-repair (DNA-PK) inhibitor; and (iii) increased levels of the DNA-damage marker protein phospho-p53, which indicates activation of the DNA-damage response (Sakaguchi et al., 1998). The data show that the human homologs are enriched for genuine DDPs that increase DNA damage upon overproduction (Figure 3B). Among the 73 human homologs, we found that 45% (33 of the 73) showed increased DNA damage (Figure 3B). This is highly significant (*p* < 0.0001 one-way Fisher’s exact test) compared with the 20 random human proteins (Figures S3G-I; Table S6). Of the 33 validated hDDPs, only one (FIGNL1) was known previously to increase DNA damage upon overproduction (Lovejoy et al., 2009). Thus, we identified and validated 33 genuine human DDPs.

**Figure 3.**
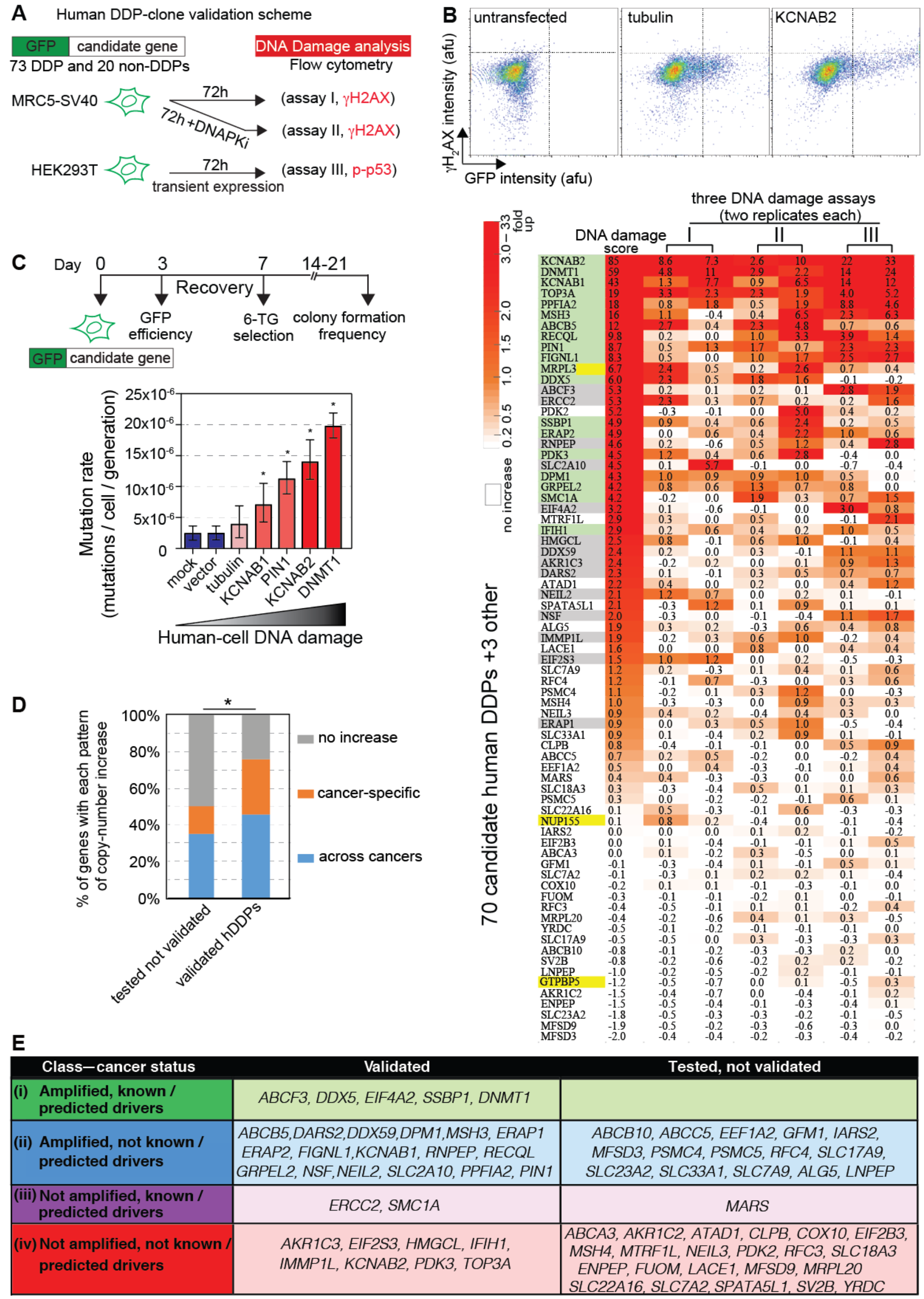
Overproduced Human Homologs Promote DNA Damage in Human Cells. (A) Scheme for validating hDDPs. 70 full-length sequence-verified human homolog candidate-hDDP-GFP N-terminal fusions (and 3 damage-down-, plus 20 non-DDP-GFP fusion controls) were transiently overproduced in MRC5-SV40 or HEK293T cell lines and green cells screened for high DNA damage by flow cytometry. (B) DNA-damage assays with candidate hDDPs identify 33 validated hDDPs. **Upper:** representative flow cytometric DNA-damage assay data (STAR Methods, Figure S3G-I; Table S6). **Lower:** heatmap of the flow-cytometric data normalized to GFP-tubulin. Data from each of the three DNA-damage assays, with 2 replicates each, ranked by a cumulative DNA-damage score that sums the fold changes of each DNA-damage assay for each candidate protein. Highlighting colors: green, significantly damage-up in ≥ 2 assays (two-tailed unpaired *t*-test with FDR correction); gray, significantly damage-up in one assay; yellow, homologs of *E. coli* damage-down proteins; white, not damage-up. 45% validated, significantly more than among 20 random human genes (*p* <0.0001, two-tailed unpaired *t*-test with FDR correction, Figure S3G-I), indicating that homologs of *E. coli* DDPs are enriched for hDDPs. (C) Increased mutation rates in human cells overproducing validated hDDPs, assayed by humancell *HPRT* forward-mutation assays in fluctuation tests (STAR Methods). **Upper**: *HPRT* assay scheme. *HPRT* loss-of-function mutants are selected as 6-thioguaine (6-TG)-resistant clones. **Lower**: mutation rates of selected hDDP overproducers shown with their DNA-damage levels; error bars, 95% confidence intervals. (D) Validated hDDP genes are enriched among genes with cancer-associated copy-number increases compared with the candidates that were tested but not validated (*p* = 0.02, one-way Fisher’s exact test). (E) New and known potential cancer-promoters predicted among 33 validated hDDPs. The 33 validated hDDP genes comprise genes that are—(i) both amplified in TCGA cancers and known (Forbes et al., 2015) or predicted (D’Antonio and Ciccarelli, 2013) cancer drivers (16%); (ii) amplified in TCGA cancers and were not known or predicted cancer-driving genes (53%); (iii) known or predicted cancer drivers that are not found to be amplified in cancers (6%); and (iv) not found to be amplified in cancers in TCGA, and not known or predicted cancer drivers (25%). The data suggest potential overexpression cancer-promoting roles for the genes in all classes.

We note, however, that the human-cell DNA-damage assays used here favor detection of DSBs, not all DNA-damage types comprehensively. Thus, many more of the human homologs may be DNA-damage instigating for other kinds of DNA damage than is estimated here.

As in *E. coli*, we found that overproduction of validated hDDPs increased mutation rates in human cells. Using a forward-mutation fluctuation-test assay for hypoxanthine-guanine phosphoribosyl transferase (HPRT) deficiency (STAR Methods), we found increased mutation rates for 4 out of 4 overproduced high-DNA-damage hDDP clones compared with cell-only, vector-only, and GFP-tubulin-overproducing negative controls (Figure 3C and Supplemental Discussion 8). Thus, increased mutation rates result from overproduction of validated hDDPs in human cells. These results support the hypothesis that hDDPs may drive cancers based on their ability to increase genome instability—a known cancer-driving phenotype (Hanahan and Weinberg, 2011).

As shown in Figure 3E, 32 of the 33 validated human DDPs are *E. coli* DDP homologs from the following categories: (i) 16% that are both known (Forbes et al., 2015) or predicted (D’Antonio and Ciccarelli, 2013) cancer drivers and amplified in TCGA cancers; (ii) 53% that are amplified in cancers and not known or predicted drivers; (iii) 6% known/predicted cancer drivers that are not known to be amplified in cancers; and (iv) 25% that are neither gene-amplified in cancers nor previously known or predicted drivers. None of these classes was predicted previously to be DNA-damage promoting, and some might not have been hypothesized to potentially promote cancer via overproduction (e.g., classes iii and iv). In Supplemental Discussion 7, we use the rates of validation in each class tested to estimate that there are likely to be many additional hDDPs among the 284-protein candidate hDDP network, that would test positive in our assays.

Bioinformatically, ~75% of the validated hDDP genes show cancer-associated copy-number increases in the TCGA patient-cohort data (Gao et al., 2013) (GISTIC threshold copy-number gain ≥ 1, Figure 3D). The fraction of cancer-specific or across-cancer copy-number-increased genes among the validated hDDP genes is higher than the candidates we tested that were not validated (Figure 3D, *p* = 0.02, one-way Fisher’s exact test). The data provide support for our hypothesis that validated hDDP genes may drive cancer via DNA damage when overexpressed, and imply that our cell-based DNA-damage assays relate to human cancer biology.

### Functional Systems Biology

Having identified DDPs in *E. coli* and human, we used the tractable *E. coli* model for further function discovery. We sought to create a multi-parameter, minable data set of phenotypes to bin the 208 *E. coli* DDPs into function clusters that reflect the kinds, causes, and consequences of DNA damage provoked by their overproduction. Our phenotypes are based on seven quantitative functional assays, many at the single-cell level (Figure 4). Two of these employ synthetic proteins that trap, fluorescently label, and allow quantification, as well as genomic mapping, of specific DNA-damage-intermediate structures in single living cells (Shee et al., 2013; Xia et al., 2016). We use the data to predict, then demonstrate, mechanisms by which dysregulation of diverse conserved proteins increase endogenous DNA damage.

**Figure 4.**
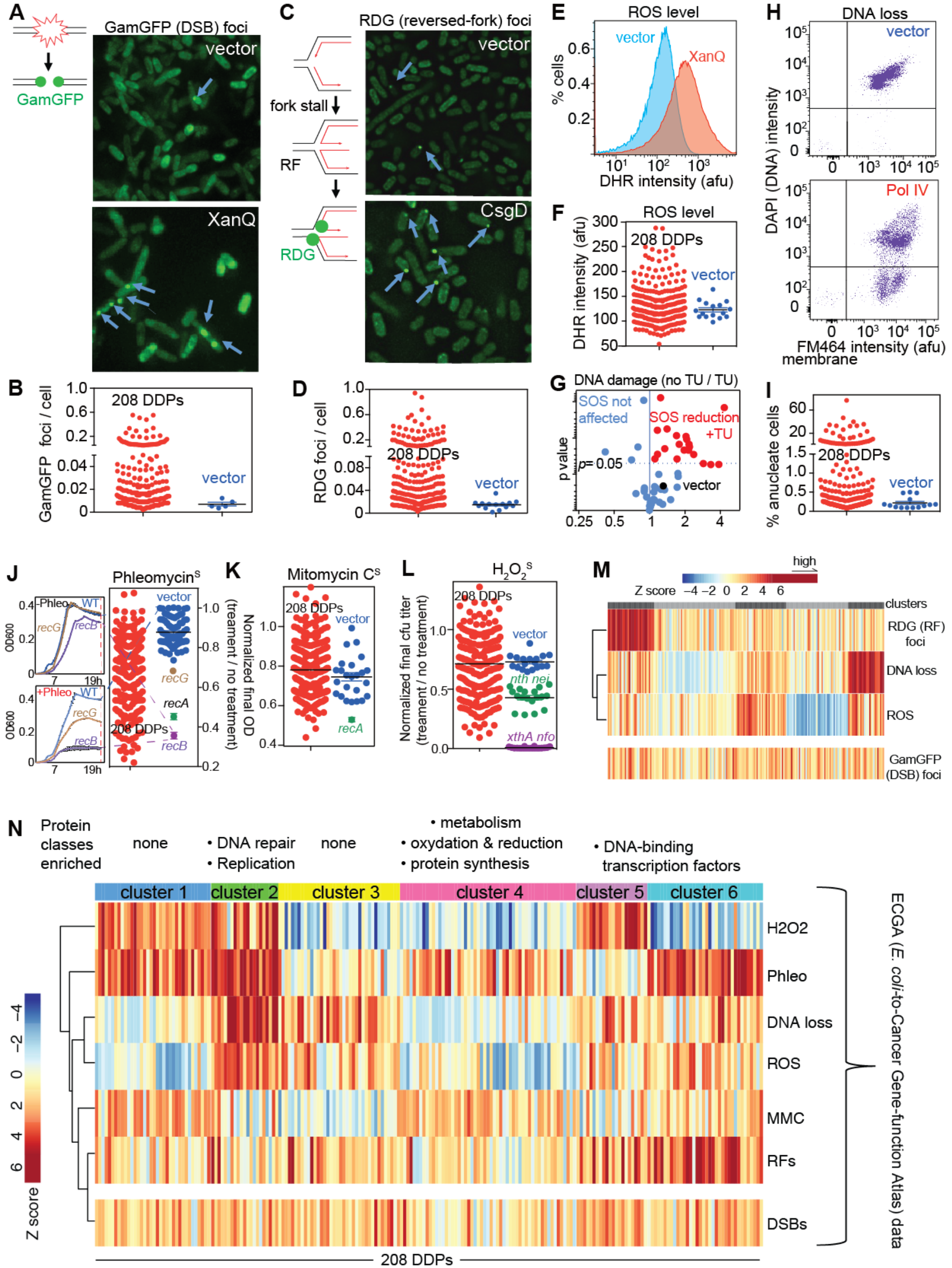
Kinds, Causes and Consequences of DNA Damage from *E. coli* DDPs Reveal Function Clusters. (A) Identification of DDPs that increase DNA double-strand breaks (DSBs), detected as GamGFP foci, per (Shee et al., 2013). Representative image in cells overproducing DDP XanQ, a membrane-spanning transporter. Diagram: lines, DNA strands; green balls, GamGFP. (B) 87 of the 208 *E. coli* DDPs promote DSBs. Quantification of GamGFP foci in 208 DDP-overproducing clones (means of two experiments, >1000 cells each; data by clone Table S1). (C) Detection of stalled, reversed replication forks (RFs) as RDG foci in *recA*^−^ cells. Engineered protein RuvCDefGFP (RDG) traps 4-way DNA-junctions, and in *recA* cells detects only RFs (Xia et al., 2016). Representative image in cells overproducing DDP CsgD. Diagram: lines, DNA strands; red lines, newly synthesized strands; green balls, RDG. (D) 106 of the 208 *E. coli* DDPs cause fork stalling and reversal. Quantification of RDG foci in 208 DDP-overproducing clones (data by clone, Table S1). (E-I) Flow-cytometric assays for— (E) elevated ROS measured as peroxide by dihydrorhodamine (DHR) fluorescence. Representative flow-cytometric histogram, XanQ overproducer, and (F) quantified in all 208 DDP-overproducing clones, mean fluorescence intensity; afu, arbitrary fluorescence units (2 experiments, mean) shows 56 (27%) of DDP clones (data by clone, Table S1). (G) Increased DNA damage in 16 of 56 high-ROS DDP clones is reduced by ROS-quenching agent thiourea (TU) indicating that the high ROS underlie the DNA damage. *P* values, unpaired two-tailed *t* test. Data by clone, Table S1. (H) DNA loss: the fraction of cells with no DNA (anucleate cells), representative example, Pol IV overproducer (DAPI indicates DNA, membrane dye FM464 indicates cells); events below the horizontal line scored as anucleate, DNA loss, (I) quantified in all 208 DDP-overproducing clones (2 experiments, mean), showing 67 (32%) of DDP clones (data by clone, Table S1). (J-L) Sensitivity to DNA-damaging agents in DDP-overproducing clones implies DNA-repair-pathway reduction (possible saturation) as a potential consequence of elevated DNA damage. Positive controls: relevant DNA-repair-defective mutants indicated. Each measured as slowed growth curves per J left (STAR Methods). Data by clone, Table S1. (J) Phleomycin sensitivity (reduced homology-directed DSB repair) seen in 106 of the DDP clones (51%). (K) Mitomycin C (MMC) sensitivity (reduced nucleotide-excision repair and/or homology-directed repair) seen 10 of the DDP clones (5%). (L) H_2_O_2_ sensitivity (reduced base-excision repair) seen in 75 of the DDP clones (36%). (M) Stalled replication (RFs) is clustered among particular DDP overproducers; DNA breakage is not. Progeny clustering (STAR Methods). (N) Cluster analysis of Z (significance) scores of assays: H_2_O_2_, hydrogen-peroxide sensitivity; Phleo, phleomycin sensitivity; DNA loss (anucleate cells); ROS (ROS levels); MMC, mitomycin-C sensitivity; RFs (reversed forks), RDG (RF) foci; DSBs, GamGFP (DSB) foci. Vertical bars along the x axis: the phenotype of each DDP clone. The 6 clusters indicate 6 DNA-damage signatures and suggest at least 6 different mechanisms of DNA-damage generation in the DDP network. Protein categories significantly increased in each cluster shown above (one-way Fisher’s exact test), cluster 2 at *p* = 0.01, cluster 4 at *p* = 0.01, and clusters 5 and 6 at *p* = 0.03.

### Proteins that Instigate DNA Double-Strand Breaks

We used the engineered double-strand-break (DSB)-end-specific binding protein GamGFP (Shee et al., 2013) to quantify DSBs in single living cells. GamGFP “traps” DSB ends and prevents their repair in *E. coli* and mammalian cells (Shee et al., 2013). GamGFP produced from a chromosomal regulatable gene cassette labels one-ended and two-ended DSBs as fluorescent foci in *E. coli* at an estimated 70% efficiency (Shee et al., 2013) (i.e., 30% of DSBs present are not seen as foci). We quantified GamGFP foci in each of the 208 DDP-overproducing clones by automated microscopy (STAR Methods). We found that 87 of the 208 DDP-overproducing clones displayed significantly more GamGFP foci than vector-only controls (Figures 4A-B and S4A, Table S1, Supplemental Discussion 9) and than 25 random SOS-negative (non-DDP) clones, none of which had increased GamGFP foci (Figure S4A). Our finding of 121 DDP-overproducing clones without increased GamGFP foci suggests that single-stranded (ss)DNA, the inducing signal for the SOS DNA-damage response, frequently accumulates at sites other than DSBs. The 41% of *E. coli* DDP clones with elevated GamGFP DSB foci are not enriched in any gene-function category (Table S1), implying that DSBs—a common result of many DNA-damaging mechanisms (Merrikh et al., 2012)—result from various cellular processes.

### Stalled Reversed Replication Forks

Stalling of DNA replication leads to DNA damage (Jackson and Bartek, 2009) and can create fourway DNA junctions when stalled replication forks “reverse” such that the new DNA strands basepair with each other (Figure 4C, reversed fork, RF) (Seigneur et al., 1998). Reversed forks (RFs) block resumption of DNA replication, lead to replication-fork breakage (Seigneur et al., 1998; Yeeles et al., 2013), and so are both a kind of DNA damage, and reflect a cause of DNA damage—stalled replication. We quantified stalled, RFs as fluorescent foci of the engineered 4way-junction-specific DNA-binding protein RuvCDefGFP (RDG) in cells lacking homology-directed-repair protein RecA (Δ*recA*), in which essentially all RDG foci represent RFs (Xia et al., 2016). RDG labels 4-way junctions with about 50% efficiency in live *E. coli* (Xia et al., 2016). We found that 106 of the 208 DDP-overproducing clones showed increased RDG (RF) foci relative to the vector-only control (Figures 4D and S4B, Table S1) and to 30 control, SOS-negative overproducing clones (Figure S4B; Table S1). Among the 106 clones with increased RDG foci, 49 also show increased DSBs (Table S1), detected as GamGFP foci, showing significant correlation (*p* = 0.03, r=0.15 Spearman’s correlation). Overall, at least 51% of DDPs promote replication stalling on overproduction, indicating the importance of DNA replication to DNA-damage generation by many of the overproduced proteins, but also suggesting that mechanisms in addition to replication may underlie DNA damage induced by dysregulation of many other DDPs.

### Reactive Oxygen via Transporters and Metabolism

We quantified intracellular levels of reactive oxygen species (ROS) in single cells using the peroxide-indicator dye, dihydrorhodamine (DHR) (Gutierrez et al., 2013) and flow cytometry (Figures 4E, F and S4C, D). We found that 56 of the overproduced DDPs caused increased ROS levels (Figures 4F and S4C, D). We show that the high ROS levels contribute to DNA damage in at least 16 of the 56 ROS-elevated DDP-producing clones, in which the DNA damage was significantly reduced by the ROS-quenching agent thiourea (Keren et al., 2013), compared with the vector-only control (Figure 4G). Thus, high endogenous ROS levels underlie DNA damage in a subset of the DDP-producing clones. These (Figure 4G) comprise five membrane-spanning transporters (investigated below), an excess compared with the prevalence of transporters among *E. coli* proteins (*p*=0.002, hypergeometric test). The other eleven proteins relate to metabolism processes (Table S1), implying that perturbation of metabolic pathways can cause DNA damage by increasing ROS.

### DNA Loss

Loss of DNA in cells can result from various problems including chromosome-segregation failure (Joshi et al., 2013), for example, from incomplete DNA replication or incomplete homology-directed repair between chromosomes, either of which can leave two chromosomes attached at cell division (Hendricks et al., 2000). We identified 67 DDP clones with increased frequencies of DNA-depleted (“anucleate”) cells (Figures 4H, I and S4E, F; Table S1) using flow cytometry of single cells with DNA and cell membranes stained separately. Overproduced DNA-repair and replication proteins are enriched among these clones (*p* = 0.04, one-way Fisher’s exact test, Table S1), implying that excessive DNA-repair and replication proteins promote DNA damage that leads to DNA erosion or chromosome-segregation failure. Their overproduction might alter stoichiometry of and hinder DNA-repair or replication complexes, and/or perturb DNA directly.

### Reduced DNA-repair Capacity via Specific Proteins

Reduction of DNA-repair capacity could increase DNA-damage levels, and could result either from saturation of repair pathways with excessive DNA damage, or from overproduction of proteins that interfere directly with DNA-repair. Either would mimic repair-deficiency (and mutator phenotype), without a DNA-repair-gene mutation. We assayed DNA-repair capacity indirectly, as sensitivity to DNA-damaging agents that produce damage repaired by specific DNA-repair mechanisms: DSB-, ssDNA-break-, and ROS-instigator phleomycin (Steighner and Povirk, 1990); base-oxidizing agent hydrogen peroxide (H_2_O_2_); and DNA cross-linking agent mitomycin-C (MMC). These cause DNA damage repaired by homology-directed repair (HR, phleomycin), base-excision repair (“BER”, H_2_O_2_), and, for MMC, both nucleotide-excision repair and HR (Friedberg et al., 2005). We found that 106, 75, and 10 of the 208 DDP clones were sensitive to phleomycin (Figures 4J and S5A-B), H_2_O_2_ (Figures 4L and S5C), and MMC (Figures 4K and S5D, E), respectively, shown by reduced cell densities in cultures (normalized for any effects of protein overproduction on overall growth rates, STAR Methods). Collectively, 140 DDP-overproducing strains were sensitive to at least one DNA-damaging agent (Figure S5G; Table S1), and 45 to multiple drugs (Figure S5G; Table S1). Non-DDP overproduction clones were not enriched for sensitivity to DNA-damaging agents and differ from the DDP-network group for sensitivity to phleomycin and H_2_O_2_, but not for MMC (Figure S5). The data suggest that overproduction of various DDPs provoke different kinds of DNA damage that overwhelm or inhibit distinct DNA-repair mechanisms.

We excluded the possibility that most DNA-damage sensitivities resulted from DDP-induced heritable mutation, by showing no sensitivity *after* removal from DNA damage (Figure S5H, Supplemental Discussion 10). We also excluded transcriptional downregulation of DNA-repair genes, by RNA-seq of DNA repair genes (Figure S5F, Supplemental Discussion 10). The data suggest that specific DNA-repair pathways are either inhibited directly or saturated by DNA damage caused by overproduction of specific DDPs and imply that dysregulating diverse proteins, such as by gene copy-number alteration, can create DNA-repair deficiency without mutation of DNA-repair genes.

### Clustering *E. coli* Function Data Implicates Mechanisms

We grouped the quantitative data from the functional assays using stability-based clustering [(Hu et al., 2015), Progeny clustering] (Figures 4M,N and S5G). The quantitative data on RDG (RF) foci analyzed with three other quantitative parameters measured in single-cells—ROS levels, DNA loss, and DSBs (all discussed above)—revealed that high RF loads are enriched in a specific cluster (Figure 4M). The RF-dense cluster is significantly enriched for DNA-binding transcription factors (examined below), with 29%, compared with 12% among the network as a whole (*p* = 0.002, oneway Fisher’s exact test, Figure 4M; Table S1). The data indicate that distinct protein functions preferentially stall replication.

Grouping all quantitative data sets revealed six discreet function clusters (Figure 4N; Table S1), which may indicate at least six different potential mechanisms and cellular consequences of DNA-damage promotion by DDPs (reduced DNA-damage classes discussed Supplemental Discussion 11). We compared the function clusters with protein-protein-association data in the DDP network (Figures 4N and S2A, Table S1). Whereas the entire DDP network shows significantly more protein-protein associations than sets of 208 random *E. coli* proteins, superimposition of the function clusters onto the protein-protein interaction network indicates that cluster 2 shows even more protein-protein interactions than the DDP network as a whole (*p* = 0.0007, one-was Fisher’s exact test). These data support associations of function clusters with particular biological mechanisms, three examined below.

### Transcription Factor Binding to DNA Promotes Replication Stalls

Clusters 5 and 6 of Figures 4N (listed Table S1) show increased replication stalling/reversed forks (RFs, per Figure 4M) and are most enriched for DNA-binding transcription factors: transcriptional activators and repressors. We hypothesized that persistent binding of a protein to DNA might create a replication “roadblock”, stall forks, and cause RFs. RFs can be regarded as DNA damage and also cause additional DNA damage when cleaved by endonucleases (Seigneur et al., 1998). In support of this hypothesis, we found, first, that mutational ablation of the DNA-binding-domains (DBDs) of three of the transcription factors—CsgD, HcaR and MhpR—abolished both their abilities to promote SOS-inducing DNA damage (Figures 5A-C), and RDG (RF) foci (Figures 5D-E and S6A). The data indicate that these transcription factors must bind DNA to provoke DNA damage and RFs upon overproduction. Second, we created an mCherry (red) fusion of the CsgD transcription factor (Ogasawara et al., 2011), and see that it forms foci DBD-dependently (Figure 5F and S6B), suggesting that foci reflect the DNA-bound transcription factor. We found that most of the CsgD-mCherry foci, and also HcaR-mCherry foci, co-localized with RDG (RF) foci (Figures 5F-H), suggesting that RFs form near the DNA-bound transcription factors. Foci of DNA-bound proteins are distinguishable at ~50kb apart on DNA, e.g., (Shee et al., 2013); thus, these data indicate that RDG/RF foci accumulate in the vicinity of the sites of transcription-factor binding to DNA (Figures 5G and 5H). High resolution mapping of RDG (RFs) by ChIP-seq in the genome of CsgD-overproducing cells showed RDG (RFs) enriched near the transcription factor’s target DNA-binding sites CsgD-DBD-dependently (Figure 5I), supporting the hypothesis that the bound transcription factor stalls replication causing fork reversal nearby. CsgD has 10 experimentally well-characterized binding sites (Ogasawara et al., 2011), and we found that the CsgD-DBD-dependent RDG (RF) ChIP-seq peaks are very significantly enriched in 10kb regions surrounding known CsgD DNA-binding sites (representative peaks, Figure 5I; Supplemental Discussion 12, rest of known sites; Figure S7). CsgD-DBD-dependent RDG (RF) ChIP-seq peaks occurred both upstream and downstream of the binding sites in the replication paths (Figure S7, discussed Supplemental Discussion 12, and Figure 5J). Our data support a model (Figure 5J) in which overproduced DNA-bound transcription factors can create roadblocks to replication, which leads to increased fork stalling and reversal near where the transcription factors bind, causing DNA damage.

**Figure 5.**
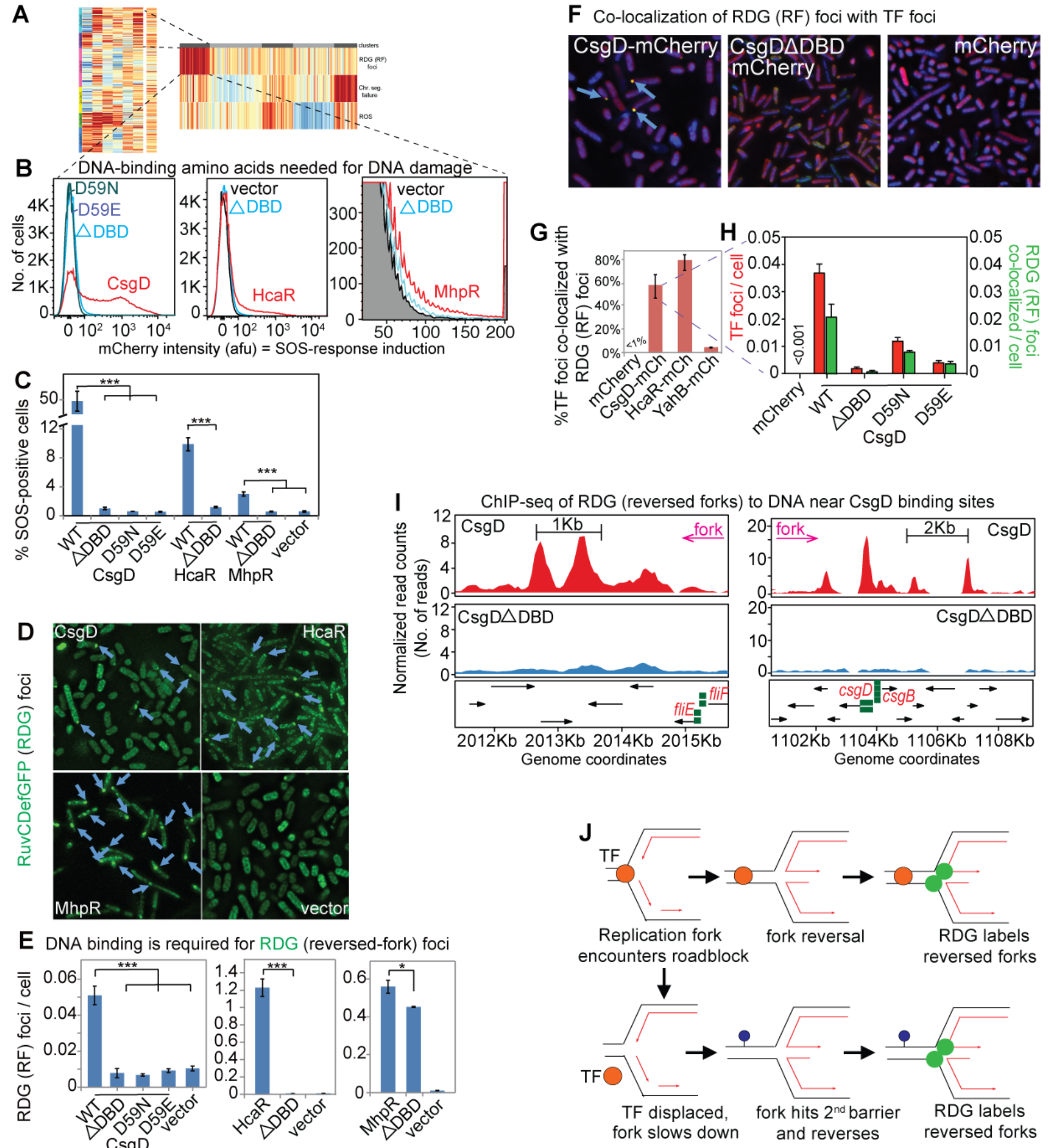
*E. coli* Transcription Factors Promote Replication-fork Stalling and Reversal DNA-binding-domain Dependently. (A) DNA-binding transcription factors (TFs) are enriched among DDP clones with high RFs (*p* =0.002, one-way Fisher’s exact test). (B) DNA-binding ability is required for DNA damage/SOS activity caused by overproduced DNA-binding TFs. Representative flow cytometry histograms of SOS induction, three TFs and their corresponding mutants: ΔDBD, DNA-binding domain in-frame deletion; D59N, D59E, single amino-acid changes that reduce CsgD DNA-binding (Ogasawara et al., 2011). (C) Mean ± SEM of ≥3 experiments. (D) DNA-binding ability of overproduced TFs is required for increased RDG (RF) foci (blue arrows). Representative images. Figure S6A, all genotypes. (E) Mean ± SEM of ≥3 experiments. (F and G) mCherry-protein fusions of DNA-binding TFs form foci that co-localize with RGD (RF) foci, placing TF binding and RFs in relative proximity (within 50kb, text) in the 4.6MB *E. coli* genome. (F) Representative data. CsgD-mCherry foci co-localize with RDG foci dependently on the CsgD DNA-binding domain (DBD). Blue arrows, co-localized red and green (CsgD-mCherry and RDG) foci. Figure S6B, all genotypes. (G) Mean ± SEM of ≥3 experiments with CsgD-, HcaR-, and YahB-mCherry co-localization with RDG. (H) Co-localization of CsgD-mCherry foci with RDG foci requires CsgD DNA-binding ability; quantification. (I) RDG ChIP-Seq peaks in HR-defective Δ*recA* cells (RFs) are enriched near CsgD-binding sites (green squares; *p* =0.01, two-tailed z-test compared with simulated data, see Supplemental Discussion 12). Representative peaks shown; Figure S7 for the complete set of RF peaks. (J) Model: overproduced TFs (orange circles) binding to DNA (parallel lines, basepaired strands) cause RFs by replication roadblock. Green circles, RDG bound to RF.

### *E. coli* and Human Transporter Overproduction Elevates ROS

Membrane-spanning transporters are the largest category of human homologs of the *E. coli* DDPs, and several are both overrepresented among known cancer drivers and also provoke DNA damage on overproduction (Figures 2A, Tables S2 and S3). We found that *E. coli* membrane transporters are overrepresented at 26% in the high-ROS cluster in Figure 4M compared with 11% over the whole network (*p* = 0.004, one-way Fisher’s exact test, Figure 6A-C; Table S1). Further, sixteen DDP clones with high ROS caused DNA damage ROS-dependently in that the damage was reduced by ROS quenching (Figure 4I). These include five transporters, the increased ROS and ROS-dependent DNA damage of which are shown in Figures 6B-D. Three of the five are H^+^ symporters, a significant enrichment compared with the frequency of H^+^ symporters encoded in the genome (Keseler et al., 2017) (*p* = 2.7×10^−5^, hypergeometric test), one transports polypeptides, and the remaining one Mg^2+^.

**Figure 6.**
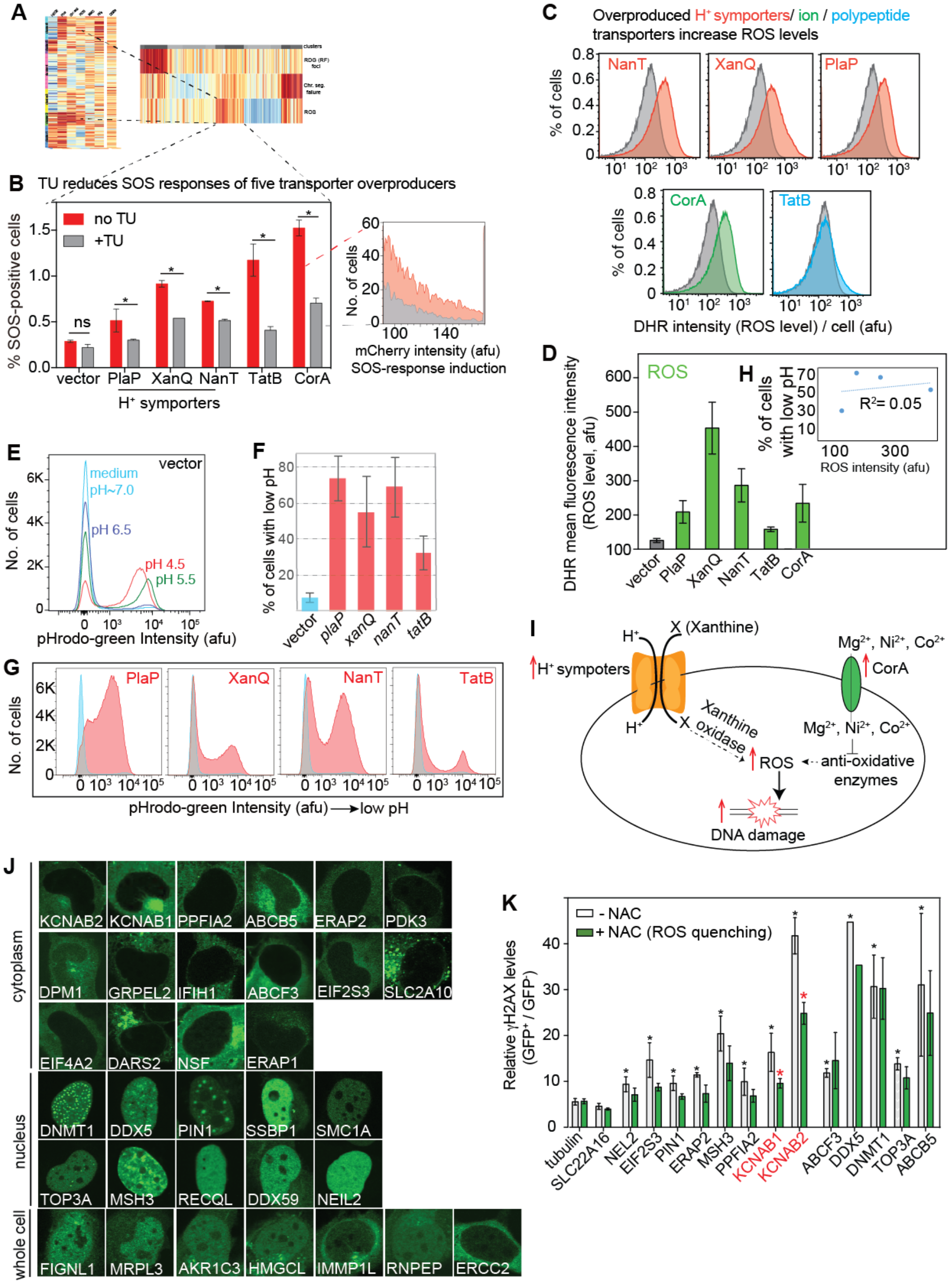
*E. coli* and Human Transmembrane Transporters Promote DNA Damage via Increased ROS. (A) *E. coli* high ROS cluster is enriched for membrane-spanning transporters (*p* = 0.004 one-way Fisher’s exact test). (B) DNA damage (SOS activity) from five overproduced *E. coli* transporters is partially reversed by ROS-scavenger thiourea (TU), implying ROS-dependent DNA damage. Quantification of flow cytometry per blow up. Mean ± range, 2 experiments. Blow up: representative flow cytometry for DNA damage: cells with chromosomal SOS-promoter-mCherry fusion. (C) ROS levels increase upon overproduction of various *E. coli* membrane-spanning transporters. ROS measured as H_2_O_2_ shown by DHR stain and flow cytometry. Representative data (Table S1 for all). Gray, vector only; red, H^+^ symporters; green, ion transporter; cyan, polypeptide transporter. (D) Means ± range of 2 experiments. (E-H) Increased *E. coli* H^+^ symporter activity is caused by overproduction, shown by reduced cellular pH. (E) Detection of *E. coli* intracellular pH by pHrodo-green dye staining followed by flow cytometry. Cells with the vector were exposed to buffers with varied pH levels and pHrodo-green dye was used to stain the cells. Control vector-bearing cells exposed to low pH cells had subpopulations with increased pHrodo-green intensity. (F, G) Reduced intracellular pH in clones overproducing the PlaP, XanQ, or NanT H^+^ symporters, or the TatB polypeptide transporter, indicates that overproduction causes gain-of-function, overall increased activity per cell of these transporters. Cyan, vector only. (F) Quantification: mean ± range of two experiments. (G) Representative flow-cytometry histograms. (H) Reduced pH in *E. coli* transporter-overproducing clones—the three H^+^ symporters and TatB polypeptide transporter—is not correlated quantitatively with increased ROS (R^2^=0.05, Pearson’s correction analysis), suggesting that the specific cargoes, not low pH, promote DNA damage. (I) Models for ROS-dependent DNA-damage promotion by overproduction of the *E. coli* XanQ and CorA transporters. Left: Overproduced H^+^ symporter XanQ might cause ROS by increased import of xanthine which is oxidized by the ROS-generating xanthine oxidase (Kelley et al., 2010). Right: CorA overproduction might cause DNA damage via increased import of Ni^2+^, which inhibits anti-oxidative enzymes (Schmidt et al., 2009) and causes DNA damage ROS-dependently (Cameron et al., 2011). (J) Overproduced validated human (h)DDPs are localized in various subcellular compartments, implying various DDP mechanisms. Of the 33 overproduced validated hDDPs, 16 were detected only in the cytoplasm, 10 only in the nucleus, and 7 throughout the cell. All show qualitatively repeaTable Subcellular localization. (K) ROS underlie at least some of the DNA damage caused by human KCNAB1/2 transporter overproduction. NAC: N-acetyl-cysteine, an ROS quencher (STAR Methods). * *p* < 0.05 relative to the NAC-untreated GFP-tubulin control; * *p* < 0.05 relative to the corresponding NAC-untreated control.

Proton (H^+^) symporters import molecules concurrently with H^+^. We found that overproduction of each of the three H^+^ symporters conferred reduced intracellular pH (increased H^+^) (Figures 6E-G), implying that overproduction increased their symporter activities. However, their induction of ROS was not well correlated with their reduction of pH (Figure 6H), suggesting that other cargos that they import may cause the increased ROS and DNA damage, or that simply compromising membrane integrity and cellular boundaries may provoke ROS and DNA damage. A specific model for XanQ, the strongest ROS-promoter among them, is illustrated in (Figure 6I), discussed Supplemental Discussion 13. Overall, the data reveal that DNA-damage induction can result from increased transporter activity, leading to high levels of DNA-damaging ROS (Figures 6A-C). The data suggest that disturbing cellular boundaries can cause DNA damage via ROS.

CorA, an inner membrane Mg^2+^ transporter, which transports Co^2+^ and Ni^2+^ less efficiently (Kehres and Maguire, 2002), elevates ROS (Figures 6C-D) and DNA damage (Figure 6B) when overproduced. Both might occur by increasing the usually minimal import of Co^2+^ and Ni^2+^ (Figure 6I). Ni^2+^ is toxic and induces DNA damage via oxidative stress (Cameron et al., 2011) because Ni^2+^ binds sulfhydryl groups commonly found in anti-oxidative enzymes (Schmidt et al., 2009). Increased Mg^2+^ import is unlikely to underlie the ROS and DNA damage because Mg^2+^ is the most abundant metal ion in cells and excessive Mg^2+^ does not seem to affect the activities of the many Mg^2+^-utilizing enzymes (Hartwig, 2001).

In human cells, multiple DNA-damaging mechanisms are implicated. First, a survey of the subcellular localization of all 33 overproduced, validated hDDPs showed that 16 were cytoplasmic; 10 were nuclear, and 7 were found throughout the cell (Figures 6J), suggesting that direct contact with DNA is not needed for many overproduced hDDPs’ instigation of DNA damage.

We screened a sample of thirteen validated hDDP-producing clones for those with DNA damage that could be suppressed by ROS quencher N-acetyl cysteine (NAC), to identify those that instigate ROS-dependent DNA damage. We found that overproduction of a membrane-spanning transporter promotes DNA damage via ROS. KCNAB1/2 promote high DNA damage (Figure 6K), are cytoplasmic when overproduced (Figure 6J), and are subunits of intracellular voltage-gated K^+^ channels that function in redox transformations of xenobiotics (Hlavac et al., 2014). Increased *KCNAB2* mRNA is found in breast cancer (Hlavac et al., 2014), but how KCNAB1/2 overproduction might promote cancer is unknown. We found that DNA-damage promotion by KCNAB1/2 relies at least partly on ROS, in that ROS-quenching NAC treatment reduced DNA-damage induction by KCNAB1 or KCNAB2 overproduction (Figure 6K). Cells overproducing several other validated hDDPs showed no reduction in DNA damage with NAC treatment (Figure 6K), indicating that these DDPs promote DNA damage by other or additional mechanisms.

### *E. coli* Pol IV and Human DNMT1 Promote DNA Damage via Replisome-Clamp Interaction

DNA polymerase (Pol) IV, encoded by *dinB*, is among the highest DNA-damage generators in the *E. coli* DDP network (Figures 7A-C; Table S1), generating both DSBs seen by increased GamGFP foci (Table S1) and ssDNA gaps, inferred as follows. SOS DNA-damage-response induction by overproduced Pol IV was partially RecB- (DSB) and partially RecF- (ssDNA-gap) dependent (Figures 7D-E); and RecB and RecF are required for SOS induction by DSBs and ssDNA gaps, respectively (McPartland et al., 1980), implicating DSBs and ssDNA gaps as Pol IV-induced damage. Overproduced Pol IV also promotes chromosome loss (Figure 4E). We show, that Pol IV-induced DNA damage occurs via its interaction with the replisome sliding clamp as follows.

Pol IV is a poorly processive, low-fidelity DNA polymerase that traverses specific damaged bases that more efficient DNA polymerases cannot copy (Wagner et al., 2000). Pol IV competes with more processive DNA polymerases (Frisch et al., 2010; Hastings et al., 2010), by competition for binding the replisome sliding clamp protein, beta (Dohrmann et al., 2016; Heltzel et al., 2012). We hypothesized that the documented ability of Pol IV to slow replication-fork progression (Heltzel et al., 2012; Uchida et al., 2008) might underlie its generation of DNA damage when overproduced. In support of this hypothesis, we found, first, that overproducing Pol IV simultaneously with DNA Pol II, its competitor for the replisome (Frisch et al., 2010), caused a roughly 50% reduction in the Pol IV-dependent DNA damage (Figures 7B-C). Second, cells producing a mutant replisome clamp-loader protein with reduced Pol IV loading and increased loading of the major replicative DNA polymerase, Pol III (Dohrmann et al., 2016), also showed reduced DNA damage from Pol IV overproduction (Figures 7D-E, *dnaX* tau-only), but no reduction of SOS on its own (Figure 7F). Third, a C-terminal Pol IV deletion that abolishes Pol IV interaction with the replisome beta sliding clamp (Uchida et al., 2008) also abolished DNA damage upon Pol IV overproduction (ΔBBD, Figures 7B-C). We conclude that Pol IV interaction with the replisome clamp is required for its generation of DNA damage on overproduction.

**Figure 7.**
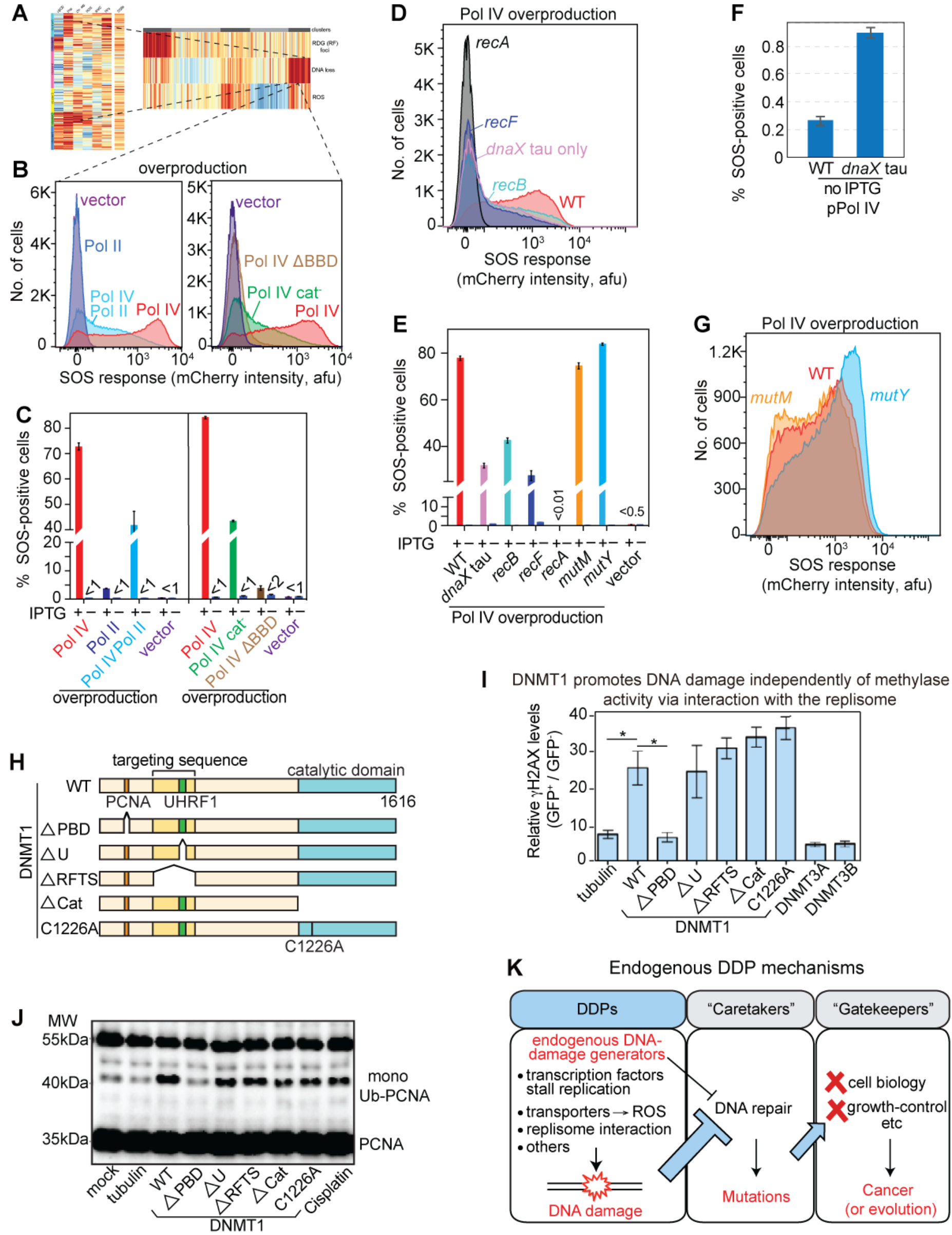
*E. coli* DNA Pol IV and Human DNMT1 Promote DNA Damage via Interaction with the Replisome Clamp. (A) *E. coli* function cluster with high DNA loss includes DNA Pol IV. (B) *E. coli* Pol IV promotion of DNA damage is reduced by co-overproduction of its competitor at the replisome, DNA Pol II (left). Right: Pol IV promotion of DNA damage requires its interaction with beta replisome sliding clamp, seen by dependence on the Pol IV beta clamp-binding domain (BBD), and is partly independent of Pol IV catalytic activity (cat^−^ mutant). Representative data. (C) Mean ± SEM of ≥3 experiments. (D) Reduction of *E. coli* Pol IV-induced DNA damage in a clamp loader tau-only mutant, which interacts with replicative DNA Pol III in preference to Pol IV (Dohrmann et al., 2016). RecB- and RecF-dependence of the SOS response induced by overproduced Pol IV implicates DSBs and single-strand gaps, respectively, as DNA-damage types produced. Representative data. (E) Mean ± SEM of ≥3 experiments. Pol IV is induced by IPTG. (F) The clamp-loader *dnaX* tau-only mutant is not generally deficient in SOS-response induction, but merely reduces the DNA damage (SOS activity) caused by Pol IV upregulation (D, E). (G) Neither the *E. coli* MutM 8-oxo-dG glycosylase nor the MutY adenine glycosylase are required for DNA damage instigated by Pol IV, indicating DNA damage produced independently of incorporation of 8-oxo-dG into DNA. Representative data, quantified (E). (H) Constructs for production of human wild-type and truncated DNA methyltransferase DNMT1 in human cells. PBD, PCNA-binding domain; U, UHRF1, ubiquitin-like containing PHD and RING-finger domains 1 binding domain; RFTS, replication-focus-targeting sequence, recruits DNMT1 to DNA-methylation sites; Cat, catalytic domain for methyltransferase activity; C1226A, mutation of the catalytic active site. (I) Human DNMT1 overproduced in human cells promotes γH2AX (DNA-break indicator) accumulation methylase-independently and replisome-clamp-interaction dependently. Overproduction of two DNMT1 catalytically dead mutants increased DNA damage similarly to overproduced WT DNMT1. Overproduction of two other de novo DNA methyltransferases (DNMT3A, DNMT3B) did not elevate DNA damage. DNA-damage promotion by DNMT1 requires the DNMT1 PBD domain, required for DNMT1 binding to the human replisome clamp: PCNA. (J) Human DNMT1 overproduction promotes PCNA monoubiquitination, a replication-stress indicator, in a replisome-interaction-dependent manner. Monoubiquitination determined by western blot with anti-PCNA antibody (STAR Methods). (K) The endogenous DDP model for cancer promotion and mechanisms: DDPs a cancer-protein functional class upstream of DNA repair. Excessive endogenous DNA damage is proposed to titrate (thick blue -|) or inhibit (thin black -|) DNA repair causing DNA-repair (“caretaker”)-protein deficiency in cells without a DNA-repair-gene mutation. Repair deficiency increases mutation rate, leading to cancer- (or evolution-) driving mutations in the cell-biology altering “gatekeeper” genes that cause the cancer cell-biological phenotypes.

Perhaps surprisingly, Pol IV catalytic activity (DNA synthesis) was not required for all of its DNA-damage induction; the catalytically-inactive Pol IV R49F mutant (Wagner et al., 1999) reduced only half the DNA damage caused by overproduction (Pol IV cat^−^, Figures 7B and 7C), without reducing Pol IV protein levels (Uchida et al., 2008; Wagner et al., 1999) (Figure S7D). Thus, mere binding of Pol IV to the replisome, not only its poorly processive catalytic activity, appears sufficient to cause DNA damage. The synthesis-dependent component of DNA damage might have resulted from the ability of Pol IV to incorporate oxidized guanine (8-oxo-dG) into DNA, per (Foti et al., 2012), which leads to two different strand-breaking BER processes that begin with base removal by MutM and MutY DNA glycosylases (Foti et al., 2012). However, loss of neither glycosylase diminished DNA damage caused by Pol IV overproduction (Figures 7E and 7G). The data imply that BER following 8-oxo-dG incorporation is not how excess Pol IV promotes most DNA damage. Overall, Pol IV promotes DNA damage dependently on replisome-clamp interaction, and only partly dependently on catalysis (model, Figure S7E).

We found that human DNMT1 overproduction also induces DNA damage in human cells based on binding the replisome clamp, and independently of its catalytic activity: a non-canonical potential cancer-driving role. DNMT1 is the major human DNA methyltransferase that methylates DNA upon replication (Jin and Robertson, 2013). Hypomorphic mutations in *DNMT1* promote microsatellite instability (Jin and Robertson, 2013). Increased DNMT1 is common to several cancer types, and causes hypermethylation, proposed to downregulate tumor-suppressor genes (Biniszkiewicz et al., 2002), which would constitute a cell-biological, rather than DNA-based, tumor-promoting role. Surprisingly, we found that, in addition, DNMT1 promotes DNA damage independently of its DNA-methylation activity, in that overproduction of DNMT1 catalytically-dead mutant proteins (Figure 7H) increased DNA damage similarly to wild-type DNMT1 (Figures 7I and 7J). Overproduction of two other DNA methyltransferases did not increase DNA-damage levels (Figure 7I). DNMT1 truncations (Figure 7H) revealed that DNMT1 promotion of DNA damage required its PCNA-binding domain (PBD), which binds the replisome sliding clamp: PCNA (Figures 7I, S3L and S3M). Rad18-mediated monoubiquitination of PCNA, a DNA damage response (Mortusewicz et al., 2005), also resulted from DNMT1 overproduction, also methylase-independently and requiring the DNMT1 PBD (Figures 7J and S3M). These data suggest that mere binding of overproduced DNMT1 to the replisome clamp promotes DNA damage, and a resulting DNA-damage response, independently of methylation. The finding that both *E. coli* DNA polymerase IV and human DNMT1 promote DNA damage dependently on replisome-clamp binding and independently of their catalytic activities (Figure 7A-G) indicates generality of this mode of generation of DNA damage. The data suggest that promotion of DNA damage by DNMT1-PCNA complexes (Figures 7H-J), and resulting mutagenesis (Figure 3C), may promote cancers other than or in addition to via the known cell-biological/regulatory function of DNMT1 in DNA methylation, and that many clamp-binding proteins may act similarly when their genes are overexpressed/amplified.

## DISCUSSION

The identities and functions of proteins in the *E. coli* and human DNA-damaging-protein networks reveal that multiple diverse proteins, cellular processes, and molecular mechanisms underlie genesis of endogenous DNA damage. Although obtained in an overexpression screen, these are likely to represent natural causes of spontaneous endogenous DNA damage, first, because gene overexpression in cells in populations is remarkably common, on the order of tens of percents of bacterial cells for any given gene (Elowitz et al., 2002), with copy-number gains of any chromosomal region occurring in 10^−3^ of cells (Reams et al., 2010), and expected to be comparable in human cells (Hastings et al., 2009). Second, the association of the 284 human DDP homologs with four aspects of human cancer data indicates natural biological relevance (reviewed below).

### Endogenous DNA Damage in Cancer

The 284 human homologs of *E. coli* DDP genes are overrepresented among known (Forbes et al., 2015) and predicted (D’Antonio and Ciccarelli, 2013) cancer drivers (Figure 2C), overrepresented in cancers as amplified (Figures 2D and S3A-C, Table S3), and their increased expression in human cancers associated with poor outcomes (Figures 2E and S3D-F) and heavy mutation loads (Figure 2F). These associations support their overexpression being both biologically relevant, and relevant to cancer biology specifically.

The DDPs appear likely to represent a new broad function class of cancer-promoting proteins, and the earliest in cancer. Cancer-gene functions have been grouped into multiple specific categories (Hanahan and Weinberg, 2011) that fit into two broad classes of function (Kinzler and Vogelstein, 1997): the cancer-cell-biology-altering “gatekeepers”, mutations in which make cell biology more cancer-like, and the genomic “caretakers”—the DNA-repair genes, mutations in which elevate mutation rate and so drive cancer by increasing gatekeeper mutations (Figure 7K). The DDPs are expected to act *before* and upstream of DNA-repair functions promoting the endogenous DNA damage that necessitates repair (Figure 7K), and so instigating some of the earliest events in cancer development.

The large number of DDPs span diverse protein functions, the cancer-driving functions of many of which may be obscure or mis-assigned. Some of the mechanisms of DDP action may necessitate reëvaluation of their cancer driving roles, and also of the drugs designed to inhibit them. For example, we found that human DNA methyltransferase DNMT1 causes DNA damage independently of its methylation activity, via its interaction with the replisome sliding clamp (Figure 7H-J). Current cancer drugs against DNMT1 target the methylase activity (Jones et al., 2016), and not replisome binding. It is unclear which activities of DNMT1 promote cancer, and so which should be drugged. Our finding may inform the development of and use of DNMT1-targeting strategies, taking into account its multifunctionality, broadly across cancer types. These results underscore the importance of determining all functions of a protein that are cancer promoting.

### The *E. coli*-to-Cancer Gene-function Atlas of Bacterial and Human DDP Phenotypes

Our data from seven quantitative assays for kinds, causes, and consequences of endogenous DNA damage promoted by *E. coli* DDPs constitute a rich resource and framework for within- and cross species discovery of conserved DNA-damage-generating mechanisms. We used them to identify six main function or phenotype clusters (Figure 4N), and implicate, then test and demonstrate, three *E. coli* DDP mechanisms (Figures 5-7), two also identified in human cells (Figures 6 and 7). We created a minable web-based resource for searching the complete *E. coli* function data, and the functional data from the validated human DDPs: the *E. coli*-to-Cancer Gene-function Atlas (ECGA) (https://microbialphenotypes.org/wiki/index.php/Special:ECGA). The ECGA data can be searched via the bacterial proteins’ or their human homologs’ names or by function key words. ECGA can be used for querying/generation of hypotheses for *E. coli* and potential conserved human-protein functions, and as it develops in future, for other organisms.

### Mechanisms that Cause Endogenous DNA Damage

In *E. coli* DNA-binding transcription factors caused replication-fork stalling and reversal (Figure 5) by apparent blocking of replication forks by the bound transcription factors (Figure 5J). Though replication-transcription conflicts have been engineered by reversing chromosome segments (Tehranchi et al., 2010), engineering multiple transcription-factor binding sites into long arrays (Magnan et al., 2015), or knock out of RNA-removal proteins (Wahba et al., 2016), the kinds of DNA damage generated were not identified, nor was it known that any of these mechanisms could occur in natural genomes with only an endogenous protein upregulated, as often occurs in cells. Our results indicate that fork reversal is common and protein-function specific. Nearly 10% of human genes encode transcription factors (Levine and Tjian, 2003), including many cancer-driving overproduction (onco-)proteins, some known to promote DNA damage when overproduced, e.g., c-Myc, and E2F1 (Pickering and Kowalik, 2006; Vafa et al., 2002). Thus, based on our observation of transcription-factor binding and DNA-damage-production in *E. coli*, many onco-protein transcription factors might promote cancer similarly to *E. coli* transcription factors—by causing genome-destabilizing DNA damage, potentially RFs. Moreover, in *E. coli* and human cells, we discovered that increased transmembrane transporter activities of several different kinds elevate ROS levels causing DNA damage (Figures 4M and 6A-F). This previously unknown mechanism, also apparent with the human KCNAB1/2 transporter, might explain the KCNAB2 association with cancers (Hlavac et al., 2014)—a hypothesis that remains to be tested. Further, we found that both *E. coli* DNA polymerase IV (Figure 7A-G) and human DNMT1 (Figures 7H-J, and S3L and M) provoke DNA damage via binding their respective replisome sliding clamps when overproduced, independently of their catalytic activities. Disruption of the replisome leading to replication-fork collapse (or other means Figure S7E), is thought to be an important source of DNA damage (Kuzminov, 1995) likely to apply to dysregulation of many kinds of proteins.

### DNA Damage as Potential Cancer Biomarker

The existence of many diverse means of increasing endogenous DNA damage, and the predicted large sizes and diversity of both bacterial and human DDP networks indicate that dysregulation of any of many proteins is likely to be mutagenic via DNA damage (Figures 1J, 2F and S1F). Because many different proteins, processes, and mechanisms instigate DNA damage, DNA damage itself might be a robust predictor of cancer and genetic-disease susceptibility. The ability to detect high DNA damage could potentially make DNA-damage screening attractive for early identification of at-risk individuals, at a time before genome-sequencing would identify disease-associated mutations. Additionally, the success of cancer immune therapy “checkpoint inhibitors” is limited to high-mutagenesis cancers, apparently because mutagenesis creates diverse tumor antigens that can be attacked by the stimulated immune system (Germano et al., 2017). Thus, immune therapy is currently aimed primarily at some DNA-repair-defective cancers (Germano et al., 2017). Our data suggest that DNA damage, or upregulation of DDP network genes, may predict additional susceptibilities in various cancers.

## AUTHOR CONTRIBUTIONS

Conceptualization, S.M.R., J.X., K.M.M., L-Y.C., C.H.,; Methodology, J.X., S.M.R., C H., R.B.N., M.A.B., J.P.P., Y.Z., Y.W., Q.M., R.T.P., A.J., C.W.H., M.L.M.G., D.M.F., C.C., M.A.M., K.L.S., H.L., C.Q.; A.T.S., D.B., S.G.H., J.C.H., D.A.S; Investigation, J.X., L-Y.C., R.B.N, M.A.B.N., Q.M., C.W.H., M.P., D.M.F., J.P.P., Y.Z., Y.W., Q.M., A.J., A.M.L., C.W.H., M.L.M.G., M.C.J., M.R.,; Writing - Original Draft, S.M.R., J.X., K.M.M., L-Y.C.; Writing - Review & Editing, all authors; Funding Acquisition, S.M.R., K.M.M, C.H., D.B., P.J.H., C.Q., K.L.S., H.L., C.C., M.A.M., D.M.F.

## ACKNOWLEDGEMENTS

Sequencing data are available in the European Nucleotide Archive (ENA) under study accession no. PRJEB21034 (RNA-Seq), PRJEB21035 (ChIP-Seq). A pre-publication draft of the *E. coli*-to-Cancer Gene-function Atlas (ECGA) is available to reviewers prepublication at https://microbialphenotypes.org/wiki/index.php/Special:ECGA) at which the complete ECGA will be publically available after publication.

This paper is dedicated to the memory of our coauthor, colleague and friend Ken Scott whose scientific acumen, intensity, and generosity inspired us. We thank M Ellis, G Ira, H Dierick, V Lundblad, RS Harris, M Wang, H Zoghbi, and C Zong for helpful comments on the manuscript, M Yamada and T Nohmi for antiserum against Pol IV, and I Matic, SJ Sandler and JD Wang for bacterial strains. This work was supported by a National Institutes of Health (NIH) Director’s Pioneer Award DP1-CA174424 (SMR), a gift from the WM Keck Foundation (SMR, KMM), and the following grants: an NIH Director’s New Innovator Award DP20-OD008371 (CQ); R01-GM102679 (DB); NIH grant R01-GM088653 (CH); R01-GM089636 (JCH, DAS); R01-GM106373 (PJH); R35-GM122598 (SMR); Cancer Prevention and Research Institute of Texas (CPRIT) grants R1116 (KMM); RP170295 and RP170005 (CC); RP150578 (MAM); RP140553 (SMR, KMM); and RP160283 Baylor College of Medicine Comprehensive Cancer Training Program Postdoctoral Fellowship (DMF); and RP150578 (MAM); NIH U01-CA168394 (KLS); NIH/NCI R01CA175486, U24 CA209851, and CPRIT RP140462 to HL; the National Aeronautics and Space Administration through the NASA Astrobiology Institute under Cooperative Agreement No. NNA13AA91A issued through the Science Mission Directorate (PJH); the BCM Cytometry and Cell Sorting Core with funding from the NIH P30-AI036211, P30-CA125123, and S10-RR024574; and the BCM Integrated Microscopy Core with funding from the NIH HD007495, DK56338, and CA125123; the Dan L Duncan Comprehensive Cancer Center, and the John S. Dunn Gulf Coast Consortium for Chemical Genomics.

## STAR⋆METHODS

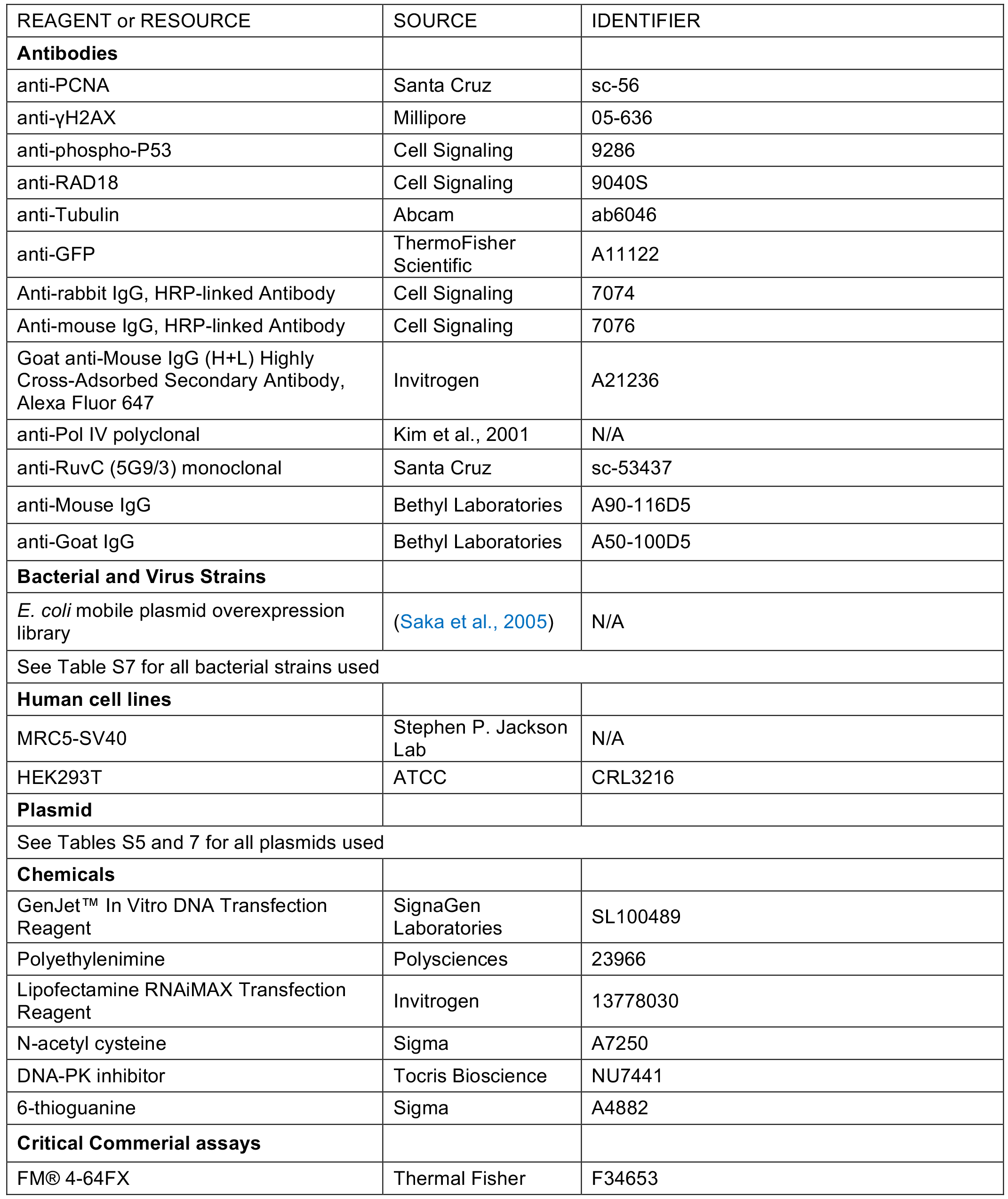
KEY RESOURCES TABLE

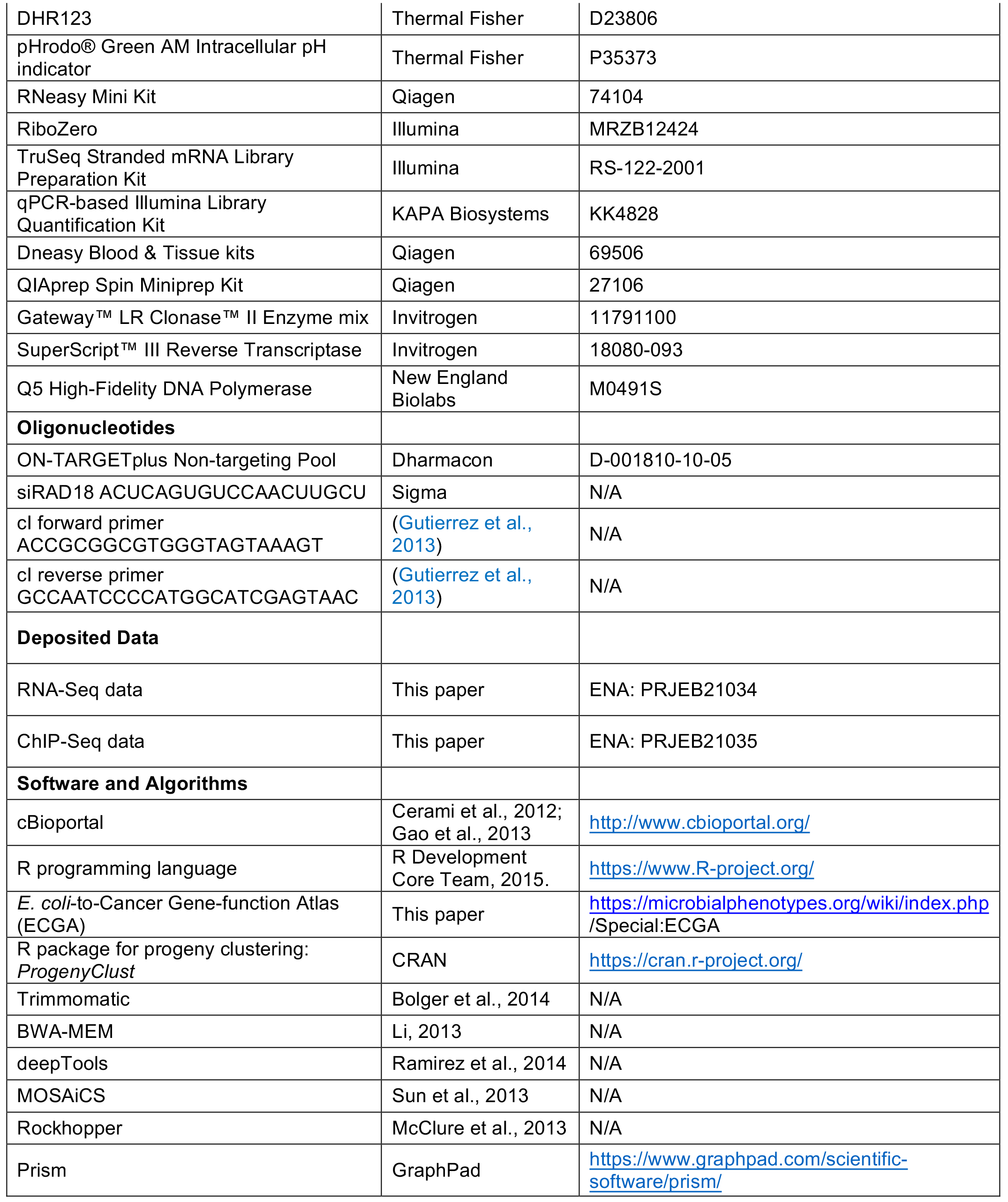

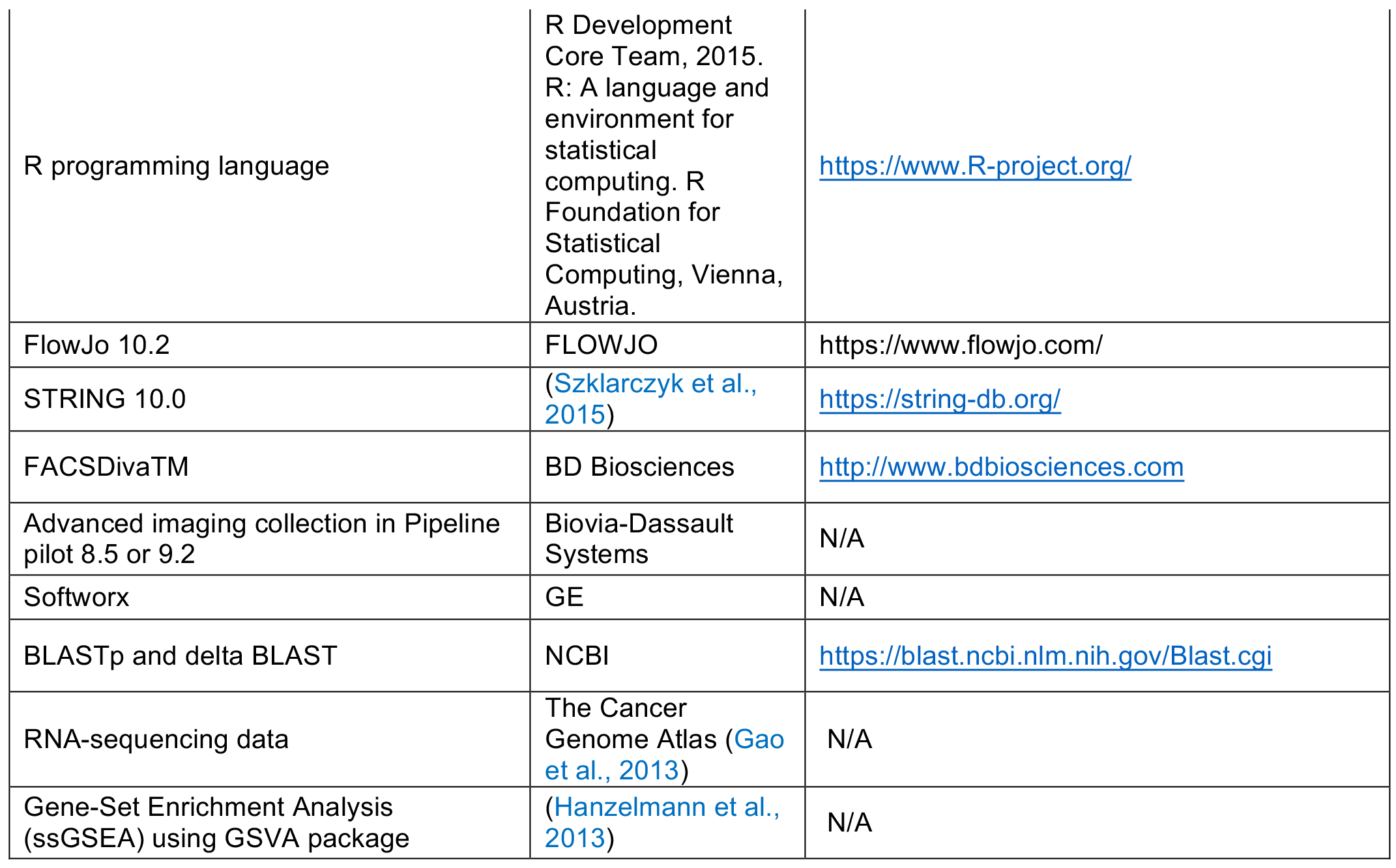

## CONTACT FOR REAGENT AND RESOURCE SHARING

Corresponding authors, S. M. Rosenberg and K. M. Miller are the contacts for reagents and resource sharing.

## EXPERIMENTAL MODEL AND SUBJECT DETAILS

*Escherichia coli* K-12 (strains MG1655 and W3110) and isogenic derivatives were used for all bacterial experiments. Human MRC5-SV40 and HEK293T cells were used for all human cell line experiments.

## METHOD DETAILS

### *Escherichia coli* strains and media

*E. coli* K12 strains and plasmids used in this work are shown in Table S7. Strains were grown in Luria Bertani Herskowitz (LBH) (Torkelson et al., 1997) rich medium or M9 minimal medium (Miller, 1993) supplemented with thiamine (10μg/ml) and 0.1% glucose or glycerol as a carbon source. Other additives were used at the following concentrations: ampicillin (100μg/ml), carbenecillin (20μg/ml), chloramphenicol (25μg/ml), kanamycin (30μg/ml), and sodium citrate (20 mM). P1 transductions were performed as described (Thomason et al., 2007). Genotypes were verified by antibiotic resistance, polymerase chain reaction (PCR) followed by sequencing, and, when relevant, UV sensitivity.

### Synthesis and generation of *E. coli* mutant and fusion genes

Mutant or truncated genes were synthesized to introduce site-specific mutations or small deletions in (GenScript) pUC57 backbone plasmids, and subsequently cloned into plasmid pNT3-SD to allow *E. coli* conjugation. Genes that encode wild-type and mutant DNA-binding transcription factors were fused with *mCherry* of (Shee et al., 2013) with a 4-6 alanine linker as described (Heckman and Pease, 2007). The plasmids mentioned above are shown in Table S7.

### *E. coli* mobile plasmid overexpression library

The mobile-plasmid collection is an ordered library of all 4229 *E. coli* protein-coding genes in a conjugation transferrable plasmid (Saka et al., 2005). Of these genes, 1017 (or 24%) encode the native *E. coli* protein, whereas 3212 (or 76%) encode the *E. coli* protein with an additional three N-terminal amino acids (Met-Arg-Ala) and an additional two C-terminal amino acids (Gly-Leu), with genes randomly distributed to one or the other kind. We found that native proteins were overrepresented significantly as positive for DNA damage in our screens (**RESULTS, A Larger Network Predicted**), implying that the 5 additional amino acids in some of the clones are more likely to confer false-negative results for the proteins that carry them than false-positive results. Table S1 shows the 208 validated DDP-gene clones and indicates the clones that produce native proteins with * next to the clone-ID number, and those with the five additional amino acids with an unmarked clone-ID number.

### Whole-genome primary DDP screen of ordered *E. coli* overexpression library

The ordered mobile-plasmid collection of 4229 *E. coli* genes in a conjugation transferrable plasmid (Saka et al., 2005) was mobilized into SOS-response-reporter strain SMR17962 (Nehring et al., 2016), to generate a DNA-damage screenable *E. coli* overproduction library. Protein overproduction is controlled by the IPTG-inducible P_*tac*_ promoter in cells that fluoresce red when they experience SOS-inducing DNA damage (single-stranded DNA) (Nehring et al., 2016). We adapted a high-throughput 96-well plate reader and robotics to screen for potential DDP-positive strains with increased mCherry fluorescence. Fluorescence intensity per unit of OD_600_ was compared in each well. Primary screens were performed on cells grown in M9 glucose or M9 glycerol medium (to survey two different conditions), each in duplicate. Ordered *E. coli* overproduction strains were grown to saturation overnight with shaking at 37°C in clear 96-well plates containing 150μl medium per well, then each well diluted 1:100 into 150μl IPTG-containing medium in 96-well plates (μ-clear, black, Greiner Bio-One, Monroe, NC, USA). The plates were shaken at 37°C for another 24h and analyzed in a Synergy 2 fluorescence plate reader (BioTek, Winooski, VT). We set thresholds of 20% (for glucose with IPTG induction) or 30% (for glycerol with IPTG induction) compared with the median fluorescence intensity per unit of OD_600_ of each individual 96-well plate to identify primary hits. Primary hits were called when two replicates done in the same medium were both above the threshold. Altogether, 414 candidate proteins were identified in this high-throughput plate-reader screen, then tested by flow cytometry for increased endogenous DNA-damage levels to eliminate false positives from the lower resolution/noisier plate-reader assay.

### *E. coli* flow-cytometry secondary screen for increased endogenous DNA damage

We screened candidate-protein hits from the primary (plate-reader) screen with our more sensitive flow-cytometric assays for SOS-induction/DNA damage (Pennington and Rosenberg, 2007). Each strain identified as positive from the primary plate-reader screen was grown at 37°C to saturation overnight in M9 glucose medium, diluted 1:100 into M9 glycerol medium, and grown for 9 hrs to early exponential phase at which time IPTG was added to 100μM to induce plasmid-protein overproduction. After 8 hours of induction, the cultures were diluted 1:100 in filtered M9 glycerol medium. Samples were analyzed in a LSR Fortessa flow cytometer (BD Biosciences) and analyzed with BD FACSDivaTM and FlowJo software. For these analyses, 10^5^ events were collected per strain, per experiment, with each strain assayed in three independent repeats. Student’s *t*-test (*p* value ≤ 0.05) and False Discovery Rate (FDR) q < 0.1 were calculated and applied based on Benjamini multiple comparison (Benjamini and Hochberg, 1995) to determine whether overproduction strains had significantly increased levels of endogenous DNA damage. Two-hundred and eight of the original 414 *E. coli* candidate proteins were validated as genuine DNA-damage-inducing DDPs when overproduced (shown Table S1).

### *E. coli* assay for RecA*GFP foci indicating single-stranded DNA

*E. coli* containing the chromosomal *recA4155gfp* allele, encoding RecA*GFP (Renzette et al., 2005), and the flow-cytometrically validated mobile-plasmid carriers were grown to saturation in M9 glucose medium at 37°C, then diluted 1:100 into M9 0.1% glycerol and grown for 9h to early log phase. IPTG was added to 100μM to induce protein overproduction for 8h as described above, then images taken and analyzed.

### *E. coli* assays for GamGFP and RDG foci

Saturated cultures of *E. coli* strains (GamGFP: SMR14334; RDG [RuvCDefGFP]: SMR19406) containing each of the 208 validated DDP-encoding mobile plasmids were grown and induced as described in flow-cytometric assays for DNA damage. 100ng/ml of doxycycline were added to induce GamGFP for 3h and RDG for 2h prior to harvesting. Cells were fixed with 1% paraformaldehyde for 15 min. and washed with PBS buffer three times before being concentrated for microscopy.

### *E. coli* microscopy and image analysis for RecA*GFP, GamGFP and RDG foci

Images were acquired using a 100x /NA = 1.4 immersion oil objective (Olympus) on a DeltaVision Elite deconvolution microscope (Applied Precision, GE). A z-series was acquired sampling every 0.2 microns for a total of 15-25 sections. The z-series was then deconvolved, and a maximum projection image rendered using Softworx (GE). Image analysis was performed using the Advanced Imaging collection in Pipeline Pilot 8.5 or 9.2 (Biovia-Dassault Systemes, San Diego). Projected images from the DeltaVision were read into Pipeline Pilot and metadata data parsed from the file name and path. A rolling ball background subtraction was applied to improve the signal-to-background ratio, and to facilitate further segmentation. Individual bacterial cells were then identified and segmented by applying a global threshold on images of the fluorescently labeled protein. Morphological manipulations (smoothing, opening and closing) were applied to refine the segmentation edges and a watershed was then performed to separate neighboring objects. Filtering was then applied to remove bacteria that fell outside a certain area threshold and that did not contain DNA. Foci were then identified using a more aggressive per-object background subtraction and peak identification method. Objects tentatively identified by this method were subsequently filtered by circularity, signal-to-background ratio, and size. Focus-positive bacteria were then determined using the co-localized objects component in the Advanced-imaging library in Pipeline Pilot. A binary metric, whether the cells were focus-positive or not, was calculated in addition to recording the total area and count of foci for those bacteria that were positive.

## STRING/network analyses

Known protein-protein interactions were displayed using CytoScape V3.4.0 software. Protein-protein interaction linkage scores were taken from the STRING 10.0 database (Szklarczyk et al., 2015) to identity interaction pairs. We used STRING, all parameters, with an interaction score cutoff of ≥0.6 (medium-to-high confidence). Random controls were produced by examining equal-size groups of random *E. coli* genes. *P* values were calculated with a hypergeometric test (Berkopec, 2007). The *E. coli* DDP network has network properties that are defined as scale-free and “small-world”, and it has significantly more edges (connectivity) compared with a random network (Figure 1G, S2A, Results). The human candidate-DDP network was generated similarly, and also has more connectivity than a random human-gene network or random human genes with *E. coli* homologs (Figures 2B, S2B, Results).

### *E. coli* forward-mutation assay

We used the forward-mutation assay of Matic and colleagues (Gutierrez et al., 2013) in which *E. coli* wild-type strain MG1655 harbors a chromosomal phage lambda *c*I transcriptional repressor gene, and a CI-repressible *tetA* gene, such that mutations that inactivate *c*I are scored as tetracycline-resistant (Tet^R^) mutant cfu. Into this strain, we conjugated 32 validated *E. coli* DDP genes in their mobile-plasmid-library vector (genes tested Table S1; Supplemental Discussion 3, and Table S1 for their mobile-library clone names). We developed a modified higher-throughput fluctuation-test assay for determining numbers of cultures with Tet^R^ mutants from which to calculate Tet^R^ mutation rates. Each DDP overproducer was grown overnight to saturation in M9 glucose with 20μg/ml carbenicillin at 37°C shaking, then diluted 1:10,000 into M9 glycerol carbenicillin and each culture split into 24 or 32 wells in 96-well-plates at 100μl per well. The plates were shaken at 37°C for 15h (early log phase), and IPTG added to attain 100μM in each well to induce protein overproduction for 8h, as described in flow-cytometric validation. From the end cultures, 5-10 μl were moved into LBH medium containing 10μg/ml tetracycline to determine the fraction of cultures that contained no Tet^R^ cells after incubation and scoring of the wells in the plate reader for OD (Tet^R^ cells) versus failure to grow (no Tet^R^ cells). The viable cell counts were estimated by sampling three wells chosen randomly. The P_0_ method was used to estimate mutation rates for each genotype as described with correction for the fraction sampled (Foster, 2006). The data reported (Figure 1J; Table S1) are the mean mutation rates (± SEM) of three experiments of at least 24 cultures per strain for each of the 32 strains assayed.

### *E. coli* Tet^R^ mutation verification by sequencing

We selected strains that overproduce the following 10 different DDPs with strong DNA-damage-up phenotypes: CsgD, TopB, CheA, YegL, MdtA, GrpE, HslU, YicR, UvrA, and Mrr. We selected 3-10 independent Tet^R^ mutant colonies, each from a separate culture from each strain, from which to sequence *c*I mutations. For the vector-only negative control, 19 independent Tet^R^ colonies were isolated. We amplified and sequenced a 1122nt region encompassing the *c*I gene as described (Gutierrez et al., 2013) to identify the mutations. For those Tet^R^ mutants that failed to yield PCR products, implying deletion of the *c*I gene, further outside primers (forward: ACCGCGGCGTGGGTAGTAAAGT, and reverse: GCCAATCCCCATGGCATCGAGTAAC) were used for PCR, and the products sequenced. In two cases (both TopB overproducers), whole-genome sequencing (WGS) was performed to determine the end-points for deletions that could not be determined via PCR and sequencing.

### *E. coli* whole-genome sequencing and analysis

Tet^R^ mutants were grown at 37°C to saturation overnight in LBH with 10μg/ml tetracycline, and genomic DNA was extracted and purified using DNeasy Blood & Tissue kits (Qiagen). Libraries were prepared using Nextera XT kits (Illumina); sequencing was performed on an Illumina Mi-Seq, and sequencing data analyzed as described (Xia et al., 2016). Sequencing reads were mapped to the MG1655 genome (NCBI RefSeq Accession: NC_000913.3). Low-quality reads and duplicates were removed. WGS files were visualized and deletion endpoints were analyzed using IGV software (Broad Institute, MA).

### Flow-cytometric assays for DNA loss

Quantification of anucleate cells by flow cytometry was adapted from (Joshi et al., 2013). Saturated cultures of *E. coli* strains derived from SMR21384 containing each of the 208 validated DDP-producing mobile plasmids were grown and induced as described in flow-cytometric assays for DNA damage. Cells were resuspended in 100 μl PBS, and stained with membrane dye FM^®^ 4-64FX (Thermal Fisher) with a final concentration of 10 μg/ml. The mix was kept on ice for 10 min. and then washed three times with PBS. A final concentration of 70% ethanol (pre-chilled) was used to fix the cells at −20°C for 1h, after which cells were washed with twice with PBS and resuspended in 100 μl PBS. 100 μl DAPI (5μg/μl) were used to stain DNA at room temperature (RT) for another 10 min. Samples were filtered and analyzed as described above.

### Flow-cytometric assay for intracellular ROS levels

Saturated cultures of *E. coli* strains derived from SMR21384 containing each of the 208 validated DDP-producing mobile plasmids were grown and induced as described in flow-cytometric assays for DNA damage. The ROS measurement protocol was modified from Gutierrez et al. (Gutierrez et al., 2013). In brief, cells were incubated with ROS-staining dye DHR123 (Invitrogen), which measures H_2_O_2_, for 30 min. at 4°C in M9 buffer. After washing twice with M9 buffer, flow cytometry analyses were performed immediately as described above.

### Flow-cytometric assay for intracellular pH

pHrodo^®^ Green AM Intracellular pH Indicator (Thermal Fisher) was used to measure intracellular pH in live *E. coli.* Protocols were adapted from (Loiselle and Casey, 2010). Cells were first washed with live-cell imaging solution (LCIS) and then 10 μl of pHrodo™ Green AM with 100 μl of PowerLoad™ concentrate were added to 10 ml of LCIS. The pHrodo™ AM/PowerLoad™/LCIS was mixed with cells and incubated at 37°C for 30 minutes. Cells were then washed twice with PBS to remove excess dye before flow-cytometric analysis. Intracellular pH calibration buffers (Thermal Fisher) were used as standards.

### Assays for sensitivity to DNA-damaging agents

Cultures of *E. coli* strain (SMR21384) containing each of the 208 validated DDP-producing mobile plasmids were grown as described in flow-cytometric assays for DNA damage with the following modifications: For hydrogen-peroxide (H_2_O_2_) treatment, 100 μM IPTG was used to induce overproduction of each DDP. Each culture was split into two tubes, prior to addition of 5mM H_2_O_2_ into one of the tubes for 15min. The cells with and without H_2_O_2_ were immediately diluted and plated onto LBH plates for assay of viable cells as cfu after incubation for a day at 37°C. For phleomycin or mitomycin C (MMC) treatment, saturated M9 glucose cultures were diluted into M9 glycerol medium with 100 μM IPTG to induce overproduction in 96-well plates. The plates were grown with shaking for 8 hours at 37°C to early log phase prior to addition of 1 μg/ml phleomycin or 0.05μg/ml MMC to each well. After 20 hours of continuous shaking, the OD600 was read using a BioTek microplate reader Synergy 2 (BioTek). DNA-damaging-agent sensitivities of the DDP-producing clones are normalized to sensitivity of vector-only controls: (treated/untreated DDP overproducer) / (treated/untreated vector-only) so that values < 1 indicate sensitivity. For all three assays for sensitivity to DNA-damaging agents, Student’s *t*-test (*p* value ≤ 0.05) with FDR adjustment (q ≤ 0.1) was used to determine whether DDP-overproducing strains were significantly more sensitive to DNA-damaging agents than the vector-only control.

### Clustering methods

For each DDP and DNA-damage outcome measure, raw data for each functional assay (overproduction versus vector) were converted into z scores and were used to delineate groupings of proteins with similar properties and patterns of response. Unsupervised discovery methods K-means in combination with Progeny Clustering (Hu et al., 2015) were performed using the R package *ProgenyClust* (Hu and Qutub, 2016) to determine the optimal number of protein clusters for the 208 DDPs. Seven functional tests were clustered by hierarchical clustering to assess the association of kinds, causes, and consequences of DNA damage.

### RNA-seq library preparation and sequencing

*E. coli* cultures were grown as described for flow-cytometric assays for DNA damage, and RNA was isolated from 1 ml of culture (~10^8^ cells) for each of two biological replicates. Total RNA was isolated using the RNeasy Mini Kit (Qiagen), according to the manufacturer’s protocol. RNAprotect Bacterial Reagent (Qiagen) was used to stabilize RNA during harvest and enzymatic cell lysis. After elution, total RNA was treated with RNase-free DNase I (NEB), according to the manufacturer’s protocol. RNA was recovered by phenol-chloroform extraction and ethanol precipitation. Ribosomal RNA was depleted using RiboZero (Epicentre/Illumina), according to the manufacturer’s protocol. Remaining RNA was concentrated by ethanol precipitation and approximately 100 ng of rRNA-depleted RNA was used to construct libraries using the TruSeq Stranded mRNA Library Preparation Kit (Illumina). Libraries were prepared according to the manufacturer’s protocol, using recommended modifications for previously isolated mRNA (McClure et al., 2013) (poly-A RNA enrichment steps excluded). Final RNA-seq libraries were run on a BioAnalyzer (Agilent) to estimate the average fragment size (~800 bp) and the concentration of adapter-ligated library fragments was determined using the qPCR-based Illumina Library Quantification Kit (KAPA Biosystems). Libraries were pooled and sequenced on an Illumina NextSeq 500 using a High Output v2 Kit (2 x 75 bp paired-end reads).

### Analysis and deposition of RNA-seq data

Read mapping, transcript assembly, and differential expression analysis were performed using Rockhopper (McClure et al., 2013), a bacteria-specific RNA-seq analysis pipeline, using MG1655 (NC_000913.3) as the reference genome. Genes were considered as differentially expressed if the fold change was greater than or equal to 2 and q-value was less than 0.01. Sequencing data are available in the European Nucleotide Archive (ENA) under study accession no. PRJEB21034.

### RDG ChIP-seq library preparation, sequencing, and data analysis

Cells were grown as for focus quantification, then crosslinked, lysed and sonicated as described (Xia et al., 2016). Immunoprecipitation and library preparation methods are based on those of (Bonocora and Wade, 2015) with small modifications as follows. RuvC antibody (Santa Cruz) was first pre-incubated with Dynabead protein A, then the RuvC-antibody-coated Dynabeads were incubated with cell lysates at 4°C overnight. Library preparation was performed while DNA fragments were still on Dynabeads. Samples were barcoded using NEBNext Multiplex Oligos for Illumina. Size selection of adaptor ligated DNA was performed on AMPure XP Beads as described in NEBNext ChIP-Seq Library Prep guidelines. Because the concentrations of eluted ChIP DNA are low, samples were amplified briefly prior to size selection, and a second amplification was performed after size selection. Sequencing was performed on an Illumina MiSeq. The pipeline for data analysis consists of the following steps: (i) reads were trimmed by Trimmomatic (Bolger et al., 2014) removing sequencing adaptors and low quality bases; (ii) reads were aligned by BWA-MEM (Li, 2013) to the W3110 genome [National Center for Biotechnology Information (NCBI) Reference Sequence (RefSeq) Database accession: NC_007779.1] and the plasmid pNT3 (Saka et al., 2005); (iii) Secondary alignment and multiple-mapped reads were discarded, this results in zero coverage in repetitive regions and regions present in both the genome and the plasmid, including the *csgD* gene; (iv) potential PCR duplicates were removed by Picard Tools MarkDuplicates; (v) bedGraph files were generated with deepTools (Ramirez et al., 2014) and imported to R for plotting; and (vi) peak calling was performed with MOSAiCS (Sun et al., 2013). Sequencing data are available in the European Nucleotide Archive (ENA) under study accession no. PRJEB21035.

### Western analyses of Pol IV protein levels

M9 glycerol cultures inducing wild-type, catalytically inactive, and β-binding-defective Pol IV were normalized to OD_600_ of 1.0, and 1 ml of each was pelleted, resuspended and boiled as described (Kim et al., 2001). Proteins were separated by 10% SDS-PAGE and transferred to PVDF membrane according to the manufacturer’s instructions (Amersham, GE Healthcare). The membranes were blocked with ECL Prime blocking agent (GE Healthcare) and probed with primary anti-Pol IV polyclonal antibody (Kim et al., 2001) (1:2000). The membrane was further probed with secondary polyclonal goat anti-rabbit IgG-Cy5 antibody (Bethyl Laboratories) and visualized by scanning in multicolor imager Typhoon detection system (GE Healthcare).

### Identification of human homologs using BLASTp and delta BLAST

“Homologs” are defined here as proteins with amino-acid similarity that could result from possible evolutionary relatedness. We used two basic local alignment search tools: the BLASTp and Delta-BLAST algorithms, searching protein sequences obtained from GenBank and other NCBI database resources. For both we used e-value < 0.01 (≤1 gene is identified by random chance in 100 queries) and sequence identity of ≥20%. Note that ≥20% sequence identity between *E. coli* and human is considerable. For example, known orthologs *E. coli* RecA and human RAD51 have 25% amino-acid identity. Given a protein query, BLASTp returns the most similar protein sequences from the protein database with e-value < 0.01 and identity ≥ 20%. Delta-BLAST uses multiple sequence alignment with conserved domains found in the CDD (Conserved Domains database from NCBI) and computes a Position Specific Score Matrix (PSSM) (Boratyn et al., 2012) with e-value < 0.01. Both methods were compared against the human protein database of NCBI. Proteins identified from either algorithm were identified as human homologs of the *E. coli* DDPs.

### Analyses of cancer survival and mutation loads

RNA-sequencing data from The Cancer Genome Atlas (TCGA) (Gao et al., 2013) were processed in the form of transcripts per million (TPM) as described (Rahman et al., 2015) and obtained via Gene Expression Omnibus (accession number GSE62944). Only the TCGA cancer types that had over 100 patients with RNA- and DNA-sequencing data were analyzed. Upon defining our gene sets of interest, RNA data were subjected to single sample Gene-Set Enrichment Analysis (ssGSEA) using GSVA package (Hanzelmann et al., 2013) in R. The resulting gene-set enrichment score for each sample was used as a representation of gene-set RNA level in each sample. Somatic mutation data for TCGA cancers were obtained in the form of mutation annotation files (raw or final) from the Broad Institute Genome Data Analysis Center (GDAC). For each sample, the sum of base-substitution and indel mutations was taken as the total mutation count, and log_e_ of this value was referred to as “mutation load.” Correlation analysis for “all human genes” was performed via bootstrapping. Briefly, we computed the mean correlation coefficient of mutation load with gene-set enrichment scores for 1000 randomly sampled gene sets, each consisting of a random number, between 10 to 1000, of genes out of over all human genes for which expression data were available. Kaplan Meier survival analysis was performed using “survival” package in R comparing the top and bottom tertiles of samples based on their gene-set enrichment score. Correlation analyses with mutation loads was performed in base R and correlation coefficients were plotted using the “corrplot” package in R.

### Cloning of human genes for DNA-damage analyses in human cells

Fifty-eight human DDP and 19 non-DDP cDNA clones (Table S5) in the Gateway entry vectors pDONR221 and pDONR223 (Invitrogen) were subcloned from an augmented library of ~32,000 Orfeome V8.1 (Yang et al., 2011) stated to contain sequenced human full-length cDNA clones, and additional full-length and commonest splice-variant length clones obtained from others including CCsBroad gene libraries. The size of cDNA from each gene was confirmed by restriction enzyme digestions. We also cloned, de novo, 15 candidate hDDP genes, one non-hDDP gene, tubulin and two *de novo* methylase genes (Table S5) that were not present as full-length clones in the Orfeome V8.1 (Yang et al., 2011) or CCsBroad gene libraries. These candidate genes were amplified from cDNAs generated from mRNAs extracted from the human cancer-cell lines U2OS or MRC5-SV40. PCR products of the correct size were cloned into the Gateway entry vector pENTR11 at restriction enzyme cut sites or into pDONR201 using *attB* site-specific recombination sites. Five DNMT1 truncated constructs were modified by using site-directed mutagenesis (Table S5). Clones were sequenced and verified as the correct gene sequence based on the Reference Sequence (RefSeq) database from NCBI. We subcloned each gene into a mammalian expression vector containing a GFP epitope tag (pcDNA6.2/N-EmGFP-DEST, Invitrogen), which allows us to analyze transfection efficiency and visualize protein localization in transfected cells. All human cell overexpression plasmids used in this study are listed in Table S5.

### Human cell lines, plasmids, and reagents

MRC5-SV40 and HEK293T cells were maintained in Dulbecco’s modified Eagle’s medium (DMEM) (Invitrogen) supplemented with 10% fetal bovine serum (FBS), 2 mM L-glutamine, 100 μg/mL penicillin, 100 μg/mL and streptomycin. Transient transfections into human cells were performed using GenJet (SignaGen Laboratories) for MRC5-SV40 and PEI (polyethylenimine, Sigma) for HEK293T. Transfections for siRNA were carried out with lipofectamine RNAiMax (Invitrogen) following the manufacturer’s instructions. The siRNAs were siNT: non-targeting pool (Dharmacon) and siRAD18: ACUCAGUGUCCAACUUGCU (Sigma). DNA-PK inhibitor (NU7441, Tocris Bioscience) was used at 2.5 μM 6 h prior to harvesting cells for flow cytometry. NAC (N-acetyl-cysteine, Sigma) treatment was performed twice, with a final concentration of 5 mM, post-24hr and -48hr transfection. To create inducible stable clones to verify DNMT1 and PCNA interaction, GFP-tubulin, GFP-DNMT1 and GFP-DNMT1-ΔPBD cDNAs were cloned into pcDNA5/FRT/TO/Intron vector (Invitrogen, CA). Inducible HEK293T FlpIn Trex GFP-tubulin, GFP-DNMT1 and GFP-DNMT1-ΔPBD cells were generated followed by manufacturer’s protocol and were cultured in the same normal medium with 15μg/ml Blasticidin and 80μg/ml hygromycin. Doxycycline (Sigma) was added to medium to trigger the production of GFP fusions.

### Human-cell DNA-damage screens by flow cytometry

We screened for increased DNA damage by flow-cytometric quantification of γH2AX- and phospho-P53-antibody signals among GFP-positive transfectants. Immunostaining was performed according to a standard procedure with minor modifications. Seventy-two hours post-transfection, cells were collected and approximately 1 × 10^6^ cells taken for staining. For staining, cells were fixed with 2% (v/v) formalin for 15 min on ice, washed twice in cold-PBS and permeabilized with 0.05% (v/v) Triton-X for 15 min on ice followed by two washes with PBS. The fixed cells were then blocked with 5% BSA-PBS for 1 hr, and stained with either γH2AX (Millipore) or phosphorylated p53 primary antibodies (Cell Signaling) overnight at 4°C. Cells were washed three times in 1% BSA-PBS followed by an incubation of Alexa Fluor 647 goat anti-mouse IgG in 5% BSA-PBS (Invitrogen) for 1 hr at room temperature in the dark, then washed three times with 1% BSA-PBS. Stained samples were measured by a BD LSRFortessa flow cytometer and analyzed using FlowJo software. Cells without transfection were used to set the threshold gating to determine the percentage of GFP- and γH2AX- or phosphorylated p53-positive cells, with 0.5% of control cells gated as the damage threshold. The DNA-damage ratio caused by protein overproduction is defined by (Q2/Q3)/(Q1/Q4), where Q2 is the number of transfected damage-positive cells; Q3 is the number of transfected damage-negative cells; Q1 is the number of untransfected damage-positive cells; and Q4 is the number of untransfected damage-negative cells. Results were obtained from at least two independent experiments. Statistical significance (*p* value) was determined using two-tailed unpaired Student’s *t*-test followed by false discovery rate (q value) correction. Both the γH2AX and phosphorylated-p53 assays show linear responses to exogenous DNA damage caused by ionizing radiation (Figures S3J and S3K), indicating their quantitative validity.

### HPRT mutagenesis assay

MRC5-SV40 cells were transfected with the plasmids indicated, and harvested 72 hours posttransfection. The percentage of GFP-positive cells of each transfectant was scored as transfection efficiency using a BD Accuri flow cytometer. The remaining cells were re-grown in 15 cm dishes for an additional 4 days. After a week of transfection, 3×10^6^ cells were plated in 15 cm dishes containing medium with 20 mM 6-thioguanine (Sigma), with five 15 cm dishes for each gene. In addition, 600 cells were plated in triplicate, per well, in a 6-well plate without 6-TG to determine plating efficiency. The plates were incubated at 37°C in a humidified incubator until colonies formed. The colonies were stained with 0.005% crystal violet. These colonies were counted, and mutation rates determined using the MSS-maximum likelihood estimator method with correction for transfection efficiency. We verified that 6-TG resistant clones result from *HPRT* mutations by sequencing the cloned *HPRT* cDNAs from four independent mutants (Supplemental Discussion 8). The *HPRT* cDNA is 657bp long, whereas *HPRT* including introns is 42kb, making sequencing the cDNAs more practical.

### Human-cell immunoprecipitation and western blot analysis

After induction of protein production using doxycycline in FlpIn-inducible HEK293T cells producing GFP-tubulin, GFP-DNMT1-WT or GFP-DNMT1-ΔPBD, cells were lysed with NETN buffer (150 mM NaCl, 1mM EDTA, 10 mM Tris-HCl, pH 8.0, and 0.5% NP-40) containing TurboNuclease (Accelagen) and 1 mM MgCl_2_ for 1 h at 4°C. Cell lysates were then centrifuged for 30 min at 4°C. GFP-tagged proteins were immunoprecipitated with 20 μl of GFP-Trap_A (Chromotek) for 1 h at 4°C. Beads were then washed three times with NETN buffer. Protein mixtures were eluted by boiling at 95°C with Laemmli buffer (4% (v/v) SDS, 20% (v/v) glycerol and 120 mM Tris-HCl, pH 6.8). For whole cell extracts, cells were collected with Laemmli buffer, and heated for 5 min at 95°C before loading. Samples were resolved by SDS-PAGE followed by western blot analysis. Primary antibodies were used as follows: anti-GFP (Invitrogen), anti-PCNA (Santa Cruz), anti-beta tubulin (Abcam), anti-RAD18 (Cell Signaling). Blots were analyzed by standard chemiluminescence (GE Healthcare, Amersham ECL Prime system) using a Bio-Rad molecular imager ChemiDoc XRS+ system.

### Statistics

All *E. coli* wet-bench experiments were performed at least three times independently, and a twotailed unpaired *t*-test was used to determine significant differences, unless otherwise specified. Error bars represent 1 SEM except where otherwise indicated. Pearson’s correlation coefficient was computed to assess the relationship between two parameters. STRING enrichment analysis was performed using hypergeometric tests with the correction for multiple comparisons. False discovery rate (FDR) adjustments are used to limit the overall type I errors in both *E. coli* and human DNA-damage flow-cytometry assays. The FDR (Benjamini Hochberg) method (Benjamini and Hochberg, 1995) is the default *p*-value adjustment method in this paper. Fisher exact test is used to determine whether two proportions are different. Wilcoxon rank-sum test was used to determine whether each gene has cancer-associated copy-number increases.

## QUANTIFICATION AND STATISTICAL ANALYSIS

Statistical details can be found in the main text, figure legends, or in the Method Details section.

## SUPPLEMENTAL INFORMATION

Supplemental information includes a discussion file, seven figures and seven tables that can be found with article online at ***

### Supplemental Discussion 1

#### Significant protein-protein interactions of random human homologs of *E. coli* proteins

Hypergeometric test analyses show that the 284 human homologs of *E. coli* DDPs have far more significant association (*p* = 1.2 × 10^−327^) than 284 random human proteins (*p* = 0.80), and 284 random human homologs of *E. coli* proteins (*p* = 1.8 × 10^−49^). Although not associated nearly as strongly as the human homologs of DDPs (1.2 × 10^−327^), the significant association of random human homologs of *E. coli* proteins (1.8 × 10^−49^) compared with random human proteins (*p* = 0.80) could potentially result if highly conserved proteins generally have more interactions with each other than random proteins. This might be because the most highly conserved, fundamental aspects of biology, and proteins that participate in them, are enriched for conserved protein machines (ribosomes, replisomes, transcription complexes, etc.), and/or fundamental pathways the actors in which have remained associated. Alternatively, it might be that proteins that function as part of interacting protein groups evolve more slowly, and so are overrepresented among conserved proteins.

### Supplemental Discussion 2

#### A larger network predicted and estimate of additional *E. coli* DDPs not discoverable in the mobile plasmid library

The 208 proteins are a large network, and occupy 5% of *E. coli* genes, but are likely to represent just over half of overproduction DDPs encoded in the *E. coli* genome. Per Figure S1E, we found that 1 of 99 random proteins not identified in the primary screen was positive in the sensitive flow-cytometry secondary assay, predicting an additional undiscovered 38 DDPs in the overproduction library used (Figure S1E). Further, although it is the most complete and least adulterated *E. coli* overexpression library, the mobile plasmid library (Saka et al., 2005]) contains some genes that encode five additional amino acids, which our data indicate were biased against in our screens. Twenty-four percent of clones in the library (STAR Methods) produce native *E. coli* proteins, and the rest produce proteins with three extra N-terminal (Met-Arg-Ala) and two extra C-terminal (Gly-Leu) amino acids, with the composition of genes in each class being random (Saka et al., 2005) (STAR Methods). We found that both the initial DDP candidates identified in the plate-reader primary screen and the 208 flow-cytometry-validated DDPs carried significantly higher fractions of native proteins than the library; there were 158 native proteins in the initial 414 candidates identified in the primary plate-reader screen (38%, differs from the library at *p* = 1.7 × 10^−11^, Fisher’s exact test), and 85 native proteins in the 208 validated DDPs, or 41% (shown in Table S1, differs from the library at *p* = 4.1 × 10^−8^, Fisher’s exact test). The data imply that some of the non-native proteins may have lost full function, and, because of that, gave false-negative readings in the screens. We found that the native genes in the library were “hit” in the primary screen at 16% (158 discovered out of 1015 native genes in the library), whereas the non-native genes were identified at 8% efficiency (256 discovered out of 3214 non-native genes in the library). If there are an additional 7.6% of the non-native proteins that would score as DNA-damage-promoting in our primary screen, if they did not carry the extra amino acids, then among the 3214 non-native-protein-encoding genes in the library, we predict that there would be an additional 244 overproduction DDP candidates found in the primary screen (7.6% of 3214). We found that candidates from the primary screen were validated in the secondary screen at 208 validated out of 414 candidates (Figure 1F; Tables S1 and S2), or just over 50%, which predicts 123 additional genuine DDPs among the predicted additional candidates.

### Supplemental Discussion 3

#### *E. coli* clones assayed for mutation rate

In Figure 1J, we assayed mutation rates in mutation-assay strains overproducing the following DDPs. DDPs that cause < 5-fold increase in DNA damage: DsbG, YijF, CadA, FolD, YddG, LeuO, UvrB, YajR, YbgQ, ORF 6106.1. DDPs that cause ≥ 5-fold increase in DNA damage: HypF, ZipA, YedA, CueO, YefU, MacB, HcaR, MdtB, SetB, DinD, RusA, YdcR, CsgD, HslU, SfsA, TopB, CorA, YegI, GrpE, PgrR, Mrr, MhpR. Non-DDPs: AceF, HprT, AceE, YaeG, YadF, PdhR, HrpB, MrcB, FhuD, YadG, Dgt, FhuA, HtrE, EcpD, FhuC, YacH, YadK.

### Supplemental Discussion 4

#### Cancer association of human proteins from a select DNA-damage screen

We analyzed published data from the limited human overexpression DNA-damage-up screen of (Lovejoy et al., 2009) by Fisher exact test against known (Forbes et al., 2015) and predicted (D’Antonio and Ciccarelli, 2013) cancer-driving genes. This overexpression screen of a set of nucleus/DNA-associated proteins discovered 96 human proteins (Lovejoy et al., 2009), which we found are overrepresented among known and predicted cancer drivers at *p* = 0.0001 and *p* = 0.0002, with DNA-repair proteins excluded (Fisher exact test, identities, Table S3). Only one protein was identified in common between the *E. coli* DDP homologs and the human overproduction screen (FIGNL1), indicating that the *E. coli* screen identified many new hDDP candidates, then validated hDDPs. Overall, the candidate hDDPs identified from human screens and the *E. coli* screen are highly significantly overrepresented among known (Forbes et al., 2015) and predicted (D’Antonio and Ciccarelli, 2013) cancer driver genes, independently of DNA-repair proteins, supporting the importance of DDPs to human cancer. We note that an unbiased screen of all human proteins for DNA damage on overproduction is not possible because the best human overexpression libraries contain a fraction of all human protein-coding genes, and many clones that are not full length. See STAR Methods, **Cloning of human genes for DNA-damage analyses in human cells.**

### Supplemental Discussion 5

#### Choice of candidate hDDP and control proteins for validation in DNA-damage assays

Of the 284 human homologs, we identified 121 candidates of particular interest according to the following criteria: (i) Many are encoded by genes amplified at high frequencies in cancer genomes from TCGA (Gao et al., 2013) (Table S4). (ii) For a minority, the genes are mutated or deleted at impressive, high frequencies in TCGA (Gao et al., 2013). (iii) Full-length clones that encode 90 of these appeared to be available in the Orfeome V8.1 or CCsBroad cDNA-clone collections (Yang et al., 2011). We determined by restriction mapping that many of the human genes in those libraries are not full length (Table S5), and cloned 18 genes including 15 candidate hDDP genes and three controls de novo as full-length cDNA clones that we sequence-verified (STAR Methods). We ultimately created 70 full-length overexpression GFP-fusion clones of human homologs of *E. coli* DDP genes, and overexpression GFP-fusion clones of 3 human homologs of *E. coli* damage-down genes, as possible negative controls (STAR Methods, Tables S5), 9 random human genes, and 11 random human homologs of *E. coli* non-DDP genes (Tables S5).

### Supplemental Discussion 6

#### Superiority of transient transfection to stable integration of genes encoding hDDP candidates

We found transient transfection to be superior to creation of stable clones because of apparent selection for mutations in the inducible hDDP candidate genes upon integration. Mutations in the hDDP candidates or other DNA damage-response pathways are selected probably because the gene products are toxic when overproduced and the genes are difficult to keep tightly “off”. The GFP-hDDP-gene fusions allow transient transfection assays to identify immediate effects of the DNA damage and to analyze only the minority population of cells that have been transfected successfully and produce the protein of interest. This cell subpopulation is GFP-positive, and easily identified in the flow-cytometric assays (e.g., Figure 3B).

### Supplemental Discussion 7

#### Estimation of additional human DDPs demonstrable in assays used here

We evaluated the validation efficiencies of four classes of human homologs of *E. coli* DDP genes (shown Figure 3D): genes that are—(i) both known (Forbes et al., 2015) or predicted (D’Antonio and Ciccarelli, 2013) cancer drivers and amplified in TCGA cancers; (ii) amplified in cancers and not known or predicted drivers; (iii) known/predicted cancer drivers that are not known to be amplified in cancers; and (iv) neither amplified in cancers nor previously known/predicted cancer drivers. Based on the number of candidates that we tested in each class among the 70 DDP homologs tested, these data correspond to the following validation rates as DNA-damage-promoting for each class: (i) 100%; (ii) 53%; (iii) 67%; and (iv) 27%. Based on the numbers of homologs not yet tested in each of these classes, our data predict that the following numbers of proteins among the remaining (284 − 70=214) human-homolog candidate hDDPs would be likely to be validated in these particular DNA-damage assays: (i) 6; (ii) 38; (iii) 34; and (iv) 7, for a total of at 85 more demonstrable hDDPs predicted among the 284-protein candidate hDDP network. We note, however, that the human-cell DNA-damage assays used favor detection of DNA doublestrand breaks, not all DNA-damage types comprehensively. Thus, many more of the human homologs may be DNA-damage promoting for other kinds of DNA damage than is estimated here.

### Supplemental Discussion 8

#### Verification of *HPRT* Mutations in 6-Thioguanine Resistant Human-Cell Clones

Four independent 6-thioguanine-resistant clones were shown to result from *HPRT* mutations by sequencing the cloned *HPRT* cDNAs. The mutations are: a single-basepair insertion between the 206-207nt of *HPRT* gene, and three identical deletions (from 403nt to 485nt). Two of the sequenced clones were independent DNMT1-overproducing transfectants, and two were from independent vector-only control transfected cells. The *HPRT* cDNA is 657bp long, whereas *HPRT* including introns is 42kb, making sequencing the cDNAs more practical.

### Supplemental Discussion 9

#### Controls

While analyzing RNA-Seq data, we identified a 2177bp deletion including the *lacl*^q^ region on the pNT3 empty vector. We have determined that this deletion does not alter results in any of our assays or any of our conclusions. The phenotypes of the truncated empty vector were compared with 10 non-DDP overproducers, and then with the full-length empty vector, in all 7 functional assays, and, by one-way ANOVA analysis, there were no significant differences between the means of all 11 strains in any of the 7 assays (*p*=0.19 GamGFP foci; *p*=0.28 RDG (reversed-fork) foci; *p*=0.99 ROS; *p*=0.99 anucleate cells/DNA loss; *p*=0.26 phleomycin sensitivity; *p*=0.08 H_2_O_2_ sensitivity; *p*=0.21 mitomycin C sensitivity), and no difference between it and the full-length vector (*p*=0.31; *p*=0.44; *p*=0.62; *p*=0.32; *p*=0.62; *p*=0.28; and *p*=0.78, respectively, two-tailed unpaired *t*-test).

### Supplemental Discussion 10

#### DNA-damage sensitivity not from mutations or *E. coli* DDP overproduction

We show that DNA-damage sensitivity does not result from heritable mutations in a sample of DDP-producing clones tested. Even highly sensitive DDP-producing strains were sensitive during overproduction, but not afterward, when colonies recovered after exposure were cultured and reexposed without DDP-gene induction (Figure S5H). The data imply that heritable mutations did not confer the DNA-damage sensitivities. Further, we used RNA-seq to quantify mRNAs of a panel of 32 *E. coli* DNA-repair genes, representing five DNA-repair mechanisms, in seven DDP-overproducing clones that display DNA-damage sensitivity (Figure S5F; Table S1), and that represent the six major biological DDP clusters (Figure 4N). The repair pathways represented were nucleotide excision repair (*uvrA*, *uvrB*, *uvrC*, *uvrD*), base-excision repair (BER: *mutT*, *mutM*, *xthA*, *nfo*, *ung*, *mug*, *nth*, *tag*, *alkA*, *nei*), mismatch repair (*mutS*, *mutL*, *uvrD*, *mutH*), homology-directed repair (*recA*, *radA*, *ruvA*, *ruvB*, *ruvC*, *recT*, *recF*, *recO*, *recR*, *recG*, *recN*), and homology-directed DSB repair (as for homology-directed repair with the addition of *recB*, *recC*, *recD*, and without *recFOR).* Thirty-one of the DNA-repair gene mRNAs were not downregulated with DDP overproduction, and 19 (*bamC*), 20 (*cusR*), 17 (*topA*), 4 (*aroP*), 8 (*hemX*), 14 (*yicR*) and 1 (*yqjD*) were significantly upregulated, presumably resulting from SOS or other DNA-damage or oxidative-stress responses (Figure S5F, Table S1). The sole DNA-repair-gene mRNA downregulated was that of *mutM*, which encodes the BER protein MutM, a DNA glycosylase that removes oxidized deoxy-guanine from DNA (Michaels et al., 1991), and which was decreased with DNA Topoisomerase (Topo I) overproduction (Figure S5F, Table S1). The data from all of the other genes show that reduced DNA-repair activities, inferred from DNA-damage sensitivities during overproduction of the seven representative DDPs (Figure S5F), did not result from transcriptional down-regulation of these DNA-repair genes. In the case of *mutM* mRNA reduction during Topo I overproduction, this is unlikely to cause the H_2_O_2_ sensitivity of Topo I overproducing cells because *mutM* null mutants are resistant to the low H_2_O_2_ levels we used (Asad et al., 2004). The data exclude the hypothesis that most or all of the DNA-damage sensitivities caused by overproduction of DDPs (Figures 4J-L and S5A-E; Table S1) result from transcriptional downregulation of DNA-repair genes. Potential post-transcriptional regulation mechanisms are not excluded. The data support the hypothesis that the DNA-damage sensitivities result from excess DNA damage overwhelming DNA-repair capacity, potentially titrating DNA-repair enzymes, or direct inhibition of DNA repair by the overproduced protein. Reduction of DNA-repair capacity could produce phenotypes like those of DNA-repair mutants, which would drive genome instability, in many DDP-gene dysregulated cells that do not possess DNA-repair-gene mutations.

### Supplemental Discussion 11

#### DNA damage reductions in some function clusters

Reduced ROS levels are apparent in a cluster in Figure 4M, as are reduced anucleate cells (DNA loss) in that figure. Similarly, clusters 3, 4, and 6 (Figure 4N) show reduced sensitivity (greater resistance than wild-type cells) to H_2_O_2_, and 2 and 3 (Figure 4N) show greater resistance to mitomycin C. A probable explanation for these “better-than-wild-type” phenotypes is that some imbalance in these cells may induce a stress response, one of the consequences of which may be improvement of the fidelity of, e.g., chromosome segregation (preventing anucleate cells), reduction of ROS levels, or resistance to DNA-damaging agents, as is documented, for example, for DNA-damage resistance induced by mild induction of the SOS response (Friedberg et al., 2005), among other stress responses.

### Supplemental Discussion 12

#### RDG ChIP-seq signals are enriched near known CsgD-binding sites, directionally

The RDG ChIP-seq experiments (Figures 5I and S7A-C) were performed twice, with peaks called against the matched control DNA-binding domain mutant, Csg*D*ΔDBD. We identified 155 total reproducible RDG ChIP-seq peaks with at least 2.5-fold increase in both repeats compared with CsgDΔDBD done in parallel. The following control simulations show that RDG ChIP-seq signals are enriched near known CsgD binding sites. We simulated CsgD-overproduction ChIP-Seq data by randomly distributing the called RDG peaks across the genome and analyzed, first, the number of binding sites with at least one RDG peak within 10kb, and, second, the distance between each known CsgD-binding site and the nearest RDG peak (median), for each of the 10 experimentally validated CsgD-binding sites (Brombacher et al., 2003; Dudin et al., 2014; Keseler et al., 2017; Ogasawara et al., 2011). Strikingly, there is at least one observed (actual) RDG peak within 10kb of 9 out of the 10 known CsgD-binding sites (Figure S7A-C), whereas there are only 5 known binding sites with at least one simulated RDG peak within 10kb of the 10 known binding sites in our simulations, which is significantly less than the observed (*p* = 0.01, two-tailed z-test). Also, the median distance between a given known CsgD-binding site and the nearest RDG ChIP-seq peak is 2.8kb for the real peaks, which is significantly closer than the 10.3kb median distance in our simulations (*p* = 0.009).

**CsgD-DBD-dependent stalled-fork RDG peaks not at known CsgD-binding sites.** For the 142 CsgD-DBD-dependent RDG peaks that are not significantly near to a known CsgD binding site, these could result from—(i) binding of CsgD to sites not yet identified in the literature, in which no comprehensive high-resolution (ChIP-seq) genome-wide binding study has been done; (ii) relaxation of the binding specificity of CsgD when overproduced such that sites not normally bound are bound on overproduction; and (ii) downstream (indirect) effects of regulation of CsgD-regulated genes by CsgD binding to its sites. For example, CsgD upregulation of other DNA-binding transcription factors could cause peaks at their binding sites, among other indirect but biologically real (CsgD-DBD-dependent) possibilities. The 142 of 155 RDG ChIP-seq peaks that are not near known CsgD binding sites do not overlap significantly with palindromic REP sequences, which are prone to form DNA cruciform structures (Stern et al., 1984) (four RDG peaks overlap with REPs, *p* =0.43, two-tailed z-test compared with median random distribution), or *terA-terC* regions, which have higher frequencies of fork convergence (11 RDG peaks overlap with *ter* sequences, *p* =0.47, two-tailed z-test, compared with compared with median random distribution).

**Upstream bias of RDG stalled-fork peaks at CsgD binding sites.** The 9 known CsgD-binding sites with one or more observed CsgD-DBD-dependent RDG peaks within 10kb are of two types: six show RDG peaks at or upstream of the binding sites in the replication path and five show RDG peaks downstream. The observed number of CsgD binding sites with upstream or colocalized RDG peaks is significantly higher than the median in the simulation (*p* = 0.04, two-tailed z-test), whereas the number of CsgD binding sites with RDG peaks downstream is not quite significant, at *p* = 0.13, two-tailed z-test. The analysis above suggests that direct binding of CsgD to its known binding sites underlies the RDG upstream and co-localized peaks, whereas the downstream peaks may reflect a component of direct CsgD-DNA-induced fork reversal, and a component of indirect effects, or binding to as yet unknown CsgD binding sites. One way that an RDG stalled-fork peak might result *downstream* in the replichore, but still via direct interaction of replication with bound CsgD, could be if some of the downstream stalled forks are caused by two events (illustrated Figure 5J, lower): first, slowing of replication forks by CsgD binding at its site, which might make the replisome susceptible to an otherwise surmounTable Second barrier of any kind downstream in the replication path.

### Supplemental Discussion 13

#### Model for DNA-damage promotion by the overproduced XanQ xanthine-proton symporter

Membrane transporters including proton (H^+^) symporters are overrepresented among DDPs that cause increased ROS when overproduced (Figures 6A-D), and cause DNA damage ROS-dependently (Figure 6B). The strongest generator of ROS-dependent DNA damage and ROS is the XanQ xanthine symporter. Although overproduction of each of three proton symporters decreases pH (increases H^+^, Figures 6F,G), their ROS promotion is not well correlated with decreased pH (Figure 6H), suggesting that the molecules they transport with H^+^ might underlie generation of ROS and DNA damage. For XanQ, overproduction of which causes high ROS but a moderate drop in pH (Figures 6F,G), one possibility (Figure 6I) is that the excess xanthine imported may be oxidized by ROS-generator xanthine oxidase (Kelley et al., 2010), causing the increased ROS, which damages DNA (Figure 6I). Other explanations are possible. Regardless of specific hypotheses, overproduction of membrane transporters generally may promote DNA damage, and for several, increased ROS, by any of many mechanisms that result from compromising compartmentalization of the cell from its environment.

**Figure S1.**
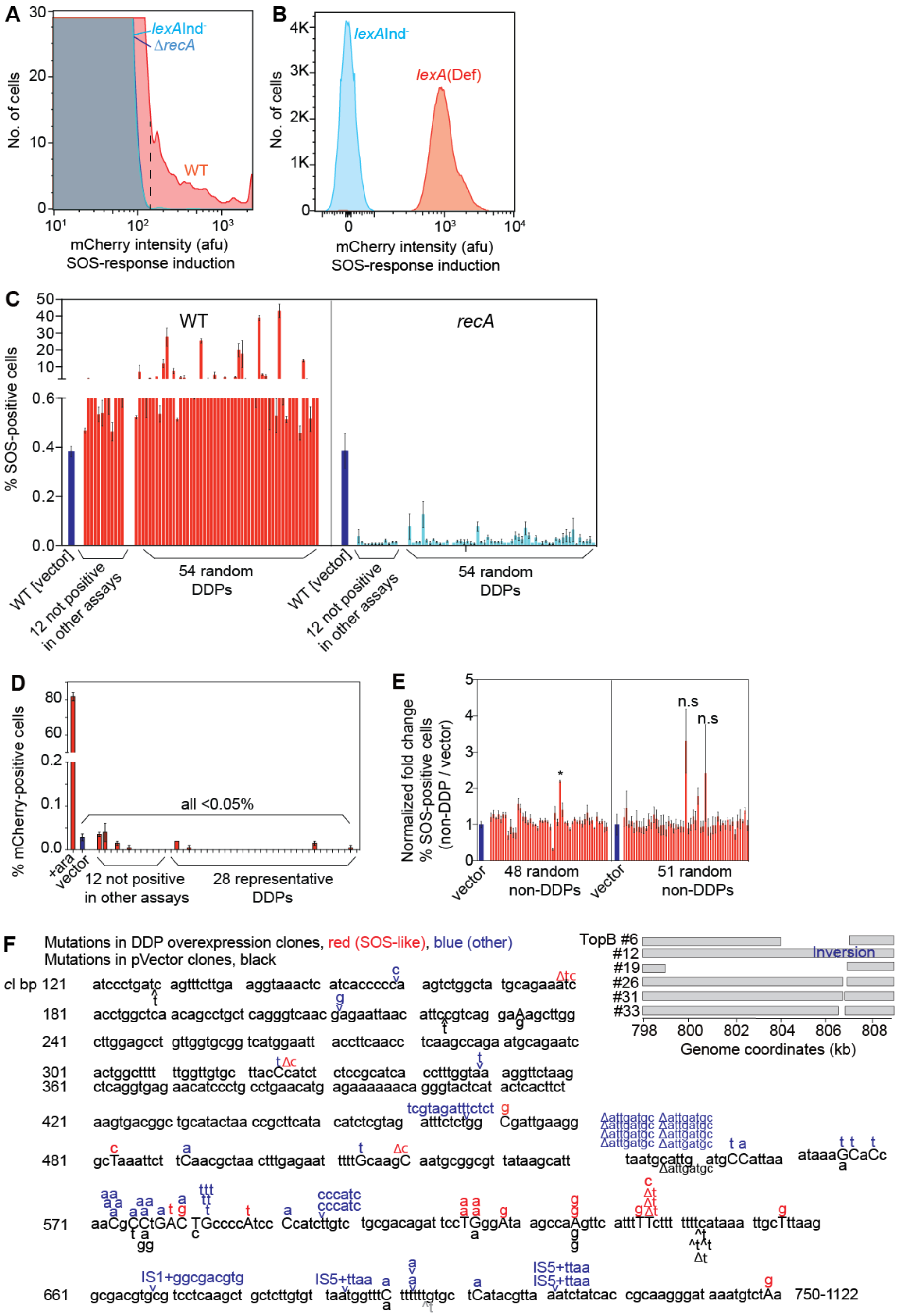
Genuine SOS Response, Detection of DNA Damage by the SOS-reporter Gene, and Evaluation of False Negatives in the Primary Screen. The *E. coli* DNA-damage assay detects fluorescence caused by upregulation of the SOS DNA-damage-response-activated promoter P*sulA* fused to mCherry in a non-genic chromosomal site (Nehring et al., 2016; Pennington and Rosenberg, 2007). This assay was shown to report on DNA damage, not spurious promoter firing with the demonstration that fluorescence induction requires the DNA-damage-sensing protein RecA, and is inhibited by a mutant SOS-response transcriptional repressor, LexAInd^−^, which does not de-repress the SOS genes during DNA damage (Pennington and Rosenberg, 2007), both also shown here (A). Further, of the spontaneous SOS-inducing DNA damage, previously 60% was shown to reflect DNA double-strand breaks (DSBs) and 40% single stranded DNA not at DSBs (Pennington and Rosenberg, 2007), and the spontaneous DSB frequency and rates per cell division and chromosome replication were confirmed in a direct, independent assay for DSB ends (Shee et al., 2013). Thus, this reporter reports on DNA damage. (A) Fluorescence requires the ability to activate the SOS response, and so is absent in SOS-induction-deficient Δ*recA* or *lexA3*(Ind^−^) mutant cells, demonstrating that DNA damage, not spurious promoter firing, underlies fluorescence increases, per (Pennington and Rosenberg, 2007). Using these negative controls, a flow-cytometry gate is set in each experiment, per (Pennington and Rosenberg, 2007), at fluorescence at which 10^−4^ of the *lexA3*(Ind^−^) negative-control cells are positive, shown as a vertical dashed line. The SOS-positive cells in the wild-type (WT) strain represent spontaneous DNA damage, per (Pennington and Rosenberg, 2007). Representative flow-cytometry histograms generated under the growth conditions used in the screens reported here. (B) Positive control, mCherry^+^ *lexA51*(Def) cells with constitutively activated SOS response, and negative control, *lexA3*(Ind^−^) SOS-off cells. (C) RecA-dependence of increased fluorescence in representative DDP clones. (D) Representative DDPs do not enhance mCherry fluorescence generally, when *mCherry* is controlled by non-SOS promoter *P*_*BAD*_. (E) Screen of 99 random *E. coli* proteins *not* identified in the primary plate-reader screen finds 1 SOS-positive clone. Because of its inherent noise level, the plate-reader primary screen is likely to generate some false-negative results; *i.e.*, it might miss clones with subtle but real DNA damage-up phenotypes. We therefore tested a sample of 99 random *E. coli* proteins that were not identified in the plate-reader primary screen for increased DNA-damage fluorescence in the sensitive flow-cytometry secondary assay, to estimate the frequency of possible false negatives. We found that 1 of the 99, or ~1%, was positive in the flow-cytometry assay. Thus, if the 3815 *E. coli* proteins that were not identified in the primary screen (4229 protein-coding genes in the library - 414 proteins found in the primary screen) also harbor 1% real but undiscovered DDPs, then an additional 38 overproduction DDPs are predicted to reside in the *E. coli* overproduction library, for an estimated total network size of 246 (38 + 208) DDP genes in the overexpression library used. In Supplementary Discussion 2, we estimate an additional 123 *E. coli* overexpression DDP genes that are likely to be in the *E. coli* genome, but would not be discoverable using the mobile plasmid overexpression library. (F) *E. coli* TetR clones from *c*I mutation assay harbor genuine mutations. Sequences of *c*I mutations from 10 different DDP-overproducing strains with strong damage-up phenotypes: CsgD, TopB, YegL, GrpE, HslU, Mrr, CheA, MdtA, YicR, and UvrA. The compiled sequences from the 10 DDP clones are shown in red and blue (n = 3-10 independent TetR mutants per clone). The 69 *c*I mutations sequenced include 3 IS (mobile)-element insertions, and 2 clones without a PCR product (both isolated from TopB overproducing cultures; PCR failure is due to large deletions, see STAR Methods). Mutations in the vector-only control strain are shown in black (19 independent mutants). Red font, mutations attributable to common errors of the SOS-upregulated error-prone DNA polymerases V and IV (Kobayashi et al., 2002; Maor-Shoshani et al., 2000; Wagner and Nohmi, 2000); blue font, mutations not attributable to common Pol V or Pol IV errors. Capital letters, basepairs that were changed or deleted; carrots, insertion points of bases indicated; Δs, deletions of the bases indicated. Upper right, diagram of gross chromosomal rearrangements (GCRs) found in TopB-overproducing cells, among which, 18% (7/40) of the TetR mutations are GCRs, including large deletions (from 200bp-6200bp), an inversion, and a transposon insertion.

**Figure S2.**
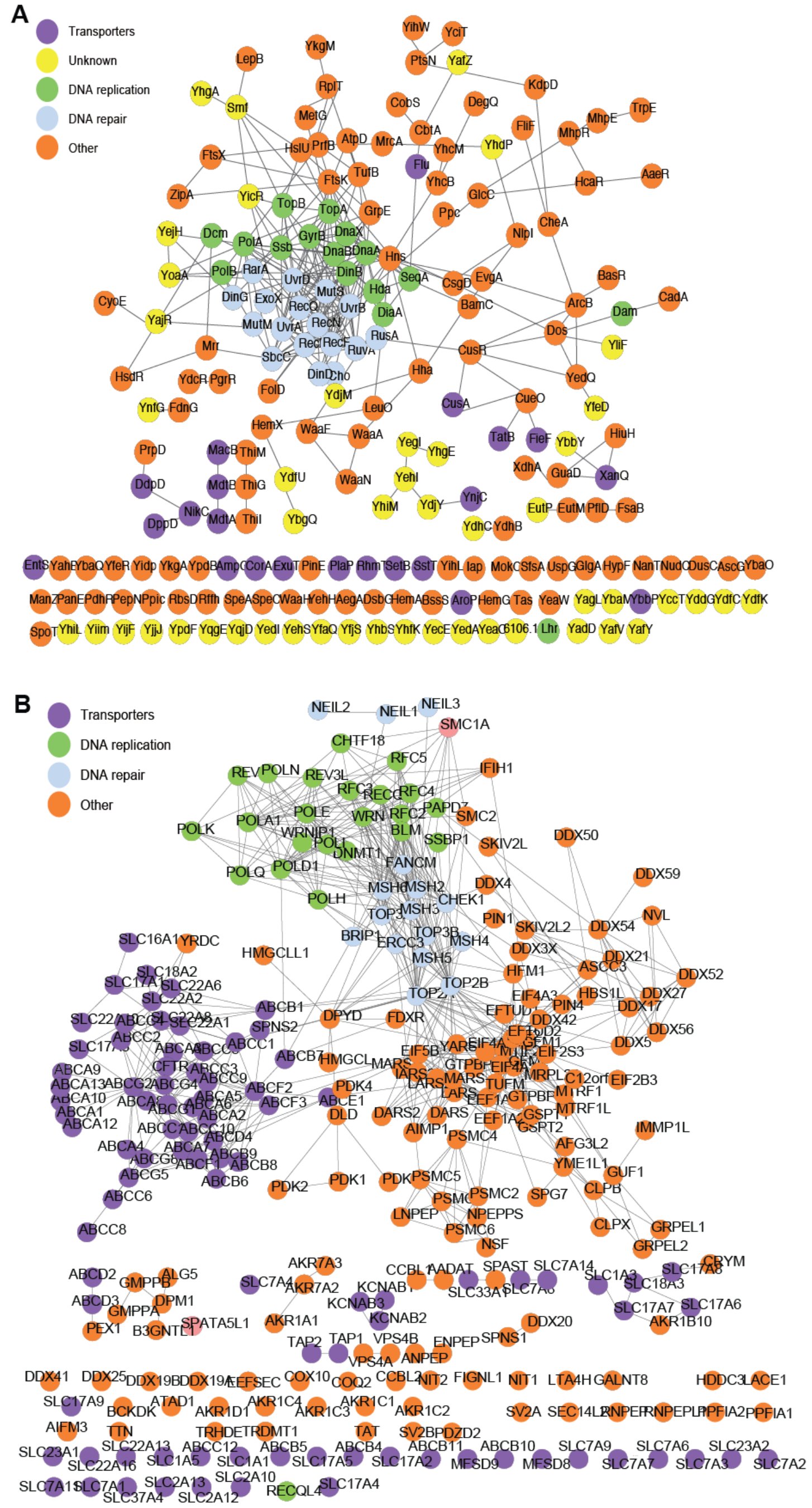
Specific Protein-Protein Interactions in *E. coli* and Human DDP Networks. (A) Specific protein-protein interactions in *E. coli* DDP network. Diagram details are as indicated in Figure 1G. Image enlarged to illustrate the specific proteins with interactions. Connectivity in random protein networks is discussed in Supplemental Discussion 1. (B) Specific protein-protein interactions of human candidate DDP network. Diagram details are as indicated in Figures 1G and 2B. Image enlarged to illustrate the specific proteins with interactions, similarly to the human candidate DDP network in Figure 2B. Although, the human homolog (candidate hDDP) network has DNA-repair and -replication genes at its center, removal of these proteins leaves the remainder with still significant connectivity: *p* = 4.1 × 10^−215^ (hypergeometric test), though less than the whole network (*p* = 1.2 × 10^−327^, hypergeometric test). Random human proteins do not form robust protein-protein association networks and the significant protein-protein interactions of well conserved proteins—the human homologs of random *E. coli* proteins—are discussed Supplemental Discussion 1.

**Figure S3.**
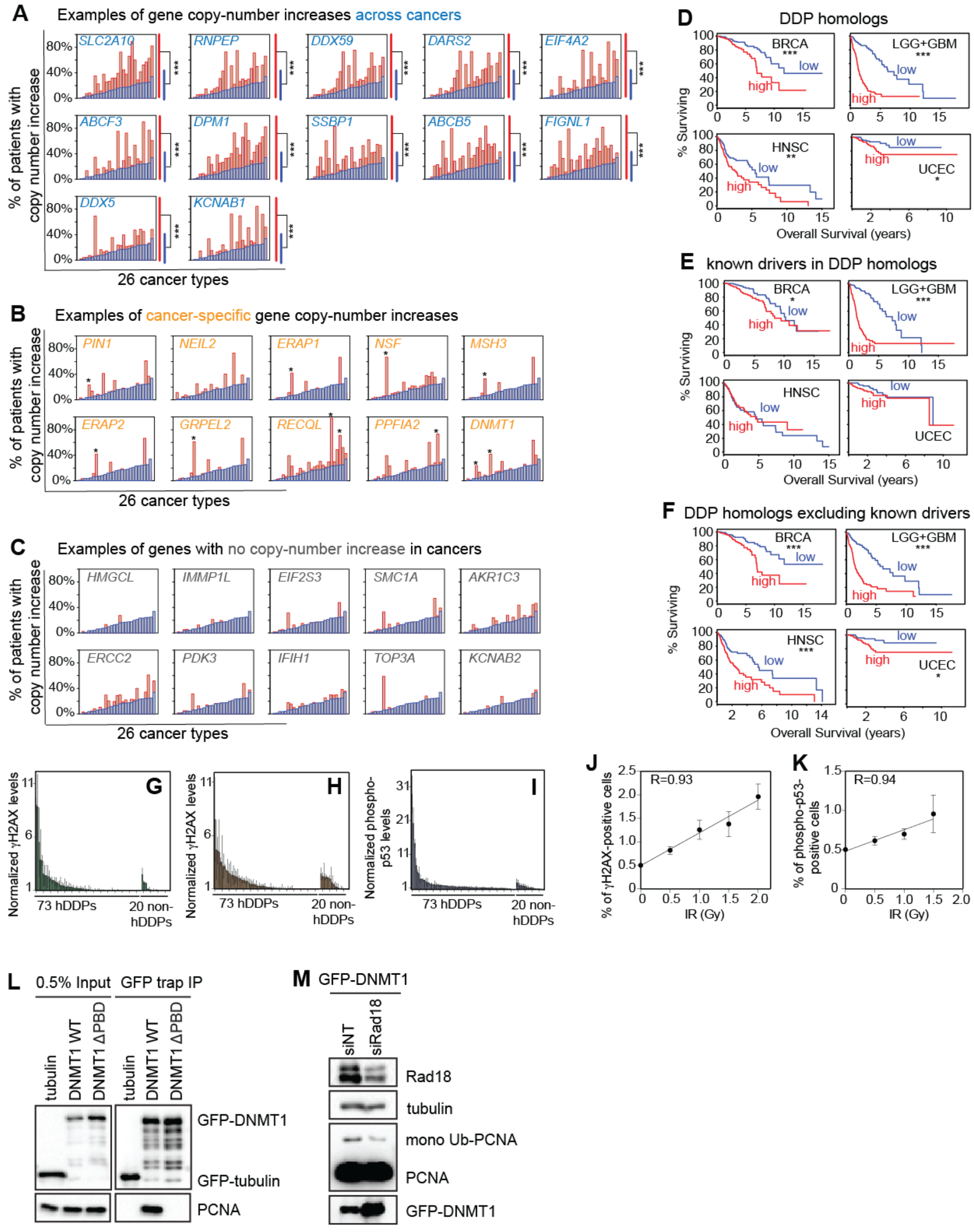
Association of Human Homolog Network with Cancers: Copy-number Gain, Survival, and Mutation Load, and Controls for Human DNA-Damage Assays and DNMT1. (A to C) Twenty-six cancer types in TCGA data (Gao et al., 2013) are displayed along the x axes. Blue, median % of patient cancers with increased copy number of any gene in their genome; red, % of patient cancers with increased copy number of the gene indicated. Copy-number-increases in human homologs of *E. coli* DDP genes are higher than those of human homologs of random *E. coli* genes (*p* < 0.05, FDR < 0.10, Wilcoxon test). Examples of the Pan-Cancer copy-number-increase analysis (GISTIC threshold copy-number gain ≥ 1) of the 284 human homologs of *E coli* DDP genes in 26 cancer types are shown here (complete analysis Table S4). The human genes fell into three categories: (A) genes with increased copy numbers across cancers (fold change > 1.5, *p* < 0.05, FDR < 0.10, Wilcoxon test); (B) genes with cancer-specific copy-number-increases (*p* < 0.05); and (C) not particularly cancer associated. (D) Decreased cancer survival is associated with high DDP-homolog RNA levels in cancers [our analyses of data from TCGA (Gao et al., 2013), STAR Methods]. BRCA, breast invasive carcinoma; LGG+GBM, gliomas (low-grade glioma + glioblastoma multiforme); HNSC, head and neck squamous cell carcinoma; UCEC, uterine corpus endometrial carcinoma. *, **, *** indicate that survival of the cancers with high and low levels of the 284 RNAs differ at *p* ≤ 0.05; ≤ 0.01, and ≤ 0.001 respectively, log-rank test. (E) Decreased cancer survival is not confined to the previously known cancer drivers in the network. RNAs of the known (Forbes et al., 2015) and predicted (D’Antonio and Ciccarelli, 2013) cancer-driving genes among the homologs show less association with poor survival than the whole homolog network. (F) RNAs of the human DDP homologs excluding known (Forbes et al., 2015) and predicted (D’Antonio and Ciccarelli, 2013) drivers are still associated with decreased patient survival, indicating that genes in the homolog network other than and in addition to the previously known drivers are also associated with decreased survival. (G-I) Human MRC5-SV40 or HEK293T were transfected with human candidate DDP and random human genes and random human non-DDP homologs of *E. coli* genes, and DNA-damage markers were analyzed by flow cytometry. (G) Assay I: γH2AX in MRC5-SV40 cells. (H) Assay II: γH2AX in MRC5-SV40 cells treated with DNA-PK inhibitor, which inhibits DNA break repair by non-homologous end joining and so is a sensitizing DNA-damage screen. (I) Assay III: phosphorylated p53 in HEK293T cells. Data represent mean ± range, n ≥ 2. The candidate hDDP genes differ from the 20 random human genes, *p* < 0.0001, Fisher exact test. (J and K) Linear response of both human-cell DNA-damage-detection assays with exogenous ionizing radiation treatment indicates quantitative validity of these assays. Percent of MRC5-SV40 cells that are positive for (J) γH2AX, and (K) phosphorylated p53 (p53-S15p), in flow-cytometric assays. MRC5-SV40 cells were treated with IR with the indicated dose and analyzed by flow cytometry. (L) DNMT1 wild-type (WT) but not ΔPBD interacts with replisome sliding clamp, PCNA. The tubulin negative control also does not interact with PCNA. GFP-trap immunoprecipitation was performed in flipIn stable inducible HEK293T cells expressing GFP-tubulin, GFP-DNMT1-WT or GFP-DNMT1-ΔPBD. Interactions were determined in immunoprecipitation samples by western blotting with anti-GFP and -PCNA antibodies. (M) Overproduction of wild-type DNMT1, but not the DNMT1-PBD-defective mutant protein, enhances RAD18-mediated ubiquitylation of replisome clamp, PCNA. Knockdown of *RAD18* reduced the level of ubiquitylated PCNA in DNMT-WT overexpressing cells. Western analyses of PCNA ubiquitylation in MRC5-SV40 cells transfected with the plasmids indicated in combination with non-targeting (NT) or *RAD18* siRNA.

**Figure S4.**
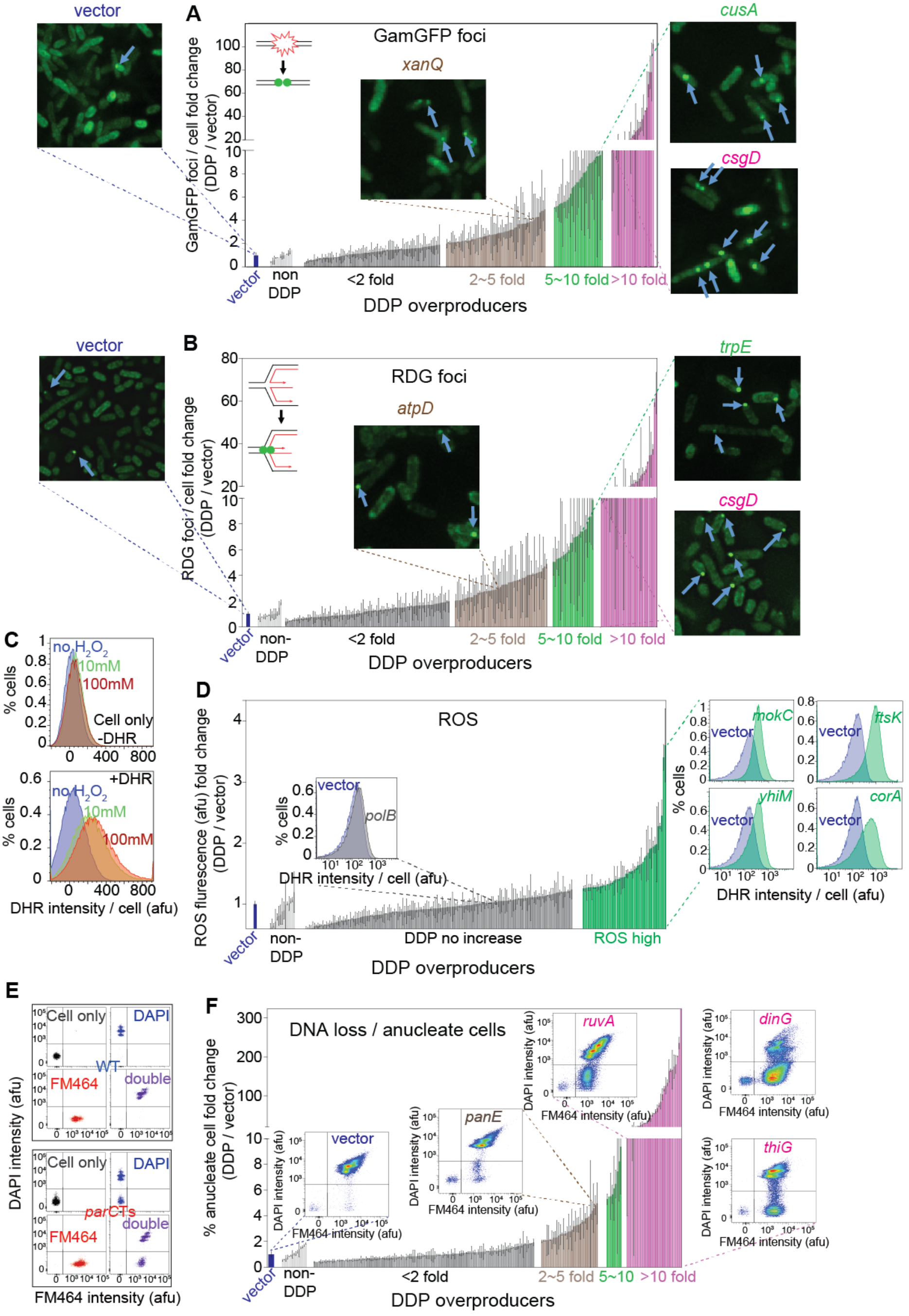
DSBs, Reversed Forks, Reactive Oxygen, and DNA Loss in DDP Clones. (A) Increased GamGFP foci indicate DSBs caused by overproduction of 87 of the 208 *E. coli* DDPs. DSBs in all 208 *E. coli* DDP-overproducing clones were visualized and quantified as GamGFP foci (Shee et al., 2013) using automated microscopy. 87 of the 208 clones, were significantly different from the vector-only control at *p* < 0.05, q <0.10 (unpaired two-tailed *t*-test with FDR adjustment). Each bar represents a DDP clone (mean ± range, n=2 experiments of >1000 cells per strain, STAR Methods). DDP-overproducing clones are grouped by the fold change of GamGFP focus levels compared with the vector-only strain. Blue, vector only; grey, < 2-fold increase; brown, 2-5-fold increase; green, 5-10-fold increase; magenta, > 10-fold increase, per Table S1 (data summary). Representative images are shown above and to the right of the bar graphs. None of 25 non-DDP overproducers had increased GamGFP foci, showing that clones with high levels of GamGFP foci are enriched in the DDP-overproducing clones (*p* = 3.6×10^−6^, one-way Fisher’s exact test). (B) Stalled reversed replication forks in most DDP-overproducing strains. Significantly increased reversed forks (RFs), visualized as RDG foci in Δ*recA* cells (Xia et al., 2016) via automated microscopy, occur in 106 of the 208 (51% of) *E. coli* DDP-overproducing clones. The 106 were significantly different from the vector-only control, *p* < 0.05, q <0.10 unpaired two-tailed *t*-test with FDR adjustment. Each bar represents a DDP clone (mean ± range, n=2 experiments of >1000 cells per strain). Blue, vector only; grey, < 2-fold increase; brown, 2-5-fold increase; green, 5-10fold increase; magenta, >10-fold increase, per Table S1 (complete data summary). Representative images shown above and on the right side of the bar graphs. None of the 30 non-DDP overproducers had increased RDG foci; so, DDP overproducers with high RDG-focus (RF) loads are enriched in the DDP-overproducing strains (*p* =4.0×10^−9^, one-way Fisher’s exact test). (C-D) Increased intracellular ROS levels in 56 *E. coli* DDP-overproducing clones identified by a flow-cytometric assay after di-hydrorhodamine (DHR) staining. Intracellular ROS levels in all 208 *E. coli* DDP-overproducing clones were measured with the peroxide-specific dye DHR (Gutierrez et al., 2013) and fluorescence was measured by flow cytometry. We found that 56 of the 208 (27%) were significantly different from the vector-only control, *p* < 0.05, q <0.10 unpaired two-tail *t*-test with FDR adjustment. Each bar represents a DDP clone (mean ± range, of two experiments). Blue, vector only; green, DDP overproducers with increased ROS levels, per Table S1. Representative flow cytometry histograms are shown above the bar graphs. None of 17 non-DDP overproducers had increased ROS, such that high-ROS clones are enriched in the DDP-overproducing clones at *p* = 0.006 (one-way Fisher’s exact test). (E-F) DNA loss in 67 *E. coli* DDP-overproducing clones identified by flow-cytometric quantification of anucleate cells. (E) Increased anucleate cells can be detected in a *parC*TS mutant, which has a severe chromosome-segregation defect at non-permissive temperature. Live cells were stained with FM464 membrane-staining dye, and then fixed and stained with DAPI for DNA staining. Cells with positive membrane but negative DAPI staining are anucleate cells (lower right quadrant). (F) We found that 67 (32%) were significantly different from the vector-only control, *p* < 0.05, q <0.10 unpaired two-tail *t*-test with FDR adjustment. Each bar represents a DDP-overproducing clone (mean ± range, n=2). DDP overproducers are grouped by the fold change of anucleate cells levels compared with the vector only strain. Blue, vector only; grey, < 2-fold increase; brown, 25-fold increase; green, 5-10-fold increase; magenta, >10-fold increase in anucleate cells, per Table S1. Representative flow cytometric histograms are shown above and to the right of the bar graphs. Only 1 of 16 non-DDP-overproducing clones had increased anucleate cells, such that clones with high-levels of anucleate cells are enriched in the DDP-overproducing strains at *p* = 0.02 (one-way Fisher’s exact test).

**Figure S5.**
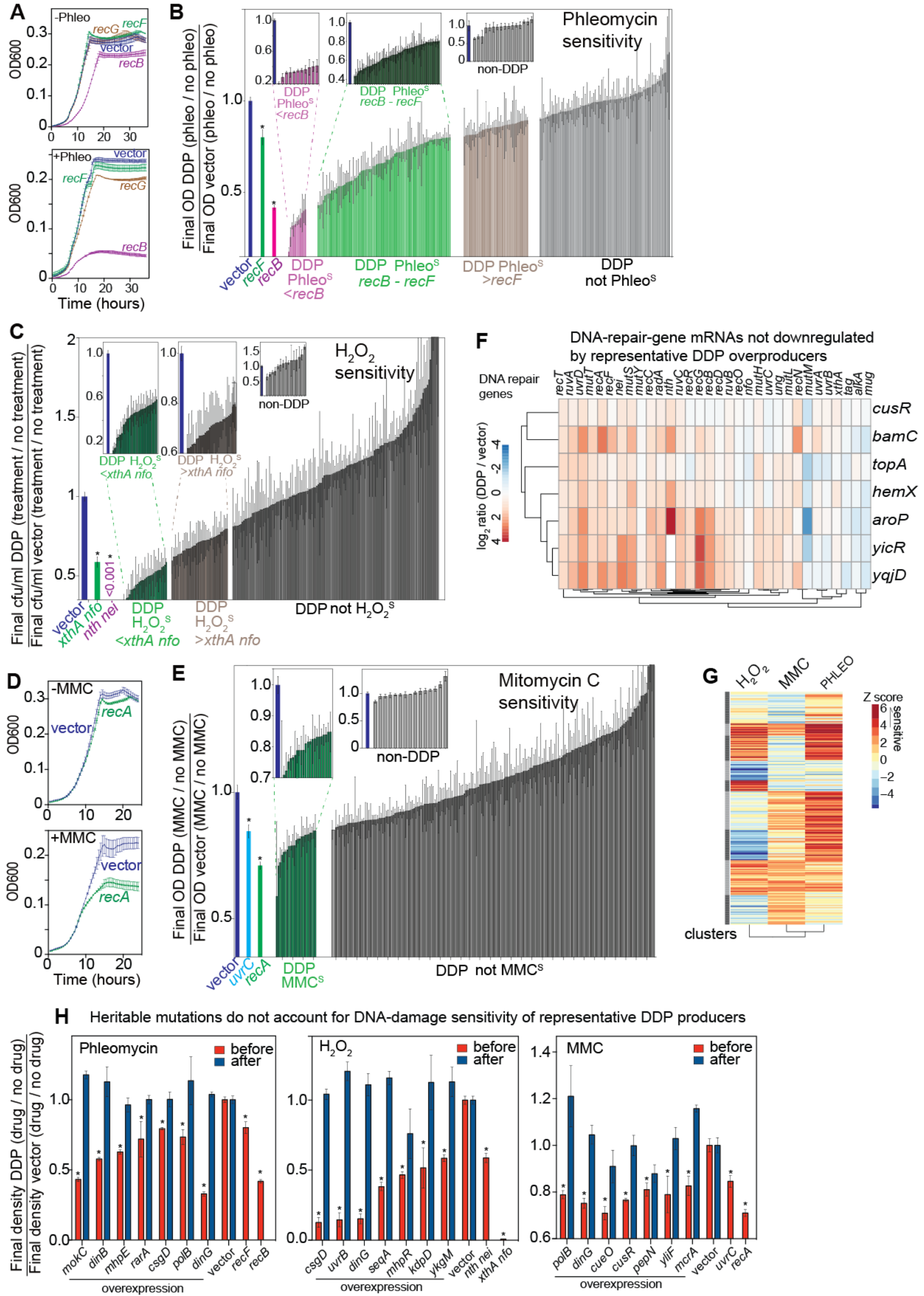
Sensitivities to DNA-Damaging Agents. (A) Phleomycin sensitivity detected by growth inhibition in known mutants with reduced homologous recombinational (HR) repair efficiency. *recF*: defective in single-strand gap HR-repair; *recG*: reduced ability to branch migrate Holliday junctions; *recB*: defective in DSB repair by HR (Kuzminov, 2011). (B) 106 *E. coli* DDP overproducers are sensitive to phleomycin. Phleomycin sensitivities of the DDP-overproducing clones are shown as normalized to sensitivity of vector-only controls: (treated/untreated DDP overproducer) / (treated/untreated vector-only) so that values < 1 indicate sensitivity. Among the 106 sensitive DDP clones, 11 are more sensitive than a *recB* mutant; 72 are within the range of *recB* and *recF* mutants; 23 are more sensitive than the vector-only control but less than *recF* mutant. One of the 16 non-DDP-overproducing strains has phleomycin sensitivity, such that overproducing clones with phleomycin sensitivity are enriched in the DDP-overproducing clones at *p* = 0.0003 (one-way Fisher’s exact test). (C) Sensitivity of 75 *E. coli* DDP-overproducing clones to oxidative-damage-inducing agent H_2_O_2_. H_2_O_2_ sensitivity was detected by reduced viability, measured by colony forming units in known mutants with reduced base excision repair (BER) efficiency. *xthA* (inset) encodes exonuclease III; *nfo* encodes endonuclease IV. *xthA nfo* (green) double mutants are reported and confirmed by us to be more sensitive to H_2_O_2_ than each single mutant (Galhardo et al., 2000). *nth* encodes endonuclease III; *nei* encodes endonuclease VIII. The *nth nei* (purple) double mutant has almost no AP lyase activity, and so has extreme sensitivity to H_2_O_2_ (Saito et al., 1997), consistent with our observation.. In our assay, H_2_O_2_ sensitivities of the DDP-overproducing clones are shown as normalized to sensitivity of vector-only controls: (treated/untreated DDP overproducer) / (treated/untreated vector-only) so that values < 1 indicate sensitivity. Among the 75 H_2_O_2_-sensitive DDP overproducers, 36 were more sensitive than the *xthA nfo* mutant; 39 were more sensitive than the vector-only control but less sensitive than the *xthA nfo* mutant. None of the 15 non-DDP-overproducers had H_2_O_2_ sensitivity, such that overproducing clones with H_2_O_2_ sensitivity are enriched in the DDP-overproducing clones at *p* = 0.002 (one-way Fisher’s exact test). (D and E) Sensitivity of 10 *E. coli* DDP-overproducing clones to interstrand-crosslinking agent Mitomycin C (MMC). (D) MMC sensitivity was detected by growth inhibition in known mutants with defective HR repair or nucleotide excision repair (NER): *recA*, defective in HR-repair and SOS response; and *uvrC*, defective in NER (shown in E, cyan). (E) Ten *E. coli* DDP-overproducing clones were sensitive to MMC. Sensitivities of the DDP-overproducing clones are normalized to sensitivity of vector-only controls: (treated/untreated DDP overproducer) / (treated/untreated vector-only) so that values < 1 indicate sensitivity. None of the 16 non-DDP overproducers was MMC sensitive. Overproduction clones with MMC sensitivity are not enriched among the DDP-overproducing clones at *p* = 0.47 (one-way Fisher’s exact test). (F) Representative DDPs do not downregulate RNAs of DNA-repair genes upon overproduction. We chose 7 DDPs that confer DNA-damage sensitivity on overproduction representative of the six DDP function clusters (Figure 4N; Table S1). We assayed RNA levels of a panel of 32 DNA-repair genes by RNA-seq with and without overproduction of each of 7 DDPs that confer sensitivity to the following agents: TopA and BamC (H_2_O_2_); CusR, HemX, AroP, YicR, and YqjD (phleomycin). Log_2_ ratio (DDP overproducer / vector) for each DNA-repair gene in each of the seven DDP-overproducing strains was calculated and clustered by hierarchy clustering (Johnson, 1967). Most of the DNA-repair genes are up-regulated during DDP overproduction, with the exception of *mutM* RNA, which is downregulated in the TopA-overproducing strain. However, *mutM* downregulation is unlikely to account for the H_2_O_2_ sensitivity of the TopA-overproducing strain because even *mutM* null mutants are not sensitive to H_2_O_2_ (Asad et al., 2004). (G) Cluster analysis of quantitative data on DNA-damaging-agent sensitivities in the *E. coli* DDP network. Each bar represents a quantitative phenotype of each strain producing each of the 208 *E. coli* DDPs, arrayed along the x axis. Assays indicated to right of the heatmap: H_2_O_2_, hydrogen-peroxide sensitivity (reduced base excision repair); PHLEO, phleomycin sensitivity (reduced DSB-repair); MMC, mitomycin-C sensitivity (reduced NER and/or HR repair). Red bar: high Z score, increased DNA-damage sensitivity of DDP-overproducing clones compared with the vector-only negative control. Heritable mutations do not account for DNA-damage sensitivities of seven highly DNA-damage-sensitive DDP-overproducing strains. DDP-overproducing strains with robust sensitivity to each of the three DNA-damaging agents were tested for sensitivity with overproduction of the DDP (red, “before”), then again after the drug treatment, with the DDP-gene repressed (blue, “after”) to determine whether their sensitivity resulted from induction of mutations in genes needed for resistance. After drug treatment, three colonies were recovered on plates with no DDP-gene inducing IPTG, and then tested for DNA-damaging-agent sensitivity again. In all cases, the recovered colonies not induced for DDP overproduction showed no sensitivity to DNA-damaging agents (blue, after). Thus, their DNA-damage sensitivities did not result from heritable mutations.

**Figure S6.**
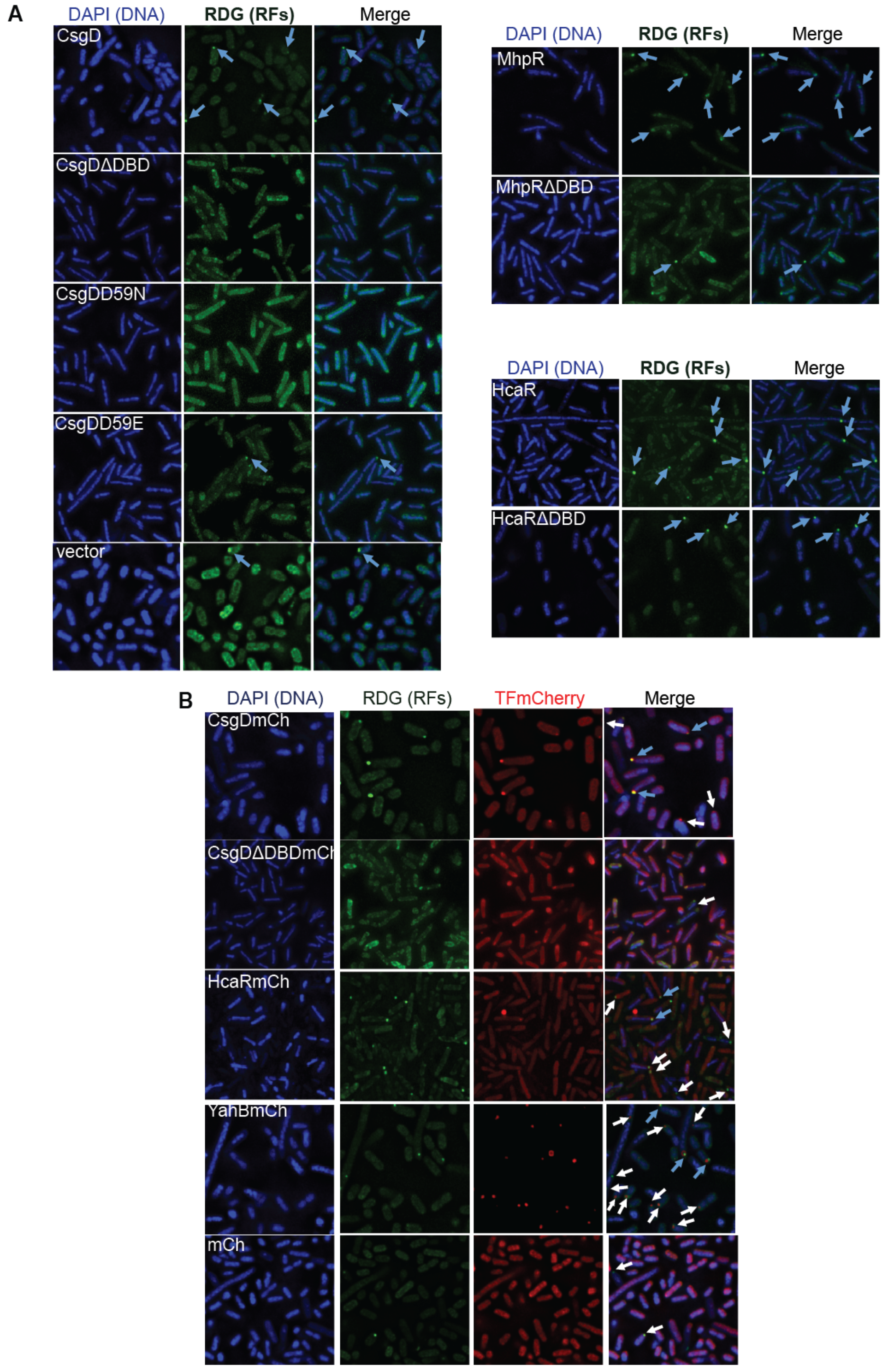
Examples of DNA-binding Transcription Factor Induction of and Co-localization with Replication-Stall (RDG) Foci. (A) DNA-binding ability of DNA-binding transcription factors is required for their promotion of increased RDG (reversed-fork) foci upon overproduction. ΔDBD, in-frame deletion of the DNA-binding domain; CsgDD59N, D59E: single amino-acid changes that reduce CsgD DNA binding (Ogasawara et al., 2011). Three wild-type transcription factors and the corresponding mutants with reduced DNA-binding ability were overproduced in Δ*recA* cells, RDG (reversed-fork) foci quantified (Figure 4C; Table S1), and representative images are shown here. (B) Foci of transcription factors CsgD-mCherry and HcaR-mCherry co-localize with RDG (reversed-fork) foci. Representative examples. Most of the CsgD-mCherry and HcaR-mCherry foci were co-localized with RDG (reversed-fork) foci. Showing weak co-localization, about 5% of the YahB-mCherry transcription-factor foci were co-localized with RDG (reversed-fork) foci. By contrast, mCherry alone rarely forms foci.

**Figure S7.**
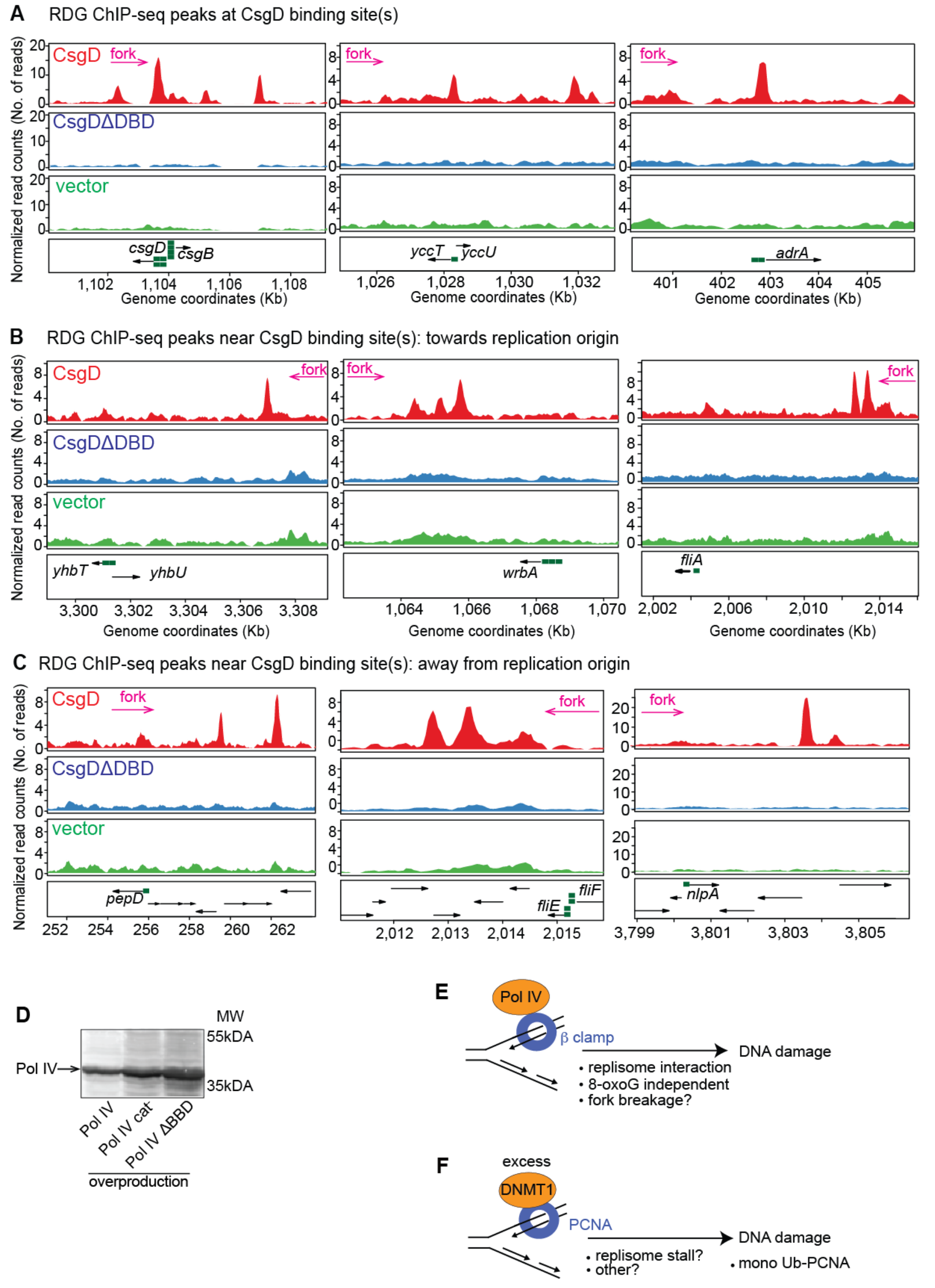
RDG ChIP-seq Detects Stalled-Fork Enrichment Near CsgD-Binding Sites and Controls for Pol IV. Nine of the 10 known, validated CsgD-binding sites (Brombacher et al., 2003; Dudin et al., 2014; Keseler et al., 2017; Ogasawara et al., 2011) showed a CsgD-DNA-binding-domain (DBD)-dependent RDG peak nearby: within 10kb (median 2.8kb), which differs from random simulations of the genomic distribution of the total number of CsgD-DBD-dependent RDG peaks (*p* = 0.01 two-tailed z test, see Supplemental Discussion 12). Isogenic cells overproducing—CsgD, red; CsgD ΔDBD (deletion of the DNA-binding domain), blue; vector only green. Green boxes, CsgD binding site(s). (A) CsgD-DBD-dependent RDG ChIP-seq peaks overlap with three known CsgD-binding sites. (B) CsgD-DBD-dependent RDG ChIP-seq peaks near three known CsgD-binding sites, upstream in the replication path. Increased negative supercoiling between the oncoming fork and the CsgD-bound site may stall replication and causes fork reversal, illustrated Figure 5J. (C) CsgD-DBD-dependent RDG ChIP-seq peaks located near three known CsgD binding sites, downstream in the replication path. Because the upstream peaks (B) are more significantly associated with the known CsgD-binding sites relative to simulation of random genomic distributions (Supplemental Discussion 12), it is possible that the downstream CsgD-DBD-dependent RDG peaks could result from either CsgD binding to sites not yet known, or from indirect consequences of CsgD DNA binding, such as effects of CsgD-regulated gene products interacting with other sites in DNA (Supplemental Discussion 12). Alternatively, some of the downstream CsgD-DBD-dependent RDG peaks could be a direct result of CsgD binding its known site, slowing replication, then encountering an otherwise surmountable obstacle downstream, per Figure 5J. (D) The Pol IV R49F catalytic-mutant (Pol IV cat^−^) and ABBD-mutant proteins do not display reduced protein levels in western blots, in agreement with previous studies (Uchida et al., 2008; Wagner et al., 1999). (E) Model: Pol IV overproduction induces DNA damage by binding the beta replisome sliding clamp. Excess Pol IV interaction with the replisome could potentially slow the replisome causing fork breakage or collapse, or displace DNA-repair proteins that interact with the beta clamp, or otherwise promote DNA damage. (F) Model and hypotheses: overproduced DNMT1 provokes DNA damage independently of its DNA-methylase activity but dependently on binding PCNA, the mammalian replisome sliding clamp and structural homolog of *E. coli* beta. Excess DNMT1 might promote DNA damage by stalling DNA replication, interfering with PCNA-coordinated DNA-repair or translesion-synthesis processes, or by other PCNA-dependent means.

## REFERENCES

Alvaro, D., Lisby, M., and Rothstein, R. (2007). Genome-wide analysis of Rad52 foci reveals diverse mechanisms impacting recombination. PLoS Genet 3, e228.

Aravind, L., Walker, D.R., and Koonin, E.V. (1999). Conserved domains in DNA repair proteins and evolution of repair systems. Nucleic Acids Res 27, 1223–1242.

Asad, N.R., Asad, L.M.B.O., Almeida, C.E.B.d., Felzenszwalb, I., Cabral-Neto, J.B., and Leitão, A.C. (2004). Several pathways of hydrogen peroxide action that damage the E. coli genome. Genetics and Molecular Biology 27, 291–303.

Benjamini, Y., and Hochberg, Y. (1995). Controlling the false discovery rate: a practical and powerful approach to multiple testing. Journal of the royal statistical society Series B (Methodological), 289–300.

Berkopec, A. (2007). HyperQuick algorithm for discrete hypergeometric distribution. Journal of Discrete Algorithms 5, 341–347.

Biniszkiewicz, D., Gribnau, J., Ramsahoye, B., Gaudet, F., Eggan, K., Humpherys, D., Mastrangelo, M.A., Jun, Z., Walter, J., and Jaenisch, R. (2002). Dnmt1 overexpression causes genomic hypermethylation, loss of imprinting, and embryonic lethality. Mol Cell Biol 22, 2124–2135.

Bolger, A.M., Lohse, M., and Usadel, B. (2014). Trimmomatic: a flexible trimmer for Illumina sequence data. Bioinformatics 30, 2114–2120.

Bonocora, R.P., and Wade, J.T. (2015). ChIP-seq for genome-scale analysis of bacterial DNA-binding proteins. Methods Mol Biol 1276, 327–340.

Boratyn, G.M., Schaffer, A.A., Agarwala, R., Altschul, S.F., Lipman, D.J., and Madden, T.L. (2012). Domain enhanced lookup time accelerated BLAST. Biol Direct 7, 12.

Brombacher, E., Dorel, C., Zehnder, A.J., and Landini, P. (2003). The curli biosynthesis regulator CsgD co-ordinates the expression of both positive and negative determinants for biofilm formation in Escherichia coli. Microbiology 149, 2847–2857.

Cameron, K.S., Buchner, V., and Tchounwou, P.B. (2011). Exploring the molecular mechanisms of nickel-induced genotoxicity and carcinogenicity: a literature review. Rev Environ Health 26, 81–92.

Carrasco, B., Cozar, M.C., Lurz, R., Alonso, J.C., and Ayora, S. (2004). Genetic recombination in Bacillus subtilis 168: contribution of Holliday junction processing functions in chromosome segregation. J Bacteriol 186, 5557–5566.

Chatterjee, N., and Walker, G.C. (2017). Mechanisms of DNA damage, repair, and mutagenesis. Environ Mol Mutagen 58, 235–263.

D’Antonio, M., and Ciccarelli, F.D. (2013). Integrated analysis of recurrent properties of cancer genes to identify novel drivers. Genome Biol 14, R52.

Dohrmann, P.R., Correa, R., Frisch, R.L., Rosenberg, S.M., and McHenry, C.S. (2016). The DNA polymerase III holoenzyme contains gamma and is not a trimeric polymerase. Nucleic Acids Res 44, 1285–1297.

Dudin, O., Geiselmann, J., Ogasawara, H., Ishihama, A., and Lacour, S. (2014). Repression of flagellar genes in exponential phase by CsgD and CpxR, two crucial modulators of Escherichia coli biofilm formation. J Bacteriol 196, 707–715.

Elowitz, M.B., Levine, A.J., Siggia, E.D., and Swain, P.S. (2002). Stochastic gene expression in a single cell. Science 297, 1183–1186.

Fitzgerald, D.M., Hastings, P., and Rosenberg, S.M. (2017). Stress-induced mutagenesis: implications in cancer and drug resistance. Annu Rev Cancer Biol 1, 119–140.

Forbes, S.A., Beare, D., Gunasekaran, P., Leung, K., Bindal, N., Boutselakis, H., Ding, M., Bamford, S., Cole, C., Ward, S., et al. (2015). COSMIC: exploring the world’s knowledge of somatic mutations in human cancer. Nucleic Acids Res 43, D805–811.

Foster, P.L. (2006). Methods for determining spontaneous mutation rates. Methods Enzymol 409, 195–213.

Foti, J.J., Devadoss, B., Winkler, J.A., Collins, J.J., and Walker, G.C. (2012). Oxidation of the guanine nucleotide pool underlies cell death by bactericidal antibiotics. Science 336, 315–319.

Friedberg, E.C., Walker, G.C., Siede, W., and Wood, R.D. (2005). DNA repair and mutagenesis (American Society for Microbiology Press).

Frisch, R.L., Su, Y., Thornton, P.C., Gibson, J.L., Rosenberg, S.M., and Hastings, P.J. (2010). Separate DNA Pol II- and Pol IV-dependent pathways of stress-induced mutation during double-strand-break repair in Escherichia coli are controlled by RpoS. J Bacteriol 192, 4694–4700.

Galhardo, R.S., Almeida, C.E., Leitao, A.C., and Cabral-Neto, J.B. (2000). Repair of DNA lesions induced by hydrogen peroxide in the presence of iron chelators in Escherichia coli: participation of endonuclease IV and Fpg. J Bacteriol 182, 1964–1968.

Gao, J., Aksoy, B.A., Dogrusoz, U., Dresdner, G., Gross, B., Sumer, S.O., Sun, Y., Jacobsen, A., Sinha, R., Larsson, E., et al. (2013). Integrative analysis of complex cancer genomics and clinical profiles using the cBioPortal. Sci Signal 6, p11.

Gao, R., Davis, A., McDonald, T.O., Sei, E., Shi, X., Wang, Y., Tsai, P.C., Casasent, A., Waters, J., Zhang, H., et al. (2016). Punctuated copy number evolution and clonal stasis in triple-negative breast cancer. Nat Genet 48, 1119–1130.

Germano, G., Lamba, S., Rospo, G., Barault, L., Magri, A., Maione, F., Russo, M., Crisafulli, G., Bartolini, A., Lerda, G., et al. (2017). Inactivation of DNA repair triggers neoantigen generation and impairs tumour growth. Nature 552, 116–120.

Gutierrez, A., Laureti, L., Crussard, S., Abida, H., Rodriguez-Rojas, A., Blazquez, J., Baharoglu, Z., Mazel, D., Darfeuille, F., Vogel, J., et al. (2013). beta-Lactam antibiotics promote bacterial mutagenesis via an RpoS-mediated reduction in replication fidelity. Nat Commun 4, 1610.

Hanahan, D., and Weinberg, R.A. (2011). Hallmarks of cancer: the next generation. Cell 144, 646–674.

Hanzelmann, S., Castelo, R., and Guinney, J. (2013). GSVA: gene set variation analysis for microarray and RNA-seq data. BMC Bioinformatics 14, 7.

Hartwig, A. (2001). Role of magnesium in genomic stability. Mutat Res 475, 113–121.

Hastings, P.J., Hersh, M.N., Thornton, P.C., Fonville, N.C., Slack, A., Frisch, R.L., Ray, M.P., Harris, R.S., Leal, S.M., and Rosenberg, S.M. (2010). Competition of Escherichia coli DNA polymerases I, II and III with DNA Pol IV in stressed cells. PLoS One 5, e10862.

Hastings, P.J., Lupski, J.R., Rosenberg, S.M., and Ira, G. (2009). Mechanisms of change in gene copy number. Nat Rev Genet 10, 551–564.

Hastings, P.J., Quah, S.K., and von Borstel, R.C. (1976). Spontaneous mutation by mutagenic repair of spontaneous lesions in DNA. Nature 264, 719–722.

Heckman, K.L., and Pease, L.R. (2007). Gene splicing and mutagenesis by PCR-driven overlap extension. Nat Protoc 2, 924–932.

Heltzel, J.M., Maul, R.W., Wolff, D.W., and Sutton, M.D. (2012). Escherichia coli DNA polymerase IV (Pol IV), but not Pol II, dynamically switches with a stalled Pol III* replicase. J Bacteriol 194, 3589–3600.

Hendricks, E.C., Szerlong, H., Hill, T., and Kuempel, P. (2000). Cell division, guillotining of dimer chromosomes and SOS induction in resolution mutants (dif, xerC and xerD) of Escherichia coli. Mol Microbiol 36, 973–981.

Hlavac, V., Brynychova, V., Vaclavikova, R., Ehrlichova, M., Vrana, D., Pecha, V., Trnkova, M., Kodet, R., Mrhalova, M., Kubackova, K., et al. (2014). The role of cytochromes p450 and aldo-keto reductases in prognosis of breast carcinoma patients. Medicine (Baltimore) 93, e255.

Hu, C.W., Kornblau, S.M., Slater, J.H., and Qutub, A.A. (2015). Progeny Clustering: A Method to Identify Biological Phenotypes. Sci Rep 5, 12894.

Hu, C.W., and Qutub, A.A. (2016). progenyClust: an R package for Progeny Clustering. R JOURNAL 8, 328–338.

Jackson, A.L., and Loeb, L.A. (2001). The contribution of endogenous sources of DNA damage to the multiple mutations in cancer. Mutat Res 477, 7–21.

Jackson, S.P., and Bartek, J. (2009). The DNA-damage response in human biology and disease. Nature 461, 1071–1078.

Jin, B., and Robertson, K.D. (2013). DNA methyltransferases, DNA damage repair, and cancer. Adv Exp Med Biol 754, 3–29.

Johnson, S.C. (1967). Hierarchical clustering schemes. Psychometrika 32, 241–254.

Jones, P.A., Issa, J.P., and Baylin, S. (2016). Targeting the cancer epigenome for therapy. Nat Rev Genet 17, 630–641.

Joshi, M.C., Magnan, D., Montminy, T.P., Lies, M., Stepankiw, N., and Bates, D. (2013). Regulation of sister chromosome cohesion by the replication fork tracking protein SeqA. PLoS Genet 9, e1003673.

Kehres, D.G., and Maguire, M.E. (2002). Structure, properties and regulation of magnesium transport proteins. Biometals 15, 261–270.

Kelley, E.E., Khoo, N.K., Hundley, N.J., Malik, U.Z., Freeman, B.A., and Tarpey, M.M. (2010). Hydrogen peroxide is the major oxidant product of xanthine oxidase. Free Radic Biol Med 48, 493–498.

Keren, I., Wu, Y., Inocencio, J., Mulcahy, L.R., and Lewis, K. (2013). Killing by bactericidal antibiotics does not depend on reactive oxygen species. Science 339, 1213–1216.

Keseler, I.M., Mackie, A., Santos-Zavaleta, A., Billington, R., Bonavides-Martinez, C., Caspi, R., Fulcher, C., Gama-Castro, S., Kothari, A., Krummenacker, M., et al. (2017). The EcoCyc database: reflecting new knowledge about Escherichia coli K-12. Nucleic Acids Res 45, D543–D550.

Kim, S.R., Matsui, K., Yamada, M., Gruz, P., and Nohmi, T. (2001). Roles of chromosomal and episomal dinB genes encoding DNA pol IV in targeted and untargeted mutagenesis in Escherichia coli. Mol Genet Genomics 266, 207–215.

Kinner, A., Wu, W., Staudt, C., and Iliakis, G. (2008). Gamma-H2AX in recognition and signaling of DNA double-strand breaks in the context of chromatin. Nucleic Acids Res 36, 5678–5694.

Kinzler, K.W., and Vogelstein, B. (1997). Cancer-susceptibility genes. Gatekeepers and caretakers. Nature 386, 761, 763.

Kobayashi, S., Valentine, M.R., Pham, P., O’Donnell, M., and Goodman, M.F. (2002). Fidelity of Escherichia coli DNA polymerase IV. Preferential generation of small deletion mutations by dNTP-stabilized misalignment. J Biol Chem 277, 34198–34207.

Kuzminov, A. (1995). Collapse and repair of replication forks in Escherichia coli. Mol Microbiol 16, 373–384.

Kuzminov, A. (2011). Homologous Recombination-Experimental Systems, Analysis, and Significance. EcoSal Plus 4.

Levine, M., and Tjian, R. (2003). Transcription regulation and animal diversity. Nature 424, 147–151.

Li, H. (2013). Aligning sequence reads, clone sequences and assembly contigs with BWA-MEM. arXiv preprint arXiv:13033997.

Loiselle, F.B., and Casey, J.R. (2010). Measurement of Intracellular pH. Methods Mol Biol 637, 311–331.

Lovejoy, C.A., Xu, X., Bansbach, C.E., Glick, G.G., Zhao, R., Ye, F., Sirbu, B.M., Titus, L.C., Shyr, Y., and Cortez, D. (2009). Functional genomic screens identify CINP as a genome maintenance protein. Proc Natl Acad Sci U S A 106, 19304–19309.

Magnan, D., Joshi, M.C., Barker, A.K., Visser, B.J., and Bates, D. (2015). DNA Replication Initiation Is Blocked by a Distant Chromosome-Membrane Attachment. Curr Biol 25, 2143–2149.

Makarova, K.S., and Koonin, E.V. (2013). Archaeology of eukaryotic DNA replication. Cold Spring Harb Perspect Biol 5, a012963.

Maor-Shoshani, A., Reuven, N.B., Tomer, G., and Livneh, Z. (2000). Highly mutagenic replication by DNA polymerase V (UmuC) provides a mechanistic basis for SOS untargeted mutagenesis. Proc Natl Acad Sci U S A 97, 565–570.

McClure, R., Balasubramanian, D., Sun, Y., Bobrovskyy, M., Sumby, P., Genco, C.A., Vanderpool, C.K., and Tjaden, B. (2013). Computational analysis of bacterial RNA-Seq data. Nucleic Acids Res 41, e140.

McPartland, A., Green, L., and Echols, H. (1980). Control of recA gene RNA in E. coli: regulatory and signal genes. Cell 20, 731–737.

Merrikh, H., Zhang, Y., Grossman, A.D., and Wang, J.D. (2012). Replication-transcription conflicts in bacteria. Nat Rev Microbiol 10, 449–458.

Michaels, M.L., Pham, L., Cruz, C., and Miller, J.H. (1991). MutM, a protein that prevents G.C----T.A transversions, is formamidopyrimidine-DNA glycosylase. Nucleic Acids Res 19, 3629–3632.

Miller, J. (1993). A short course in bacterial genetics: a laboratory manual and handbook for Escherichia coli and related bacteria. Trends in Biochemical Sciences-Library Compendium 18, 193.

Mortusewicz, O., Schermelleh, L., Walter, J., Cardoso, M.C., and Leonhardt, H. (2005). Recruitment of DNA methyltransferase I to DNA repair sites. Proc Natl Acad Sci U S A 102, 8905–8909.

Nehring, R.B., Gu, F., Lin, H.Y., Gibson, J.L., Blythe, M.J., Wilson, R., Bravo Nunez, M.A., Hastings, P.J., Louis, E.J., Frisch, R.L., et al. (2016). An ultra-dense library resource for rapid deconvolution of mutations that cause phenotypes in Escherichia coli. Nucleic Acids Res 44, e41.

Ogasawara, H., Yamamoto, K., and Ishihama, A. (2011). Role of the biofilm master regulator CsgD in cross-regulation between biofilm formation and flagellar synthesis. J Bacteriol 193, 2587–2597.

Paulsen, R.D., Soni, D.V., Wollman, R., Hahn, A.T., Yee, M.C., Guan, A., Hesley, J.A., Miller, S.C., Cromwell, E.F., Solow-Cordero, D.E., et al. (2009). A genome-wide siRNA screen reveals diverse cellular processes and pathways that mediate genome stability. Mol Cell 35, 228–239.

Pennington, J.M., and Rosenberg, S.M. (2007). Spontaneous DNA breakage in single living Escherichia coli cells. Nat Genet 39, 797–802.

Pickering, M.T., and Kowalik, T.F. (2006). Rb inactivation leads to E2F1-mediated DNA double-strand break accumulation. Oncogene 25, 746–755.

Putnam, C.D., Srivatsan, A., Nene, R.V., Martinez, S.L., Clotfelter, S.P., Bell, S.N., Somach, S.B., de Souza, J.E., Fonseca, A.F., de Souza, S.J., et al. (2016). A genetic network that suppresses genome rearrangements in Saccharomyces cerevisiae and contains defects in cancers. Nat Commun 7, 11256.

Rahman, M., Jackson, L.K., Johnson, W.E., Li, D.Y., Bild, A.H., and Piccolo, S.R. (2015). Alternative preprocessing of RNA-Sequencing data in The Cancer Genome Atlas leads to improved analysis results. Bioinformatics 31, 3666–3672.

Ramirez, F., Dundar, F., Diehl, S., Gruning, B.A., and Manke, T. (2014). deepTools: a flexible platform for exploring deep-sequencing data. Nucleic Acids Res 42, W187–191.

Reams, A.B., Kofoid, E., Savageau, M., and Roth, J.R. (2010). Duplication frequency in a population of Salmonella enterica rapidly approaches steady state with or without recombination. Genetics 184, 1077–1094.

Renzette, N., Gumlaw, N., Nordman, J.T., Krieger, M., Yeh, S.P., Long, E., Centore, R., Boonsombat, R., and Sandler, S.J. (2005). Localization of RecA in Escherichia col K-12 using RecA-GFP. Mol Microbiol 57, 1074–1085.

Saito, Y., Uraki, F., Nakajima, S., Asaeda, A., Ono, K., Kubo, K., and Yamamoto, K. (1997). Characterization of endonuclease III (nth) and endonuclease VIII (nei) mutants of Escherichia coli K-12. J Bacteriol 179, 3783–3785.

Saka, K., Tadenuma, M., Nakade, S., Tanaka, N., Sugawara, H., Nishikawa, K., Ichiyoshi, N., Kitagawa, M., Mori, H., Ogasawara, N., et al. (2005). A complete set of Escherichia coli open reading frames in mobile plasmids facilitating genetic studies. DNA Res 12, 63–68.

Sakaguchi, K., Herrera, J.E., Saito, S., Miki, T., Bustin, M., Vassilev, A., Anderson, C.W., and Appella, E. (1998). DNA damage activates p53 through a phosphorylation-acetylation cascade. Genes Dev 12, 2831–2841.

Schmidt, K., Wolfe, D.M., Stiller, B., and Pearce, D.A. (2009). Cd2+, Mn2+, Ni2+ and Se2+ toxicity to Saccharomyces cerevisiae lacking YPK9p the orthologue of human ATP13A2. Biochem Biophys Res Commun 383, 198–202.

Seigneur, M., Bidnenko, V., Ehrlich, S.D., and Michel, B. (1998). RuvAB acts at arrested replication forks. Cell 95, 419–430.

Serres, M.H., Gopal, S., Nahum, L.A., Liang, P., Gaasterland, T., and Riley, M. (2001). A functional update of the Escherichia coli K-12 genome. Genome Biol 2, RESEARCH0035.

Shee, C., Cox, B.D., Gu, F., Luengas, E.M., Joshi, M.C., Chiu, L.Y., Magnan, D., Halliday, J.A., Frisch, R.L., Gibson, J.L., et al. (2013). Engineered proteins detect spontaneous DNA breakage in human and bacterial cells. Elife 2, e01222.

Sottoriva, A., Kang, H., Ma, Z., Graham, T.A., Salomon, M.P., Zhao, J., Marjoram, P., Siegmund, K., Press, M.F., Shibata, D., et al. (2015). A Big Bang model of human colorectal tumor growth. Nat Genet 47, 209–216.

Stanton, R.C. (2012). Glucose-6-phosphate dehydrogenase, NADPH, and cell survival. IUBMB Life 64, 362–369.

Steighner, R.J., and Povirk, L.F. (1990). Bleomycin-induced DNA lesions at mutational hot spots: implications for the mechanism of double-strand cleavage. Proc Natl Acad Sci U S A 87, 8350–8354.

Stern, M.J., Ames, G.F., Smith, N.H., Robinson, E.C., and Higgins, C.F. (1984). Repetitive extragenic palindromic sequences: a major component of the bacterial genome. Cell 37, 1015–1026.

Stratton, M.R. (2011). Exploring the genomes of cancer cells: progress and promise. Science 331, 1553–1558.

Sun, G., Chung, D., Liang, K., and Keles, S. (2013). Statistical analysis of ChIP-seq data with MOSAiCS. Methods Mol Biol 1038, 193–212.

Szklarczyk, D., Franceschini, A., Wyder, S., Forslund, K., Heller, D., Huerta-Cepas, J., Simonovic, M., Roth, A., Santos, A., Tsafou, K.P., et al. (2015). STRING v10: protein-protein interaction networks, integrated over the tree of life. Nucleic Acids Res 43, D447–452.

Tehranchi, A.K., Blankschien, M.D., Zhang, Y., Halliday, J.A., Srivatsan, A., Peng, J., Herman, C., and Wang, J.D. (2010). The transcription factor DksA prevents conflicts between DNA replication and transcription machinery. Cell 141, 595–605.

Thomason, L.C., Costantino, N., and Court, D.L. (2007). E. coli genome manipulation by P1 transduction. Curr Protoc Mol Biol Chapter 1, Unit 1 17.

Tipparaju, S.M., Liu, S.Q., Barski, O.A., and Bhatnagar, A. (2007). NADPH binding to beta-subunit regulates inactivation of voltage-gated K(+) channels. Biochem Biophys Res Commun 359, 269–276.

Torkelson, J., Harris, R.S., Lombardo, M.J., Nagendran, J., Thulin, C., and Rosenberg, S.M. (1997). Genome-wide hypermutation in a subpopulation of stationary-phase cells underlies recombination-dependent adaptive mutation. EMBO J 16, 3303–3311.

Tubbs, A., and Nussenzweig, A. (2017). Endogenous DNA Damage as a Source of Genomic Instability in Cancer. Cell 168, 644–656.

Uchida, K., Furukohri, A., Shinozaki, Y., Mori, T., Ogawara, D., Kanaya, S., Nohmi, T., Maki, H., and Akiyama, M. (2008). Overproduction of Escherichia coli DNA polymerase DinB (Pol IV) inhibits replication fork progression and is lethal. Mol Microbiol 70, 608–622.

Vafa, O., Wade, M., Kern, S., Beeche, M., Pandita, T.K., Hampton, G.M., and Wahl, G.M. (2002). c-Myc can induce DNA damage, increase reactive oxygen species, and mitigate p53 function: a mechanism for oncogene-induced genetic instability. Mol Cell 9, 1031–1044.

Wagner, J., Fujii, S., Gruz, P., Nohmi, T., and Fuchs, R.P. (2000). The beta clamp targets DNA polymerase IV to DNA and strongly increases its processivity. EMBO Rep 1, 484–488.

Wagner, J., Gruz, P., Kim, S.R., Yamada, M., Matsui, K., Fuchs, R.P., and Nohmi, T. (1999). The dinB gene encodes a novel E. coli DNA polymerase, DNA pol IV, involved in mutagenesis. Mol Cell 4, 281–286.

Wagner, J., and Nohmi, T. (2000). Escherichia coli DNA polymerase IV mutator activity: genetic requirements and mutational specificity. J Bacteriol 182, 4587–4595.

Wahba, L., Costantino, L., Tan, F.J., Zimmer, A., and Koshland, D. (2016). S1-DRIP-seq identifies high expression and polyA tracts as major contributors to R-loop formation. Genes Dev 30, 1327–1338.

Xia, J., Chen, L.T., Mei, Q., Ma, C.H., Halliday, J.A., Lin, H.Y., Magnan, D., Pribis, J.P., Fitzgerald, D.M., Hamilton, H.M., et al. (2016). Holliday junction trap shows how cells use recombination and a junction-guardian role of RecQ helicase. Sci Adv 2, e1601605.

Yang, X., Boehm, J.S., Yang, X., Salehi-Ashtiani, K., Hao, T., Shen, Y., Lubonja, R., Thomas, S.R., Alkan, O., Bhimdi, T., et al. (2011). A public genome-scale lentiviral expression library of human ORFs. Nat Methods 8, 659–661.

Yeeles, J.T., Poli, J., Marians, K.J., and Pasero, P. (2013). Rescuing stalled or damaged replication forks. Cold Spring Harb Perspect Biol 5, a012815.

Yuen, K.W., Warren, C.D., Chen, O., Kwok, T., Hieter, P., and Spencer, F.A. (2007). Systematic genome instability screens in yeast and their potential relevance to cancer. Proc Natl Acad Sci USA 104, 3925–3930.

Zack, T.I., Schumacher, S.E., Carter, S.L., Cherniack, A.D., Saksena, G., Tabak, B., Lawrence, M.S., Zhsng, C.Z., Wala, J., Mermel, C.H., et al. (2013). Pan-cancer patterns of somatic copy number alteration. Nat Genet 45, 1134–1140.

